# Rapid Determination of Drug-to-Antibody Ratios in Antibody–Drug Conjugates Using Ultrafast Microdroplet Digestion Technology

**DOI:** 10.64898/2026.06.02.729562

**Authors:** Yongqing Yang, Juana Perez Sancheza, Timothy Yaroshuk, Md Tanim-Al Hassan, Praneeth Ivan Joel FNU, Thomas Walker, Jim Lau, Mike Knierman, Hui Zhao, Xi Qiu, Karen Luo, Harsha P. Gunawardena, Munkhtsetseg Baatar, Hao Chen

## Abstract

Accurate determination of drug-to-antibody ratios (DARs) is essential for the development, quality control, and performance evaluation of antibody–drug conjugates (ADCs); yet conventional analytical approaches often require extensive sample preparation, long analysis time, and substantial sample consumption. The peak distribution of intact ADCs is highly complex due to inherent glycosylation heterogeneity and variable drug conjugation. By applying enzymatic digestion, ADC can be converted into smaller subunits or deglycosylated species, thereby significantly simplifying the mass spectral profile. This reduction in structural heterogeneity facilitates clearer peak assignment and enables more accurate and reliable DAR quantification. Herein, we report an ultrafast microdroplet digestion–mass spectrometry strategy for rapid DAR characterization of ADCs. Microdroplet enzymatic digestion of antibodies and ADCs occurs within microsecond-time scales during spray ionization, enabling direct online subunit analysis with minimal sample preparation. The method was validated using NIST monoclonal antibody (mAb) conjugated to ADC mimics spanning low to high DAR (0-14) ranges, Cetuximab-derived ADC mimics (DAR∼5) with complex glycosylation, and the commercial ADC Kadcyla (DAR∼3.5). Consistent DAR values were obtained across multiple enzymatic workflows (IdeS, EndoS2, and EndoF3) with good reproducibility (%CV typically <5%). This approach substantially reduces analysis time while maintaining analytical accuracy and structural specificity, providing a rapid, sensitive platform for high-throughput ADC characterization and process monitoring.

**Graphic Abstract:** 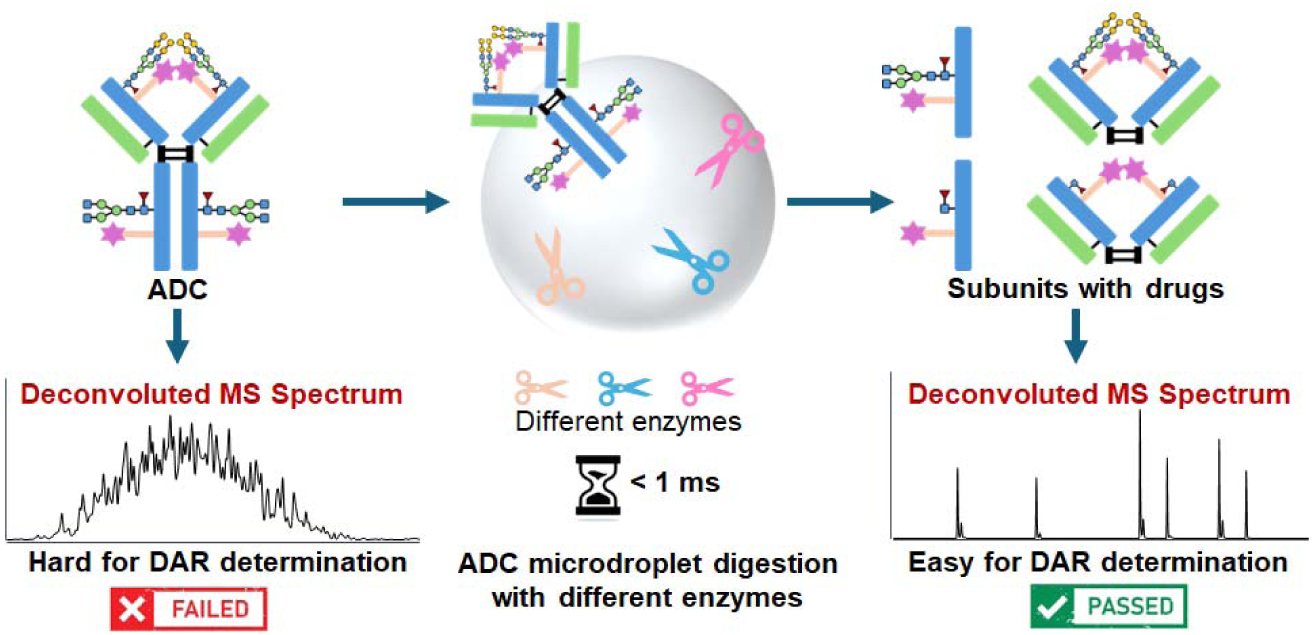

## Introduction

Antibody–drug conjugates (ADCs) represent one of the most transformative therapeutic modalities in modern medicine because they uniquely combine the high target specificity of monoclonal antibodies with the potent cytotoxicity of small-molecule drugs.^1–3^ This targeted delivery strategy enables selective destruction of diseased cells while minimizing systemic toxicity, overcoming limitations of conventional chemotherapy. As a result, ADCs are reshaping precision oncology and expanding treatment options for patients with previously difficult-to-treat cancers.^4,5^ Despite these therapeutic successes, the inherent structural complexity and molecular heterogeneity of ADCs present rigorous analytical exigencies.^6,7^ These challenges span the entire developmental pipeline, from initial bioconjugation and manufacturing to stringent quality control protocols, necessitating advanced characterization techniques to ensure product consistency and safety. One of the most critical quality attributes of ADCs is the drug-to-antibody ratio (DAR), defined as the average number of drug molecules conjugated to each antibody. DAR strongly influences the pharmacokinetics, efficacy, stability, and toxicity of ADCs.^8,9^ ADCs with excessively high DAR values often exhibit increased aggregation and rapid clearance,^10^ whereas those with low DAR values may suffer from insufficient therapeutic potency.^9,11^ Moreover, ADC preparations typically consist of a distribution of species with different DAR values rather than a single homogeneous population,^12^ making accurate determination of both DAR and DAR distribution essential for process optimization and batch-to-batch consistency.

A variety of analytical techniques have been developed to characterize ADC’s DAR, including UV/Vis spectroscopy,^13,14^ hydrophobic interaction chromatography (HIC),^15,16^ size-exclusion chromatography (SEC),^17,18^ and capillary electrophoresis (CE)^19^. While these methods are robust and well-established, they often require relatively long analysis time (typically > 30 min) and substantial sample consumption and may provide limited structural information. Mass spectrometry (MS)-based methods, which have already been well-developed for antibody characterization and proteomics analysis,^20–23^ including intact mass analysis,^24^ ion mobility spectrometry coupled to mass spectrometry (i.e., IMS-MS),^25,26^ reduced ADC analysis, and middle-down or subunit-level characterization, offer superior resolution and molecular specificity, enabling direct identification of individual DAR species. Recently, Jennifer and co-workers^27^ developed a software tool termed iFishMass for the quantitative analysis of digested ADC peptides, representing a significant advance in mass spectrometry–based characterization of ADCs. Nevertheless, these MS-based workflows typically involve extensive sample preparation steps, such as buffer exchange, reduction, alkylation and enzymatic digestion, which increase analysis time and limit throughput. As ADC pipelines continue to expand, there is a growing demand for rapid, low-consumption, and high-throughput analytical methods capable of providing timely DAR information during conjugation optimization, formulation screening, and process development. Ideally, such methods should be compatible with direct MS detection.

Microdroplet-based techniques have recently emerged as powerful tools for accelerating chemical and biochemical reactions.^28–43^ Microdroplets, typically generated as micrometer-scale droplets, exhibit extremely high surface-area-to-volume ratios, enhanced mass transfer, and unique interfacial environments that can significantly alter reaction kinetics and analytical performance compared to bulk solutions. Previously, we reported an ultrafast microdroplet-based digestion approach in which antibody light and heavy chains were digested by trypsin within less than 1 ms.^32,44,45^ Ultrafast digestion of intact antibodies using the enzyme FabRICATOR (IdeS), FabALACTICA (IgdE) and PNGase F was also demonstrated.^46,47^ To integrate this rapid digestion protocol into a robust analysis workflow, microdroplet reactions were further coupled with automated flow injection (FI) and Agilent Jet Stream (AJS) ion source for ionization,^46,48^ providing online mass spectrometric analysis capability.^46–48^ More recently, we demonstrated efficient and reproducible ultrafast microdroplet digestion (<1 ms) of mAbs bearing Fc mutations (LALA, LAGA, and LFLE) using FabRICATOR Xtra.^48^ However, the application of microdroplet-based enzymatic digestion techniques for ADC characterization with high digestion efficiency, particularly for rapid determination of DAR, remains largely unexplored.^49^

In this study, we report a microdroplet mass spectrometry approach for rapid (< 1 ms) digestion of ADCs, to determine DAR in antibody conjugates and their subunits. The performance of the microdroplet-based DAR analysis was evaluated using ADC mimics of NISTmAb and Cetuximab as well as commercial ADC, Kadcyla. NIST mAb ADC mimics having low-, medium-, and high-levels of NHS-PEG_4_-Biotin conjugates were synthesized and subjected to microdroplet digestion analysis. Microdroplet IdeS digestion generated F(ab’)2 (fragment antigen-binding) and Fc (fragment crystallizable) subunits while GlycINATOR (EndoS2) and endoglycosidase F3 (EndoF3) could rapidly remove the heterogeneous glycans found in the Fc and F(ab’)2 domains in microdroplets. As a result, analysis of the resulting F(ab’)2 and Fc subunits enabled accurate determination of DAR values and detailed assessment of drug distributions across individual subunits. This strategy provides a rapid and reliable platform for ADC characterization and offers new opportunities for accelerating ADC development and quality assessment.

### Experimental Section

Detailed chemical source information, sample desalting, sample synthesis and LC/MS analysis protocols are provided in SI. Briefly, for the synthesis of ADC mimics (Figure 1a), 10 µL of intact antibody stock solution (10 µg/µL) in 100 mM HEPES buffer (pH 8.4) was added with a freshly prepared 10 mM EZ-Link NHS-PEG_4_-Biotin (linker+biotin) stock solution in different antibody-to-payload (i.e., linker+biotin) molar ratios and reaction times with gentle shaking (Figure 1a), to achieve different DARs. NISTmAb and Cetuximab, two antibodies carrying different glycoforms, were chosen as the antibody substrate for synthesizing ADC mimics. Specifically, the following synthetic conditions were used for NISTmAb-based ADCs (Table 1): antibody-to-payload molar ratios of 1:5 (30 min, product: **AM1**), 1:20 (30 min, product: **AM2**), 1:10 (50 min, product: **AM3**), 1:20 (50 min, product: **AM4**), 1:40 (30 min, product: **AM5**), and 1:50 (30 min, product: **AM6**). For Cetuximab-based ADC, an antibody-to-payload molar ratio of 1:10 with a reaction time of 50 min was employed (product: **AM7**). Upon completion of the reaction, excess linker was removed using Zeba desalting columns and the samples were subsequently buffer exchanged using a 10 kDa MWCO centrifugal filter and diluted with deionized water to a final concentration of 0.5 µg/µL.

**Figure 1.**
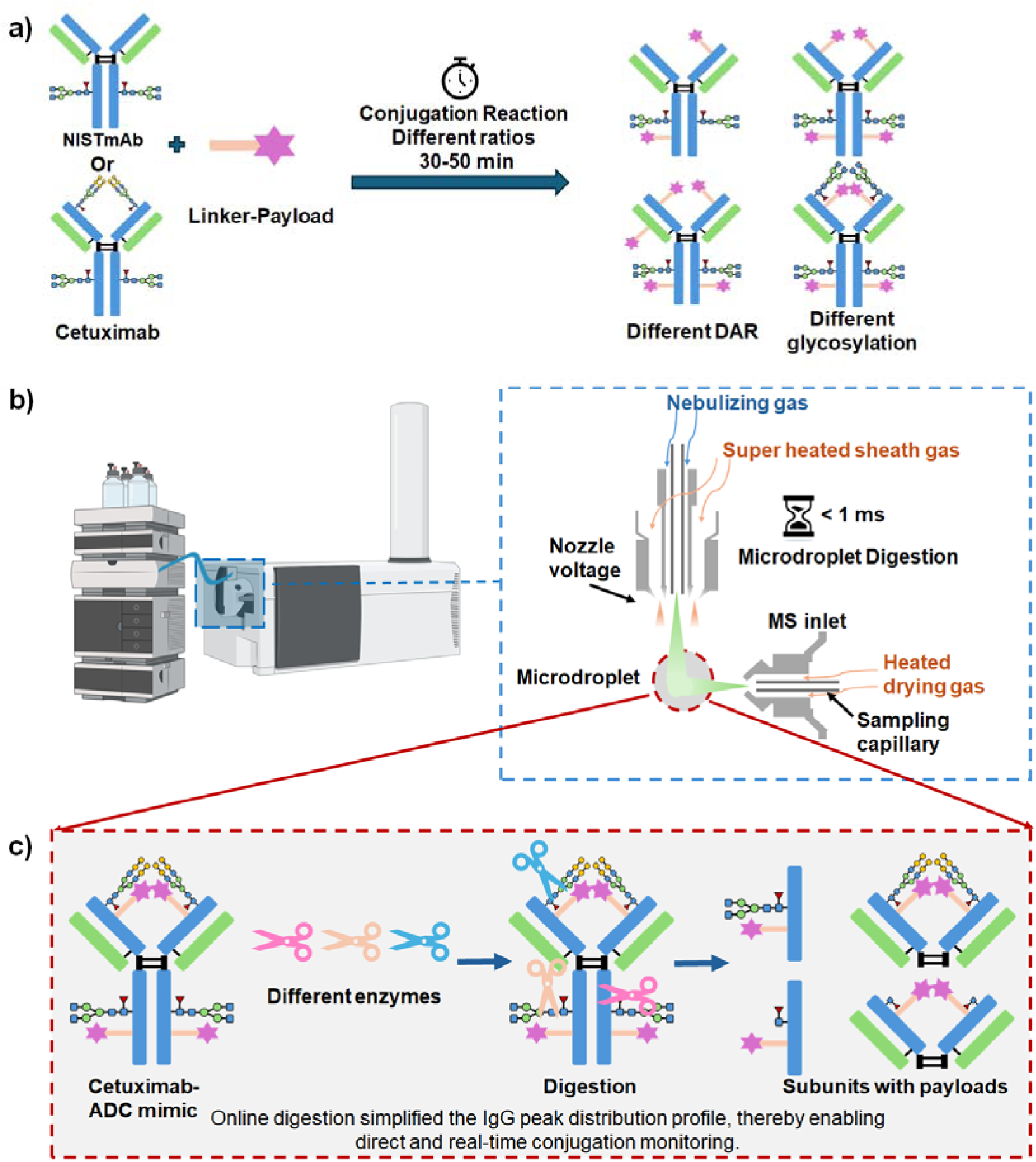
**a)** Scheme showing the conjugation reaction of house-made ADC mimics; b) a general scheme showing performing microdroplet reactions in AJS and c) the subunits formed after digestion with enzymes in microdroplets.

**Table 1.** DAR values of NISTmAb-ADC mimics determined using different enzymes microdroplet digestion.

| Samples | NISTmAb:NHS-PEG<sub>4</sub>-Biotin,<br>reaction time | IdeS | EndoS2 | IdeS + EndoS2 | Average DAR<br>(%CV) |
| --- | --- | --- | --- | --- | --- |
| **AM1** | 1:5, 30 min | 0.45±0.02 | 0.44±0.04 | 0.48±0.01 | 0.46 (4.6%) |
| **AM2** | 1:20, 30 min | 2.71±0.07 | 2.55±0.03 | 2.59±0.10 | 2.62 (3.2%) |
| **AM3** | 1:10, 50 min | 5.30±0.12 | 5.37±0.08 | 5.45±0.08 | 5.37 (1.4%) |
| **AM4** | 1:20, 50 min | 9.56±0.11 | 9.65±0.21 | 9.36±0.12 | 9.52 (1.6%) |
| **AM5** | 1:40, 30 min | 12.11±0.32 | 11.84±0.52 | 12.63±0.20 | 12.19 (3.3%) |
| **AM6** | 1:50, 30 min | 13.51±0.60 | 13.36±0.42 | 13.74±0.55 | 13.54 (1.4%) |
| **Kadcyla** | - | 3.51±0.13 | - | 3.68±0.07 | 3.60 (3.3%) |

### Microdroplet Generation via Flow Injection Analysis (FIA)

Sample injection was performed using an Agilent 1290 Infinity II LC system equipped with a 1290 Infinity II Multisampler and a 1290 Infinity II High-Speed Pump (Agilent Technologies, Santa Clara, CA, USA). The autosampler outlet was directly connected to the AJS source without an analytical column. Samples were introduced into the mass spectrometer in flow injection analysis (FIA) mode. In all experiments, 5 mM ABC was used as the mobile phase.

Microdroplets were generated by spraying an aqueous sample solution containing ADC samples and enzymes (IdeS, EndoS2, or EndoF3) using a preprogrammed autosampler injection sequence. For all experiments, the autosampler sequentially aspirated 1 µL of the ADC solution (0.5 µg/µL) and 1 µL of the enzyme solution (1 unit/µL) from separate vials, followed by thorough mixing within the sampler. The resulting mixture was then injected into the AJS source via the LC system for mass spectrometric analysis. The LC pump flow rate was set to 100 µL/min, and the total FIA run time was 1 min. The column compartment temperature was maintained at 50 °C, while the autosampler temperature was set to 6 °C. The microdroplet formation and the digestion reaction process are illustrated in Figure 1b and 1c. All enzymes used in these experiments were dissolved to 1 unit/µL in deionized water prior to use. For bulk digestion as control, the same ADC-to-enzyme ratio was used.

All digestion experiments in this study were performed at least three times to ensure data reproducibility.

### Mass Spectrometry

Mass spectrometric analysis was performed using a 6545XT Q-TOF mass spectrometer (Agilent Technologies, Santa Clara, CA, USA) equipped with an AJS electrospray ionization source. FIA was conducted in the positive electrospray ionization mode. The following source and ion optics parameters were used for all the experiments: drying gas (N_2_) temperature, 350 °C; drying gas flow rate, 8 L/min; nebulizer pressure, 35 psi; sheath gas temperature, 400 °C; sheath gas flow rate, 8 L/min; nozzle voltage, 500 V; capillary voltage, 3500 V; skimmer voltage, 65 V; octupole radio frequency (RF), 750 V; and fragmentor voltage, 250 V. The time-of-flight (TOF) analyzer was tuned and calibrated in high-mass mode with an upper mass range of 30 kDa. Mass spectra were acquired over an m/z range of 1000–8000 at an acquisition rate of 1 spectrum per second.

### Data analysis

The LC-MS workflow was executed using MassHunter software Ver 11.0 (Agilent Technologies, Santa Clara, CA). Intact mass deconvolution was performed on all MS spectra using the intact MS workflow in the Bioconfirm Software Ver. 12.1 (Agilent Technologies, Santa Clara, CA). Peaks from the deconvoluted mass spectra were automatically assigned to each subunit. For DAR calculations, we first used Bioconfirm to assign the different DAR peaks of antibodies or subunits and then did the calculation based on the peak intensities using eq. 1 and 2 as described below.

## Results and Discussion

In this study, we prepared in-house ADC mimics by conjugating NHS-PEG_4_-Biotin to NISTmAb and Cetuximab (Figure 1a and Figure S1), in which biotin serves as a payload mimic. By varying the antibody-to-payload ratios and reaction times, 6 different NISTmAb-based ADC mimics were generated (**AM1-AM6**, Table 1). It is expected that, with increased reaction time and antibody-to-payload ratio, more payloads would be added to antibody, thus providing an increased DAR value. We tentatively classified **AM1** to **AM6** into 3 groups, low conjugated ADCs (i.e., **AM1** and **AM2**), moderately conjugated ADCs (i.e., **AM3** and **AM4**) and highly conjugated ADCs (i.e., **AM5** and **AM6**). Before we investigated the DAR values of the **AM1**-**AM6** using our microdroplet-digestion approach, intact NISTmAb was examined first for microdroplet digestion as control. Figures S2a and b show the mass spectrum of intact NISTmAb and the corresponding deconvoluted mass spectrum (Figure S2b). The zoomed-in spectrum (Figure S2c) reveals multiple glycoforms of NISTmAb, including G0F/G0F, G0F/G1F, G1F/G1F, G1F/G2F, and G2F/G2F species. IdeS is an NISTmAb-specific cysteine protease that cleaves antibodies at a single site below the hinge region, generating homogeneous F(ab’)2 and Fc-derived fragments (Figure S3a). Following IdeS digestion, two dominant peaks corresponding to scFc (single-chain fragment crystallizable) and F(ab’)2 were observed (Figure S3c). Expanded views of the spectrum reveal the presence of multiple glycoforms (G0F, G1F, and G2F) in the scFc region (Figure S3d), whereas the F(ab’)2 region exhibits a single dominant peak without observable glycoform heterogeneity and a small peak with 162 Da heavier corresponding to glycation (Figure S3e, probably due to the trace amount of glucose present in the purchased NISTmAb which modified the antibody). In addition, another enzyme EndoS2, an IgG-specific endoglycosidase that hydrolyzes all glycoforms present at the Fc N-glycosylation sites (Figure S4a) to generate antibodies bearing a residual N-acetylglucosamine and fucose (GlcNAc+Fuc) moiety on each heavy chain, was used. Following EndoS2 digestion (MS spectrum shown in Figure S4b), a single dominant peak corresponding to the GlcNAc+Fuc/GlcNAc+Fuc-modified NIST (145851.24 Da) was observed without glycoform heterogeneity (Figures S4c and 4d). Simultaneous digestion of the antibody using Ides and EndoS2 was also explored. If both enzymes react simultaneously under microdroplet conditions, scFc and F(ab’)2 fragments are expected to form, with the Fc (Fragment crystallizable)-associated glycans removed (Figure S5a). Indeed, after dual enzyme digestion (MS spectrum shown in Figure S5b), peaks corresponding to scFc and F(ab’)2 were clearly observed (Figure S5c). An expanded view of the scFc region reveals a molecular weight of 24136.91 Da for the GlcNAc+Fuc-modified scFc (Figure S5d), which is lower than those of the G0F, G1F, and G2F glycoforms observed in the IdeS-only digestion (Figure S3c). In contrast, the expanded F(ab’)2 region (Figure S5e) closely resembles that obtained from IdeS digestion alone (Figure S3e), indicating that EndoS2 selectively affects Fc glycosylation without altering the F(ab’)2 subunit. All the digest reactions showed above happens within 1 ms^48^ which is much faster than the traditional digestion methods. Note that, under all three digestion conditions (Figures S3–S5), the signal corresponding to intact NISTmAb was nearly absent, indicating that digestion efficiencies approached 100% under AJS microdroplet conditions.

We then examined the low-conjugation ADC samples, including **AM1** and **AM2**. The corresponding mass spectra of intact **AM1** and **AM2** are shown in Figures S6a and S6b, respectively. After deconvolution, the molecular weights of **AM1** (148677.63 Da; Figure S6c) and **AM2** (149631.26 Da; Figure S6d) were found to be higher than that of intact NISTmAb (148367.31 Da; Figure S2), indicating the successful payload conjugation to NISTmAb. Compared to intact NISTmAb (Figure S2c), a significantly larger number of peaks were observed from zoomed-in view of the **AM1** spectrum (Figure S6e), reflecting contributions from both different glycoforms and multiply added payloads. The high spectral complexity makes peak assignment challenging. In this study, the EZ-Link NHS-PEG_4_-Biotin (MW: 588.67 Da) was used, which selectively reacted with lysine residues. Conjugation of a single NHS-PEG_4_-Biotin resulted in a net mass increase of 473.22 Da on the antibody. It is noteworthy that this mass shift is comparable to the mass differences among different glycoforms. For instance, the mass differences between G0F/G0F and G1F/G2F, as well as between G0F/G1F and G2F/G2F, are both approximately 486 Da. As a result, deconvoluted peaks overlap with each other; for example, the NISTmAb-G1F/G2F glycoform has a molecular weight similar to that of the NISTmAb-G0F/G0F species conjugated with one payload (only has 10 Da difference). This overlap complicates direct DAR determination based solely on mass spectra. The spectral complexity was even more pronounced for the **AM2** sample (Figure S6f). To enable unambiguous assignment of peaks corresponding to different drug loads, in this study, ADC mimics were digested with IdeS to generate subunits and with EndoS2 to remove glycans, reducing the spectral complexity and allowing the DAR of subunits to be clearly analyzed.

The Ides microdroplet digestion mass spectra and deconvoluted spectra of **AM1** and **AM2** are shown in Figures S7a–S7d. As indicated in Figures S7c and S7d, the primary digestion products were scFc and F(ab’)2 subunits, both of which exhibited high signal intensities, enabling reliable subunit-level DAR analysis. Expanded views of the scFc region (Figure S7e) closely matches that of intact NISTmAb digested with IdeS (Figure S3d) and has no evidence of payload conjugation, indicating that DAR[scFc] for **AM1** is zero. In contrast, comparison of the F(ab’)2 region of **AM1** with that of Ides-digested NISTmAb (Figure S3e) reveals the presence of both unconjugated F(ab’)2 and F(ab’)2 species conjugated with one payload (F(ab’)2 + D), as shown in Figure S7g. Using eq. 1, the DAR[F(ab’)2] for **AM1** was calculated to be 0.45. Consequently, the overall DAR for **AM1** was determined to be 0.45 as the sum of the DARs from each subunit (eq. 2).

In eq. 1, DAR is calculated using an approach similar to previous reports,^26,50^ in which the weighted average of the MS peak intensities in the deconvoluted mass spectrum is used.

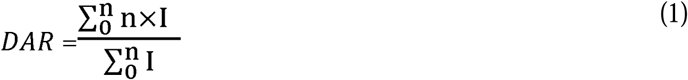

In eq. 2, **n** denotes the number of payload/drug molecules conjugated to a given subunit species, **I_i_**represents the corresponding deconvoluted peak intensity corresponding to that species, and the subscripts scFc and F(ab’)2 indicate the scFc and F(ab’)2 subunits, respectively. The factor of 2 (in front of the scFc term) accounts for the stoichiometry of the antibody structure, as two Fc-related chains contribute to the total DAR calculation.

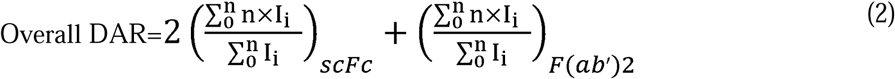

For **AM2**, partial payload conjugation was observed on the scFc subunits (Figure S7f). The DAR values for the G0F and G1F glycoforms were calculated to be 0.44 and 0.43 using eq. 1, respectively, resulting in a DAR[scFc] of 0.44. In addition, the F(ab’)2 subunits exhibited multiple peaks corresponding to species conjugated with 0–4 linker molecules (Figure S7h). Based on eq. 1, the DAR[F(ab’)2] for **AM2** was calculated to be 1.83. Accordingly, the overall DAR for **AM2** determined from IdeS microdroplet digestion using eq. 2 was DAR=2DAR[scFc]+DAR[F(ab’)2]=2.71. A summary of these results is provided in Table 1 and Table S1. The microdroplet digestion efficiencies of these ADC mimics are presented in Table S2. As shown, all ADC mimics subjected to microdroplet digestion in this study exhibited digestion efficiencies above 96% (the digestion efficiency reported in this study was calculated based on the comparison of signal intensities of the G0F/G0F species before and after digestion. Figure S7i is provided as an example to illustrate the calculation), indicating that nearly all ADC mimics were completely digested within 1 ms under microdroplet conditions.

Subsequently, EndoS2 was employed to digest **AM1** and **AM2**. The mass spectra and corresponding deconvoluted spectra of **AM1** and **AM2** after EndoS2 digestion are shown in Figures S8a–S8d. Comparison of Figures S8c–d with Figures S6c–d reveals a mass decrease corresponding to cleavage products of EndoS2 for both **AM1** and **AM2** following microdroplet digestion, confirming efficient removal of G0F/G0F on the Fc domain (digestion efficiency: 97.3% and 97.1% respectively; the reported digestion efficiency was calculated similar to the IdeS digestion case and one calculation example is provided in Figure S8g). An expanded view of the deconvoluted spectrum of EndoS2-digested **AM1** (Figure S8e) shows only three major peaks, corresponding to NISTmAb–GlcNAc+Fuc, NISTmAb–GlcNAc+Fuc+D, and NISTmAb– GlcNAc+Fuc+2D (note that +×D means the conjugation with x number of NHS-PEG_4_-Biotin).

Compared with the spectrum prior to EndoS2 microdroplet digestion (Figure S6e), the spectral complexity was substantially reduced. Using eq. 1, the DAR of **AM1** was determined to be 0.44, which is in excellent agreement with the value obtained from IdeS digestion (0.45; Figure S7). Similarly, the expanded spectrum of EndoS2-digested **AM2** (Figure S8f) contains conjugates with 0–6 NHS-PEG_4_-Biotin, representing a marked simplification relative to the intact **AM2** spectrum prior to EndoS2 microdroplet digestion (Figure S6f). Application of eq. 1 yielded a DAR of 2.55 for **AM2**, which closely matches the value obtained from IdeS digestion (2.71; Figure S7).

When IdeS and EndoS2 were co-sprayed with ADC mimics and allowed to react simultaneously under microdroplet conditions, subunit signals analogous were obtained to enable subunit-level DAR analysis. The mass spectra obtained from microdroplet reactions of **AM1** and **AM2** using combined IdeS and EndoS2 digestion are shown in Figures S9a and S9b, respectively. After deconvolution, clear signals corresponding to the resulting subunits were observed (Figures S9c and S9d). Analysis of the scFc region for **AM1** (Figure S9e) revealed the presence of an scFc–GlcNAc+Fuc+D peak and DAR[scFc] for **AM1** was calculated to be 0.04. In addition, analysis of the F(ab’)2 region (Figure S9g) yielded a DAR[F(ab’)2] of 0.40. Accordingly, the overall DAR for **AM1** determined using the combined microdroplet digestion approach was DAR = 2DAR[scFc] + DAR[F(ab’)2] = 0.48 (Table 1). For **AM2**, similarly, the overall DAR for **AM2** determined using this approach was therefore DAR = 2DAR[scFc] + DAR[F(ab’)2] = 2.59 (Table 1, Figures S9f and S9h). The digestion efficiency of this method is shown in Table S2.

Comparison of the results obtained using the three analytical approaches (Table 1) shows that the overall DAR values determined for **AM1** and **AM2** were 0.46 and 2.68, respectively, with coefficients of variation (%CV) below 5% for both samples. These results indicate excellent agreement among the three methods. Therefore, for low-conjugation samples, all three approaches are capable of reliably determining DAR with strong consistency.

Next, we investigated moderately conjugated NISTmAb-ADC mimic samples, **AM3** and **AM4**. Compared with low-DAR samples, ADCs with higher DAR values exhibit increased molecular heterogeneity and spectral complexity, necessitating more careful analysis. The MS spectra and deconvoluted results of **AM3** and **AM4** shown in Figures S10a-d. Zoomed-in views of the deconvoluted spectra reveal extensive peak splitting and complexity for **AM3** (Figure 2a) and **AM4** (Figure S10e). Following the same analytical strategy used for **AM1** and **AM2**, IdeS microdroplet digestion applied to **AM3** and **AM4** produced subunits with high efficiency (97.9% and 99.3%, respectively, Table S2; MS spectra are shown in Figures S11a, Figures 2b and S12a-b). Zoomed-in analysis of the scFc region for **AM3** (Figure 2c) shows that the DAR values of the G0F and G1F glycoforms are similar (0.58 and 0.56, respectively). For **AM4**, the DAR values of these two glycoforms are also very close (1.27 and 1.04, respectively; Figure S12c), Notably, due to increased payload conjugation, the scFc DAR values for **AM3** and **AM4** are higher than those observed for **AM1** and **AM2**. Analysis of the F(ab’)2 region reveals that **AM3** contains 2–6 NHS-PEG_4_-Biotin molecules (Figure 2d), while **AM4** contains 4–10 NHS-PEG_4_-Biotin molecules (Figure S12d), both substantially higher than those observed for **AM1** and **AM2**. The calculated DAR[F(ab’)2] values for **AM3** and **AM4** were 4.16 and 7.24, respectively. Accordingly, the total DAR values determined from IdeS digestion were 5.30 for **AM3** and 9.56 for **AM4** (Table 1 and Table S1). To further validate these results, EndoS2 digestion was also performed. The results are shown in Figures S11b and 2e-f (**AM3**) and S13a-c (**AM4**) and discussed in SI. Subsequently, **AM3** and **AM4** were digested using a combination of IdeS and EndoS2 under microdroplet conditions. The resulting mass spectra are shown in Figures S11c (**AM3**) and S14a (**AM4**), and the deconvoluted subunit spectra are shown in Figure 2g (**AM3**) and Figure S14b (**AM4**), respectively. In contrast to IdeS-only digestion (Figure 2b and Figure S12b), the F(ab’)2 signals were markedly weaker relative to scFc. This effect probably arises from two factors: removal of Fc glycans by EndoS2 increases scFc ionization efficiency, while extensive NHS-PEG_4_-Biotin conjugation on F(ab’)2 suppresses its ionization efficiency. We observed F(ab’)2 and its conjugates by filtering the *m/z* regions of the F(ab’)2 spectrum. As a result, local spectral expansion was required to clearly observe F(ab’)2 signals. Analysis of the expanded scFc region revealed only GlcNAc+Fuc glycoform on scFc subunit conjugated with NHS-PEG_4_-Biotin, yielding DAR[scFc] values of 0.64 for **AM3** (Figure 2h) and 1.10 for **AM4** (Figure S14c), consistent with results obtained using IdeS alone (Table S1). Importantly, removal of glycoform heterogeneity improves the accuracy of scFc-based DAR determination. Despite the reduced signal intensity, expanded views of the F(ab’)2 region (Figures 2i and S14d) clearly reveal the DAR distributions, yielding DAR[F(ab’)2] values of 4.17 for **AM3** and 7.16 for **AM4**, again in close agreement with previous results (Table S1). Accordingly, the overall DAR values determined using the combined digestion approach were 5.45 for **AM3** and 9.36 for **AM4**, consistent with values obtained using IdeS-only and EndoS2-only digestion methods (Table 1). These results demonstrate that all three analytical strategies are applicable to moderately conjugated NISTmAb–ADC mimic samples (3 < DAR < 10) and yield accurate and consistent DAR values, which is significant as most commercial ADCs with optimal antitumor activity in vivo have DAR values in the range of 3–10.^51,52^

**Figure 2.**
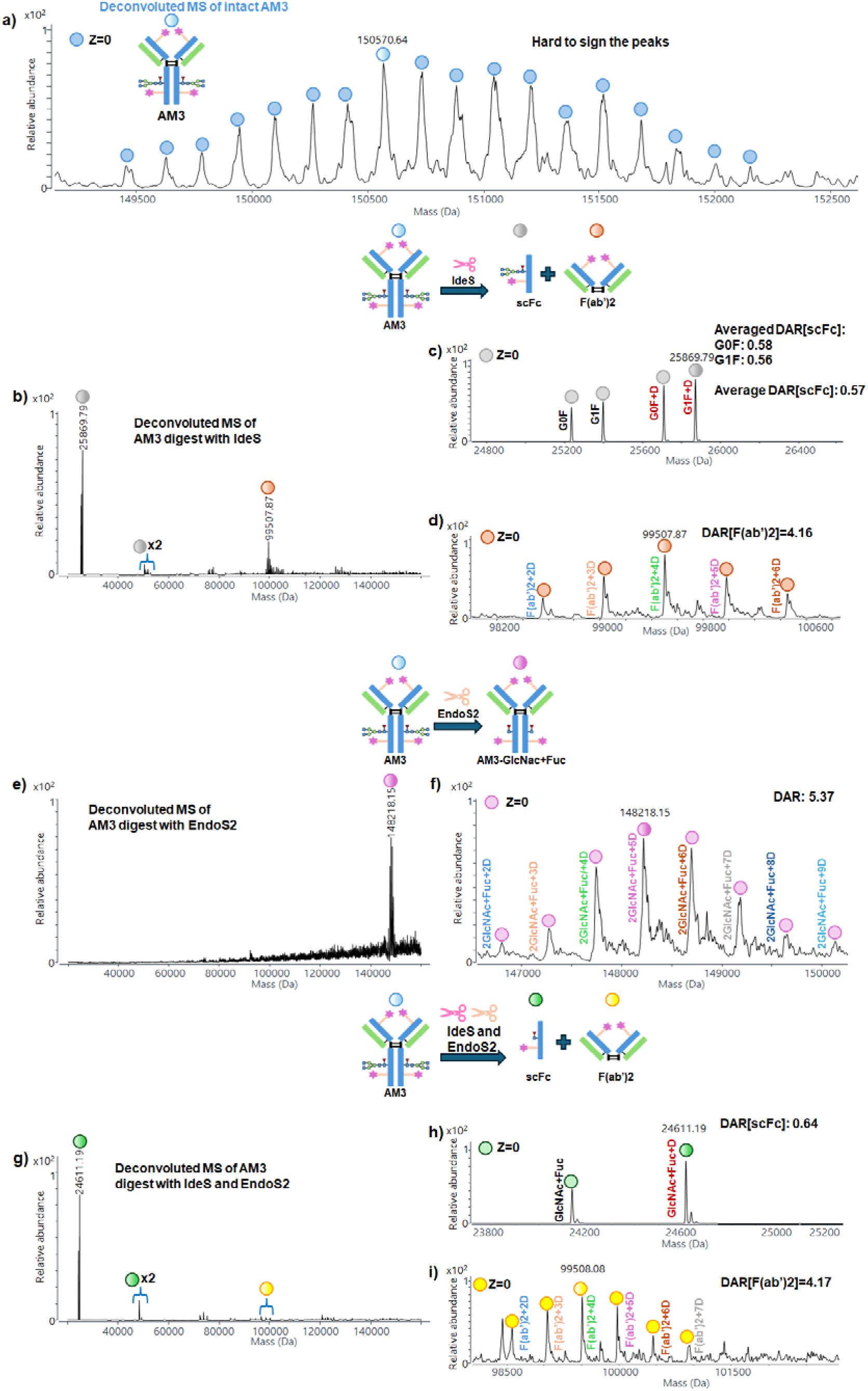
(a) Zoomed-in mass spectrum of intact **AM3**; (b) deconvoluted mass spectrum of **AM3** after microdroplet digestion with IdeS; zoomed-in deconvoluted MS spectra of (c) scFc and (d) F(ab’)2 subunits region of **AM3** after Ides microdroplet digestion; (e) deconvoluted mass spectrum of **AM3** after EndoS2 microdroplet digestion; (f) zoomed-in deconvoluted MS spectrum of **AM3** after microdroplet digestion with EndoS2; (g) deconvoluted mass spectrum of **AM3** after microdroplet digestion with IdeS and EndoS2; zoomed-in deconvoluted MS spectra of (h) scFc and (i) F(ab’)2 subunits region of **AM3** after microdroplet digestion with IdeS and EndoS2.

Highly conjugated NISTmAb–ADC mimic samples, **AM5** and **AM6**, were also examined. **AM5** and **AM6** exhibited highly complex intact MS spectra due to overlapping glycoform and NHS-PEG_4_-Biotin conjugation distributions, making direct DAR determination impractical (Figure S15); thus, enzymatic digestion strategies were applied. IdeS digestion enabled subunit-level DAR determination (Figure S16), while EndoS2 alone effectively simplified glycoform heterogeneity and provided reliable total DAR values (Figure S17), demonstrating its suitability for highly conjugated samples. Combined IdeS/EndoS2 digestion (Figure S18) further improved scFc accuracy, and all methods showed high digestion efficiency (>96%), confirming that even highly conjugated ADC mimics are compatible with microdroplet digestion. Based on the different enzyme microdroplet digestion approaches, the average DAR values were determined to be 12.19 for **AM5** and 13.55 for **AM6** (Table 1). The detailed discussion for the measurement is provided in SI. Although few commercial ADC drugs reach such high DAR values like **AM5** and **AM6**, investigation of this regime is still important considering future ADC development and to ensure highly conjugated payloads are amenable to microdroplet digestion and methodological completeness.

Therefore, for NISTmAb–ADC mimic samples measured above, accurate DAR values can be obtained from the subunit-level analysis regardless of the conjugation level, including low-conjugated samples (DAR < 3), moderately conjugated samples (3 < DAR < 10), and highly conjugated samples (DAR > 10), using IdeS digestion alone, EndoS2 digestion alone, or the combined IdeS/EndoS2 digestion strategy.

In addition to NISTmAb–based ADC mimics, this study also aimed to evaluate whether the microdroplet digestion approach could be applied to structurally more complex ADCs. Therefore, Cetuximab was selected as a model system. Unlike a typical IgG1, Cetuximab contains N-linked glycans that are sialylated in its Fab region (at Asn^88^), in addition to the conserved glycosylation site in the Fc region.^53^ As a result, its mass spectra are substantially more complex due to the various glycoforms. To facilitate comparison with Cetuximab–ADC mimics, we first characterized intact Cetuximab and its subunits generated by microdroplet digestion with different enzymes. Intact Cetuximab exhibited extensive glycoform heterogeneity (Figure S19) compared with NISTmAb, particularly in the F(ab’)2 region due to sialylated glycans, which complicated intact mass analysis. Microdroplet digestion with IdeS produced scFc and F(ab’)2 subunits, but the F(ab’)2 signal was relatively weak probably due to the presence of sialic-acid–containing glycans (Figure S20); and EndoS2-alone microdroplet digestion still led to multiple antibody glycoform peaks (Figure S21). Similarly, co-microdroplet digestion with IdeS and EndoS2 enhanced the scFc signal but no detectable F(ab’)2 signal was observed (Figure S22). To remove Fab-associated glycans and recover the F(ab’)2 signal, EndoF3 was introduced. EndoF3 is an endoglycosidase that cleaves within the chitobiose core of N-linked, fucosylated biantennary and triantennary complex oligosaccharides on glycoproteins. However, EndoF3-alone microdroplet digestion also showed multiple antibody glycoform peaks (Figure S23). Subsequent co-microdroplet digestion with IdeS and EndoF3 clearly resolved both scFc and F(ab’)2 subunits with simplified glycoforms (Figure S24), while combined EndoS2/EndoF3 digestion removed glycans from both Fab and Fc regions (Figure S25). Finally, one-pot microdroplet digestion using IdeS, EndoS2, and EndoF3 achieved efficient Fc cleavage and near-complete deglycosylation (∼96.7%), enabling robust subunit-level analysis of highly glycosylated Cetuximab (Figure S26). The relevant discussion is shown in SI.

To demonstrate our method capability of measuring DAR values for ADCs based on structurally complex antibodies such as Cetuximab, **AM7** was synthesized using Cetuximab and NHS-PEG_4_-Biotin (Figure S1b, details are provided in the SI). The MS spectrum and the deconvoluted spectrum are shown in Figures S27a-b. A zoomed-in view of the main peak region (Figure S27c) reveals extensive peak overlap, arising from multiple glycoforms on both scFc and F(ab’)2 regions and heterogeneous DAR distributions. For **AM7**, neither IdeS, EndoS2, nor their combination enabled total DAR determination due to severe ionization suppression and persistent spectral complexity, particularly arising from Fab glycans and extensive linker conjugation (Figures 3a-c and S28-S30). Even EndoF3 digestion, while confirming glycan removal, failed to sufficiently simplify the spectrum for reliable DAR analysis (Figure S31). Details are shown in SI. Based on prior observations with intact Cetuximab, combined digestion with EndoF3 and IdeS (Figure 3d) was expected to selectively enhance F(ab’)2 ionization by removing Fab glycans while leaving most Fc glycans intact. Indeed, this strategy successfully generated both scFc and F(ab’)2 subunits with sufficiently strong signals (Figures 3e–g and S32a). The scFc region (Figure 3f) exhibited mixed glycosylated and deglycosylated species, yielding a DAR[scFc] of 0.46. This value differs from that obtained via IdeS and EndoS2 digestion (0.59), likely due to glycoform interference. In contrast, the zoomed-in F(ab’)2 region (Figure 3g) clearly showed species conjugated with 2–6 linkers, yielding a DAR[F(ab’)2] of 3.89. The overall DAR calculated using this approach was therefore 2 × DAR[scFc] + DAR[F(ab’)2] = 4.81. Dual deglycosylation with EndoS2 and EndoF3 enabled complete glycan removal (Figure S33), yielding a simplified spectrum and an overall DAR of ∼5.19 for **AM7**. This value closely matched that obtained from IdeS and EndoF3 digestion, confirming consistency across methods. One-pot microdroplet digestion with IdeS, EndoS2, and EndoF3 (Figures 3h-j and S32b) provided accurate DAR[scFc] (∼0.61) by eliminating Fc glycan interference, highlighting the importance of Fc deglycosylation. However, due to persistently weak F(ab’)2 signals, DAR[F(ab’)2] was determined separately using IdeS and EndoF3 digestion. The digestion efficiencies for all microdroplet digestions using different enzymes exceeded 96% (Table S3), and the DAR values of the subunits, calculated using different methods, are summarized in Table S4. Details of **AM7** microdroplet digestion results are shown in SI. Accordingly, the most reliable DAR for **AM7** was calculated as 2 × 0.60 + 3.89 = 5.09, in excellent agreement with the value obtained from EndoS2 and EndoF3 digestion (5.19). A comparison of DAR values obtained by all methods is summarized in Table 2, yielding an average DAR of 5.03 with a %CV of 3.9%.

**Figure 3.**
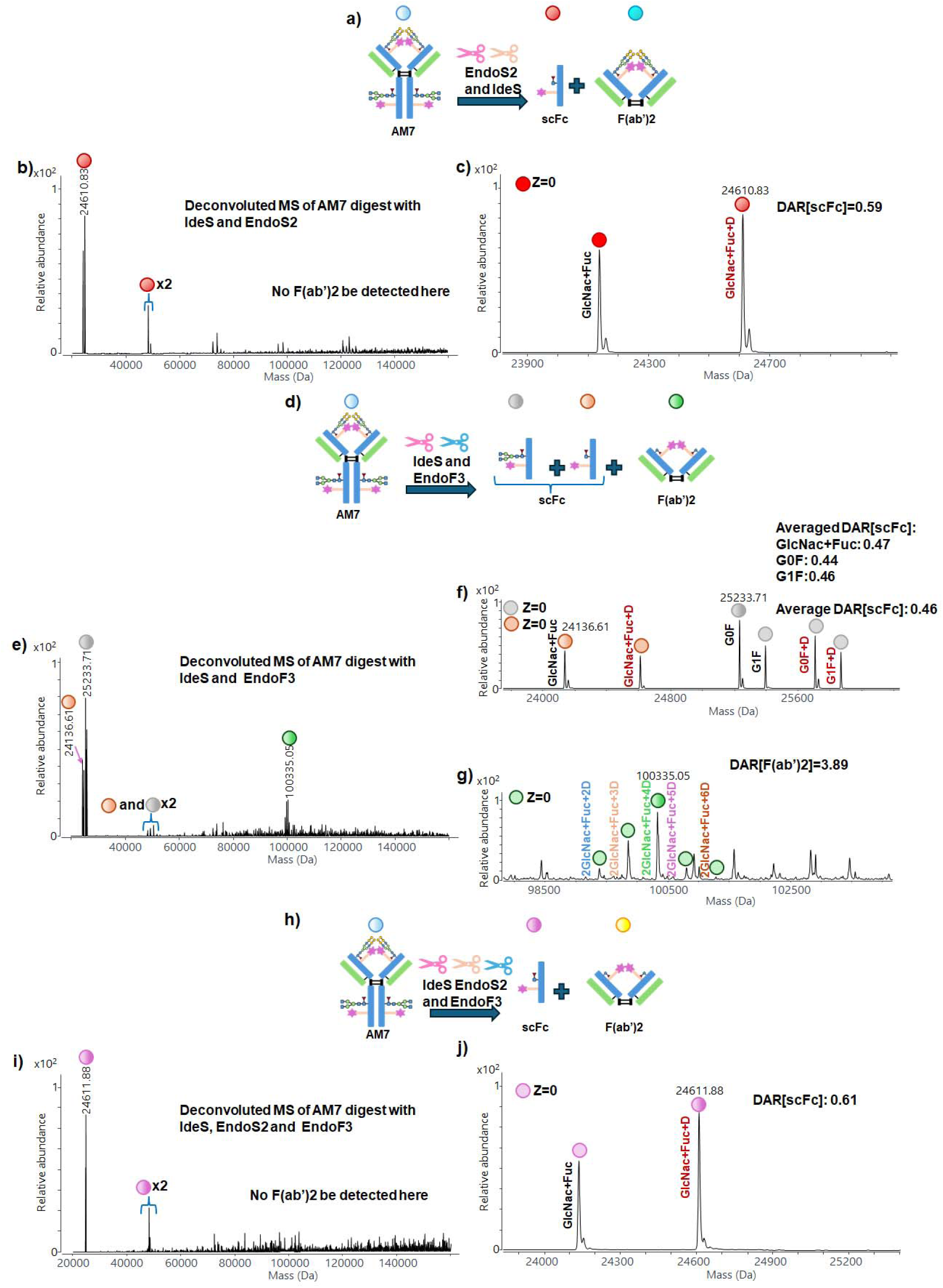
(a) Scheme showing microdroplet digestion of **AM7** with IdeS and EndoS2; (b) deconvoluted MS spectrum of **AM7** microdroplet digestion with IdeS and EndoS2; (c) zoomed-in deconvoluted MS spectrum of scFc; (d) scheme showing microdroplet digestion of **AM7** with IdeS and EndoF3; (e) Deconvoluted mass spectrum of **AM7** after microdroplet digestion with IdeS and EndoF3; zoomed-in deconvoluted MS spectra of (f) scFc and (g) F(ab’)2 subunits region of **AM7** after microdroplet digestion with IdeS and EndoF3; (h) **s**cheme showing microdroplet digestion of **AM7** with IdeS, EndoS2 and EndoF3; (i) deconvoluted mass spectrum of **AM7** after microdroplet digestion with IdeS, EndoS2 and EndoF3; (j) zoomed-in deconvoluted MS spectrum of scFc subunits region of **AM7** after microdroplet digestion with IdeS, EndoS2 and EndoF3.

**Table 2.** DAR values of Cetuximab-ADC mimic determined by microdroplet digestion using different enzymes.

| Samples | Cetuximab:NHS-PEG<sub>4</sub>-Biotin, reaction time | IdeS and EndoF3 | EndoS2 and EndoF3 | IdeS, EndoS2 and EndoF3 | Average DAR (%CV) |
| --- | --- | --- | --- | --- | --- |
| **AM7** | 1:10, 50 min | 4.81±0.07 | 5.19±0.10 | 5.09±0.07* | 5.03 (3.9%) |

\*Under this method, only the DAR of the scFc subunit could be directly determined, whereas the DAR of the F(ab')<sub>2</sub> subunit was obtained from digestion with IdeS and EndoF3.

Collectively, these results demonstrate that ultrafast microdroplet digestion coupled with direct MS analysis using the AJS source enables robust and accurate DAR determination for both NISTmAb- and Cetuximab-based ADC mimics (**AM1**–**AM7**), spanning low, moderate, and high conjugation regimes.

After validating the feasibility of rapid microdroplet digestion for the determination of DAR and subunit-level DAR using various ADC mimics, we next extended this strategy to a commercially approved ADC, Kadcyla (ado-trastuzumab emtansine). Kadcyla is a HER2-targeted ADC composed of the humanized anti-HER2 IgG1 antibody Trastuzumab, which is covalently linked (via lysine residues) to the microtubule inhibitor DM1 (an emtansine derivative) via a stable thioether linker, MCC (4-[N-maleimidomethyl]cyclohexane-1-carboxylate). On average, approximately 3.5 DM1 molecules are conjugated to each Trastuzumab molecule.^54^ Kadcyla is classified as a second-generation ADC and represents one of the most successful commercial ADCs to date. It was first approved by the U.S. Food and Drug Administration (FDA) in 2013 for the treatment of patients with HER2-positive metastatic breast cancer and was the first ADC approved for solid tumors. Owing to its well-characterized structure, clinically relevant DAR distribution, and widespread use as a benchmark ADC, Kadcyla serves as an ideal model system and real-world ADC for evaluating the performance of our microdroplet-based digestion and mass spectrometric DAR analysis.

Before evaluating Kadcyla, its unconjugated antibody, Trastuzumab, was systematically analyzed with digestion using different enzymes to establish the reference mass spectra. Intact Trastuzumab showed multiple glycoform peaks (148234.78 Da, Figure S34), confirming glycosylation heterogeneity. Digestion with IdeS generated both scFc and F(ab’)2 subunits with strong signals (Figure S35), while treatment with EndoS2 removed Fc glycans and reduced the mass to 145893.71 Da (Figure S36). Combined IdeS/EndoS2 microdroplet digestion produced scFc containing a single GlcNAc+Fuc glycoform with 98.8% efficiency, although the F(ab’)2 signal decreased (Figure S37). The detailed discussion is shown in SI.

Next, we analyzed Kadcyla. The molecular structure of Kadcyla is shown in Figure S38. Kadcyla is composed of the monoclonal antibody Trastuzumab conjugated at several lysine residues to the cytotoxic payload DM1 through a non-cleavable MCC linker, as mentioned above. As a result, attachment of a single MCC–DM1 moiety to Trastuzumab leads to an approximate mass increase of 957 Da, which enables direct DAR determination based on mass shifts observed in MS spectra.^54^ Figures S39 show the intact MS spectrum of Kadcyla and its corresponding deconvoluted spectrum. The main peak of Kadcyla is centered at 151095.18 Da (Figure S39b), which is significantly higher than that of intact Trastuzumab (148234.78 Da, Figure S34), confirming multiple conjugation. Upon zooming into the 148000–155000 Da region of the deconvoluted spectrum (Figure S39b), multiple peak clusters corresponding to Trastuzumab conjugated with 1–6 MCC–DM1 units can be clearly observed. Owing to the large mass increment introduced by each MCC–DM1 conjugation (∼957 Da), the conjugated glycoforms are well resolved and can be readily distinguished, in contrast to the overlapping peak patterns observed for ADC mimics prepared using EZ-Link NHS-PEG_4_-Biotin conjugation. After peak assignments (Figure S39c), the DAR values for individual glycoforms were calculated as follows: G0F/G0F, 3.23; G0F/G1F, 2.96; G1F/G1F, 3.14; and G1F/G2F, 3.12. The G2F/G2F glycoform conjugated with 2 or 6 MCC–DM1 units exhibited insufficient signal intensity and was therefore excluded from DAR calculation (Figure S39c). Based on the weighted average of all assigned glycoforms, the overall DAR of intact Kadcyla was determined to be 3.11. However, several minor satellite peaks (not assigned) were observed adjacent to the glycoform peaks, which were attributed to MCC linkers conjugated to the antibody without the DM1 payload. These species became more evident following enzymatic microdroplet digestion (as shown below). The presence of these peaks significantly interferes with accurate DAR determination, thereby motivating further DAR analysis at the subunit level after enzymatic microdroplet digestion. Kadcyla was first digested using IdeS, and the resulting MS spectrum is shown in Figure S40a. The corresponding deconvoluted spectrum is presented in Figure 4a, where the scFc signal remains highly intense. Upon magnification (Figure 4b), a small fraction of scFc species conjugated with one MCC–DM1 unit can be observed, yielding a DAR[scFc] of 0.06. In contrast, the F(ab’)2 signal intensity in Figure 4a is relatively low, which can be attributed to the presence of the bulky DM1 payload that suppresses its ionization. Nevertheless, after zooming into the corresponding mass region, F(ab’)2 species conjugated with 0–6 MCC–DM1 units are clearly resolved, allowing calculation of a DAR[F(ab’)2] of 3.39. Consequently, the overall DAR was calculated as 2 × DAR[scFc] + DAR[F(ab’)2] = 2 × 0.06 + 3.39 = 3.51, in excellent agreement with the theoretical value of 3.5. Notably, for each major peak in the deconvoluted spectra, a lower-intensity satellite peak exhibiting an ∼+220 Da mass shift was consistently observed. These minor peaks were attributed to MCC linkers conjugated to the antibody without the DM1 payload.^54^ Finally, Kadcyla was subjected to a one-pot microdroplet digestion with IdeS and EndoS2 digestion. The MS spectrum and deconvoluted spectrum are shown in Figures S40b and 4d, respectively. Deglycosylated scFc species exhibited high signal intensity, and the zoomed-in spectrum (Figure 4e) yielded a DAR[scFc] of 0.10, consistent with the value obtained from IdeS digestion alone. Although the enhanced scFc signal further suppressed the relative intensity of F(ab’)2, the latter could still be clearly resolved after magnification (Figure 4f), revealing conjugation with 0–6 MCC–DM1 units. The calculated

**Figure 4.**
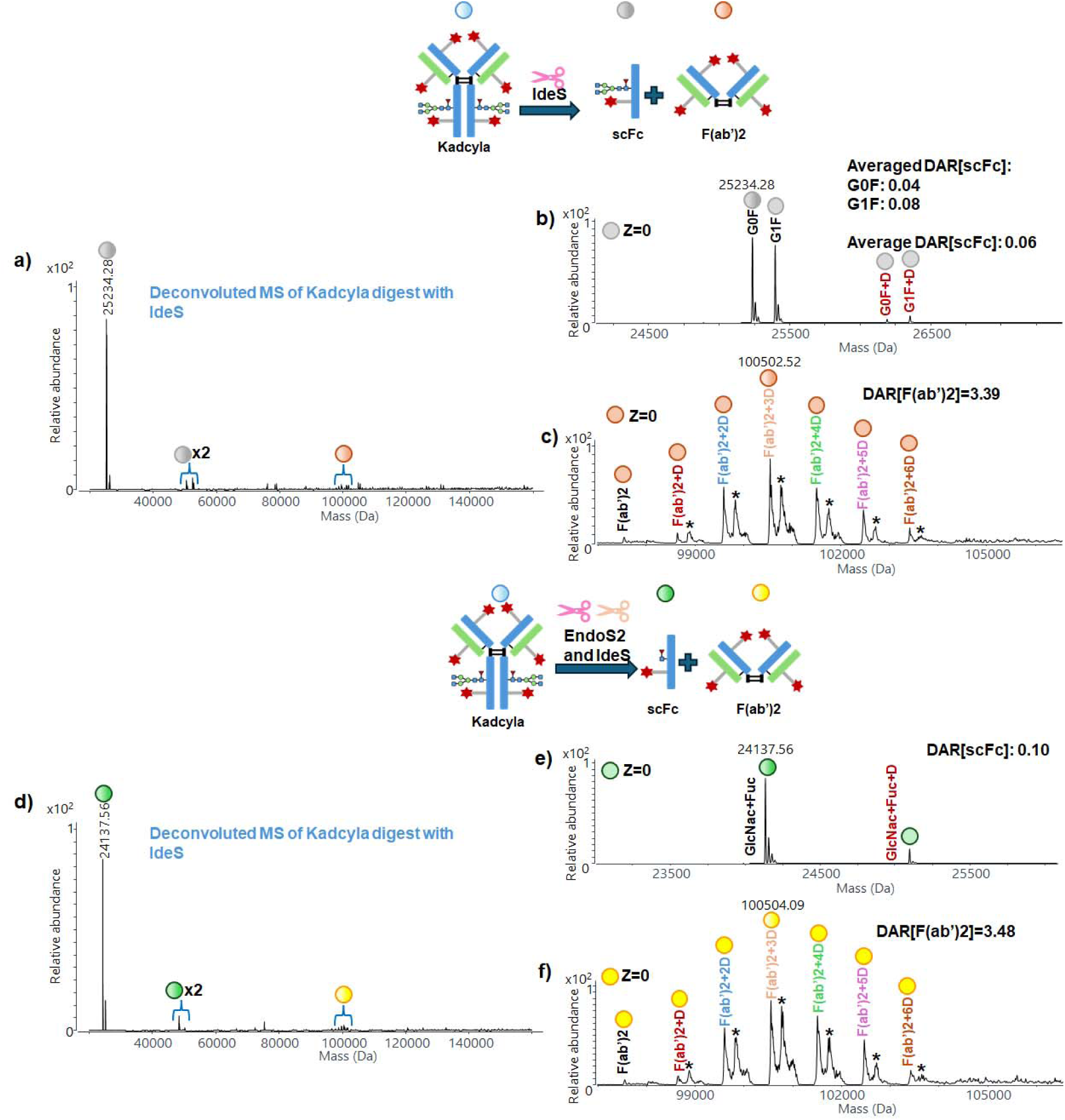
(a) Deconvoluted mass spectrum of Kadcyla after microdroplet digestion with IdeS; zoomed-in deconvoluted MS spectra of (b) scFc and (c) F(ab’)2 subunits region of Kadcyla after microdroplet digestion with IdeS; (d) deconvoluted mass spectrum of Kadcyla after microdroplet digestion with IdeS and EndoS2; zoomed-in deconvoluted MS spectra of (e) scFc and (f) F(ab’)2 subunits region of Kadcyla after microdroplet digestion with IdeS and EndoS2. **Footnote**: These minor peaks assigned as **\*** were attributed to MCC linkers conjugated to the antibody without the DM1 payload.

DAR[F(ab’)2] was 3.48. Across all measurements, the mass errors of the F(ab’)2 subunit and its corresponding conjugated species were consistently below 30 ppm. Accordingly, the overall DAR was determined to be 2 × 0.10 + 3.48 = 3.68. The total DAR values obtained from IdeS digestion alone (3.51) and combined IdeS/EndoS2 digestion (3.68) are highly consistent (%CV = 3.33%) and both closely match the theoretical DAR of Kadcyla (∼3.5), further validating the robustness and applicability of the ultrafast microdroplet digestion–MS workflow. All calculated DAR values and subunit-specific DARs are summarized in Table 1 and Table S1, respectively.

The DAR of Kadcyla was also independently verified using orthogonal LC/MS analytical methods. The details of validation are shown in SI. Briefly, intact Kadcyla was first analyzed by LC/MS to improve spectral clarity and enable accurate DAR determination (Figure S41). Compared with flow injection analysis (FIA), LC separation produced cleaner spectra and well-resolved deconvoluted peaks due to the removal of salt, revealing ADC species bearing 1–6 drugs and distinct glycoform distributions (Figure S41). Independent DAR calculations for major glycoform pairs yielded a consistent DAR of 3.31. As an orthogonal approach, TCEP reduction of Kadcyla (to produce light chain LC and heavy chain HC) followed by LC/MS analysis gave DAR[LC] = 0.38 and DAR[HC] = 1.48, corresponding to an overall DAR of 3.72 (Figure S42-S44). In addition, EndoS2 digestion and LC/MS analysis of deglycosylated species produced a DAR of 3.55 (Figure S45). The detailed discussion is shown in SI. All orthogonal results closely agreed with our microdroplet-based measurements (DAR: 3.51 and 3.68, Table 1). The overall DAR across methods (LC/MS and our method) was 3.56 with a %CV of 4.55%, demonstrating strong inter-method consistency and confirming the accuracy and reliability of the proposed microdroplet workflow. Additionally, although all orthogonal analytical methods employed here, which involved chromatographic separation, provided improved spectral quality, they required extensive sample preparation and longer analysis times. For example, TCEP reduction requires reagent preparation and approximately 30 min of reaction time, while EndoS2 in-solution digestion requires about 1 h, followed by an additional 18 min for chromatographic separation. In contrast, our microdroplet digestion method requires no additional sample preparation and achieves digestion within less than 1 ms. Overall, this approach offers a practical, efficient, and broadly applicable solution for DAR analysis, with significant potential for routine use in industrial ADC characterization workflows.

## Conclusions

In this work, we demonstrated an ultrafast microdroplet digestion–mass spectrometry strategy for rapid and reliable determination of DARs in ADCs. By exploiting the accelerated reaction kinetics within microdroplets generated in the AJS ion source, enzymatic digestion of intact antibodies and ADCs can be achieved on a sub-millisecond timescale, enabling direct online subunit-level mass spectrometric analysis with minimal sample preparation. This workflow substantially reduces total analysis time compared with conventional chromatographic or bulk digestion methods while preserving analytical accuracy and structural specificity. Systematic evaluation using NISTmAb-based ADC mimics spanning low (DAR < 3), moderate (3–10), and high (>10) conjugation levels demonstrated that IdeS-, EndoS2-, and combined enzymatic digestion strategies provide consistent DAR measurements with coefficients of variation generally below 5%. For all ADC mimics evaluated in this study, the microdroplet digestion efficiencies exceeded 96% under all conditions tested. Moreover, highly conjugated ADC mimics (DAR > 10) did not adversely affect the microdroplet digestion efficiency (higher than 96%, Table S2). The approach also proved applicable to structurally complex ADC systems, including Cetuximab-derived ADC mimics containing Fab-region glycans, for which the incorporation of EndoF3 enabled improved spectral simplification and reliable DAR determination. Furthermore, analysis of the commercial ADC Kadcyla yielded DAR values consistent with previously reported data, supporting the robustness and practical relevance of the method. Beyond DAR determination, the microdroplet-based platform enables rapid assessment of subunit-specific DAR distributions while minimizing sample consumption and preparation steps. These capabilities are particularly advantageous for ADC screening, conjugation optimization, formulation studies, and quality control applications. Overall, the integration of microdroplet reaction acceleration with direct mass spectrometric detection provides a powerful analytical framework for fast, sensitive, and structurally informative ADC characterization, with strong potential to streamline ADC development workflows and expand real-time analytical capabilities in biopharmaceutical analysis.

## Supporting Information

Information about sample preparation, instrument conditions, data analysis, additional MS spectra and detailed microdroplet digestion result discussions are included.

## Acknowledgment

HPG acknowledges the support from Johnson & Johnson. HC, YY, JPS, TY, MTAH and JP thank NSF (CHE-2505727) for financial support. MB thanks the Fullbright Visiting Scholar Program. The authors thank Amir Liba from Agilent Technologies for the research collaboration and for providing instrument support. The authors also thank Agilent Technologies for the ACT-UR Award.

## References

1. Thomas A, Teicher BA, Hassan R. Antibody-drug conjugates for cancer therapy. Lancet Oncol. 2016;17(6):e254–e262.

2. Lambert JM, Berkenblit A. Antibody-Drug Conjugates for Cancer Treatment. Annu Rev Med. 2018;69:191–207.

3. Norsworthy KJ, Ko CW, Lee JE, et al. FDA Approval Summary: Mylotarg for Treatment of Patients with Relapsed or Refractory CD33-Positive Acute Myeloid Leukemia. Oncologist. 2018;23(9):1103–1108.

4. Fu Z, Li S, Han S, Shi C, Zhang Y. Antibody drug conjugate: the “biological missile” for targeted cancer therapy. Signal Transduct Target Ther. 2022;7(1):93.

5. Maecker H, Jonnalagadda V, Bhakta S, Jammalamadaka V, Junutula JR. Exploration of the antibody–drug conjugate clinical landscape. In MAbs, 2023; Taylor & Francis: Vol.15, p 2229101..

6. Behrens CR, Ha EH, Chinn LL, et al. Antibody-Drug Conjugates (ADCs) Derived from Interchain Cysteine Cross-Linking Demonstrate Improved Homogeneity and Other Pharmacological Properties over Conventional Heterogeneous ADCs. Mol Pharm. 2015;12(11):3986–3998.

7. Tang H, Liu Y, Yu Z, et al. The Analysis of Key Factors Related to ADCs Structural Design. Front Pharmacol. 2019;10:373.

8. Wakankar A, Chen Y, Gokarn Y, Jacobson FS. Analytical methods for physicochemical characterization of antibody drug conjugates. MAbs. 2011;3(2):161–172.

9. Hamblett KJ, Senter PD, Chace DF, et al. Effects of drug loading on the antitumor activity of a monoclonal antibody drug conjugate. Clinical Can.Res.. 2004;10(20):7063–7070.

10. King HD, Dubowchik GM, Mastalerz H, et al. Monoclonal antibody conjugates of doxorubicin prepared with branched peptide linkers: inhibition of aggregation by methoxytriethyleneglycol chains. J. Med. Chem. 2002;45(19):4336–4343.

11. Strop P, Delaria K, Foletti D, et al. Site-specific conjugation improves therapeutic index of antibody drug conjugates with high drug loading. Nat. Biotechnol. 2015;33(7):694–696.

12. Debaene F, Boeuf A, Wagner-Rousset E, et al. Innovative native MS methodologies for antibody drug conjugate characterization: High resolution native MS and IM-MS for average DAR and DAR distribution assessment. Anal. Chem. 2014;86(21):10674–10683.

13. Chen Y. Drug-to-Antibody Ratio (DAR) by UV/Vis Spectroscopy. In: Ducry L, ed. Antibody-Drug Conjugates. Totowa, NJ: Humana Press; 2013:267–273.

14. Andris S, Rudt M, Rogalla J, Wendeler M, Hubbuch J. Monitoring of antibody-drug conjugation reactions with UV/Vis spectroscopy. J. Biotechnol. 2018;288:15–22.

15. Le LN, Moore JM, Ouyang J, Chen X, Nguyen MD, Galush WJ. Profiling antibody drug conjugate positional isomers: a system-of-equations approach. Anal. Chem. 2012;84, 7479–7486.

16. Bobaly B, Randazzo GM, Rudaz S, Guillarme D, Fekete S. Optimization of non-linear gradient in hydrophobic interaction chromatography for the analytical characterization of antibody-drug conjugates. J. Chromatogr. A. 2017;1481:82–91.

17. Song W, Yin L, Ren J, et al. Simultaneous Analysis of the Drug-to-Antibody Ratio, Free-Drug-Related Impurities, and Purity of Antibody-Drug Conjugates Based on Size Exclusion Chromatography. Anal. Chem. 2025;97, 9748–9754.

18. Liu JZ, Du CY, Gao H, Wang H, Hu F, Fang WJ. An Underlying Cause and Solution to the Poor Size Exclusion Chromatography Performance of Antibody-Drug Conjugates. Pharm. Res. 2024;41, 2299–2317.

19. Redman EA, Mellors JS, Starkey JA, Ramsey JM. Characterization of Intact Antibody Drug Conjugate Variants Using Microfluidic Capillary Electrophoresis-Mass Spectrometry. Anal. Chem. 2016, 88, 2220–2226.

20. Wang Z, Liu X, Muther J, James JA, Smith K, Wu S. Top-down Mass Spectrometry Analysis of Human Serum Autoantibody Antigen-Binding Fragments. Sci. Rep. 2019;9(1):2345.

21. Fang M, Wang Z, Norris K, James JA, Wu S, Smith K. Hydrogen-Deuterium Exchange Mass Spectrometry Reveals a Novel Binding Region of a Neutralizing Fully Human Monoclonal Antibody to Anthrax Protective Antigen. Toxins (Basel). 2022;14(2).

22. Duan Z, Wang C, Wang Y. Optimization of q Value for Sensitive Detection of Uridine and Thymidine Nucleosides by MS(3). J. Am. Soc. Mass Spectrom. 2025;36(10):2017–2021.

23. Liu FC, Lee J, Pedrete T, Panczyk EM, Pengelley S, Bleiholder C. Differential glycosylation does not modulate the conformational heterogeneity of a humanised IgGk NIST monoclonal antibody. Chem. Commun. 2024;60, 0740–10743.

24. Wagner-Rousset E, Janin-Bussat MC, Colas O, et al. Antibody-drug conjugate model fast characterization by LC-MS following IdeS proteolytic digestion. MAbs. 2014;6(1):273–285.

25. Tian Y, Lippens JL, Netirojjanakul C, Campuzano IDG, Ruotolo BT. Quantitative collision-induced unfolding differentiates model antibody-drug conjugates. Protein Sci. 2019;28(3):598–608.

26. Nagy G, Attah IK, Conant CR, et al. Rapid and Simultaneous Characterization of Drug Conjugation in Heavy and Light Chains of a Monoclonal Antibody Revealed by High-Resolution Ion Mobility Separations in SLIM. Anal. Chem. 2020;92(7):5004–5012.

27. Aguilan J, Madrid-Aliste C, Zandkarimi F, et al. Direct Injection Mass Spectrometry and iFishMass for the High-Throughput Analysis of Antibody Modifications. ACS Pharmacology & Translational Science. 2026, 9, 165–176

28. Yang Y, Hassan MT, Yaroshuk T, et al. Ultrafast PFAS degradation using oxidant-containing microdroplets. Chem. Commun. 2025;61(90):17629–17632.

29. Wang Y, Luo J, Fang YG, et al. Catalyst-Free Nitrogen Fixation by Microdroplets through a Radical-Mediated Disproportionation Mechanism under Ambient Conditions. J. Am. Chem Soc. 2025;147(3):2756–2765.

30. Shi Z, Nie J, Chen ZC, et al. Synergistically Activating N(2) and CO(2) at Water Microdroplet Interfaces. Angew Chem Int Ed Engl. 2025:e19068.

31. Yao H, Zhou D, VanSickle S, et al. Microdroplet Mass Spectrometry Enables Rapid and Sensitive Measurement of Glucose Homeostasis for Elucidating Mechanism of Drug Action. Anal Chem. 2025;97(36):19480–19488.

32. Ai Y, Xu J, Gunawardena HP, Zare RN, Chen H. Investigation of Tryptic Protein Digestion in Microdroplets and in Bulk Solution. J Am Soc Mass Spectrom. 2022;33(7):1238–1249.

33. Breitfeld M, Dietsche CL, Saucedo-Espinosa MA, Berlanda SF, Dittrich PS. Ultrafast Formation of Microdroplet Arrays with Chemical Gradients for Label-Free Determination of Enzymatic Reaction Kinetics. Small. 2025;21(26):e2410275.

34. Cheng H, Tang S, Yang T, Xu S, Yan X. Accelerating Electrochemical Reactions in a Voltage-Controlled Interfacial Microreactor. Angew Chem Int Ed Engl. 2020;59(45):19862–19867.

35. Li Y, Yan X, Cooks RG. The Role of the Interface in Thin Film and Droplet Accelerated Reactions Studied by Competitive Substituent Effects. Angew Chem Int Ed Engl. 2016;55(10):3433–3437.

36. Gnanamani E, Yan X, Zare RN. Chemoselective N-Alkylation of Indoles in Aqueous Microdroplets. Angew Chem Int Ed Engl. 2020;59(8):3069–3072.

37. Li J, Aikonen S, Morato NM, et al. Catalyst-Free C-N Coupling under Ambient Conditions via High-Throughput Microdroplet Reactions. J Org Chem. 2025;90(51):18172–18180.

38. Ghosh J, Morato NM, LeFever WA, Cooks RG. Accelerated and Green Synthesis of N,S-and N,O-Heterocycles in Microdroplets. J Am Chem Soc. 2026;148(3):2920–2929.

39. Holden DT, Shira BA, Edwards MQ, Morato NM, Cooks RG. Mechanisms of ionization and of chemical reactions in charged microdroplets. Chem Sci. 2025;16(37):17020–17033.

40. Lee JK, Kim S, Nam HG, Zare RN. Microdroplet fusion mass spectrometry for fast reaction kinetics. Proc Natl Acad Sci U S A. 2015;112(13):3898–3903.

41. Gong K, Nandy A, Song Z, et al. Revisiting the Enhanced Chemical Reactivity in Water Microdroplets: The Case of a Diels-Alder Reaction. J Am Chem Soc. 2024;146(46):31585–31596.

42. Rainer T, Eidelpes R, Tollinger M, Muller T. Microdroplet Mass Spectrometry Enables Extremely Accelerated Pepsin Digestion of Proteins. J Am Soc Mass Spectrom. 2021;32(7):1841–1845.

43. Ma C-H, Chen C-L, Hsu C-C. Real-time bottom-up characterization of protein mixtures enabled by online microdroplet-assisted enzymatic digestion (MAED). Chem. Comm. 2023;59(84):12585–12588.

44. Zhong X, Chen H, Zare RN. Ultrafast enzymatic digestion of proteins by microdroplet mass spectrometry. Nat. Commun. 2020;11(1):1049.

45. Xiao M, Yang Y, Schladebeck A, et al. Rapid Antibody Structural Characterization and Quantification via Microdroplet Trypsin Digestion. J. Am Soc Mass Spectrom. 2026, in press.

46. Gunawardena HP, Ai Y, Gao J, Zare RN, Chen H. Rapid Characterization of Antibodies via Automated Flow Injection Coupled with Online Microdroplet Reactions and Native-pH Mass Spectrometry. Anal Chem. 2023;95(6):3340–3348.

47. Zhao P, Gunawardena HP, Zhong X, Zare RN, Chen H. Microdroplet Ultrafast Reactions Speed Antibody Characterization. Anal Chem. 2021;93(8):3997–4005.

48. Yang Y, Xiao M, Lau J, et al. Ultrafast Microdroplet Digestion of Antibodies with Fc-Silencing Mutations. Anal Chem. 2025;97(24):12813–12823.

49. Anapindi KDB, Hsieh EJ, Walker T, Lau J, Tomazela D. High-Speed Automated Microdroplet Reactions with Ion-Mobility for Rapid Therapeutic Protein Characterization. J Am Soc Mass Spectrom. 2025;36(11):2397–2404.

50. Tang Y, Tang F, Yang Y, et al. Real-Time Analysis on Drug-Antibody Ratio of Antibody-Drug Conjugates for Synthesis, Process Optimization, and Quality Control. Sci Rep. 2017;7(1):7763.

51. Drago JZ, Modi S, Chandarlapaty S. Unlocking the potential of antibody–drug conjugates for cancer therapy. *Nature Rev*. Clinical Oncology. 2021;18(6):327–344.

52. Beck A, Goetsch L, Dumontet C, Corvaïa N. Strategies and challenges for the next generation of antibody–drug conjugates. Nature Rev. Drug discovery. 2017;16(5):315–337.

53. Liu S, Gao W, Wang Y, et al. Comprehensive N-Glycan Profiling of Cetuximab Biosimilar Candidate by NP-HPLC and MALDI-MS. PLoS One. 2017;12(1):e0170013.

54. Chen L, Wang L, Shion H, et al. In-depth structural characterization of Kadcyla(R) (ado-trastuzumab emtansine) and its biosimilar candidate. MAbs. 2016;8(7):1210–1223.

