## Supporting information for "Rapid Determination of Drug-to-Antibody Ratios in Antibody–Drug Conjugates Using Ultrafast Microdroplet Digestion Technology"

**Table of Contents**


| | Page number |
|---|---|
| Chemicals and materials | S-4 |
| Sample desalting | S-4 |
| Synthesis conditions for ADC mimics with different DAR values | S-4 |
| LC/MS analysis of Kadcyla | S-5 |
| Table S1 | S-6 |
| Table S2 | S-7 |
| Table S3 | S-8 |
| Table S4 | S-9 |
| Figure S1 | S-10 |
| Figure S2 | S-11 |
| Figure S3 | S-12 |
| Figure S4 | S-13 |
| Figure S5 | S-14 |
| Figure S6 | S-15 |
| Figure S7 | S-16 |
| Figure S8 | S-17 |
| Figure S9 | S-18 |
| Figure S10 | S-19 |
| EndoS2 microdroplet digestion of **AM3** and **AM4** | S-20 |
| Figure S11 | S-21 |
| Figure S12 | S-22 |
| Figure S13 | S-23 |
| Figure S14 | S-24 |
| Microdroplet digestion of **AM5** and **AM6** | S-25 |
| Figure S15 | S-27 |
| Figure S16 | S-28 |
| Figure S17 | S-29 |
| Figure S18 | S-30 |
| Intact Cetuximab microdroplet digestion | S-31 |
| Figure S19 | S-35 |
| Figure S20 | S-36 |
| Figure S21 | S-37 |
| Figure S22 | S-38 |
| Figure S23 | S-39 |
| Figure S24 | S-40 |
| Figure S25 | S-41 |
| Figure S26 | S-42 |
| Figure S27 | S-43 |
| **AM7** microdroplet digestion results | S-44 |

| | |
|---|---|
| Figure S28 | S-46 |
| Figure S29 | S-47 |
| Figure S30 | S-48 |
| Figure S31 | S-49 |
| Figure S32 | S-50 |
| Figure S33 | S-51 |
| Intact Trastuzumab microdroplet digestion | S-52 |
| Figure S34 | S-53 |
| Figure S35 | S-54 |
| Figure S36 | S-55 |
| Figure S37 | S-56 |
| Figure S38 | S-57 |
| Figure S39 | S-58 |
| Figure S40 | S-59 |
| Orthogonal analytical methods to verify Kadcyla's DAR | S-60 |
| Figure S41 | S-63 |
| Figure S42 | S-64 |
| Figure S43 | S-65 |
| Figure S44 | S-66 |
| Figure S45 | S-67 |
| References | S-68 |

### Chemicals and materials

Enzymes including FabRICATOR (IdeS) and GlycINATOR (EndoS2) were purchased from Genovis Inc. (Cambridge, MA, USA). Endoglycosidase F3 (EndoF3) was purchased from New England Biolabs (Cambridge, MA, USA). EZ-Link NHS-$PEG_4$-Biotin which used NHS-$PEG_4$ as the linker and biotin as the payload (the linker+biotin) and Zeba Spin Desalting Columns were purchased from Thermo Scientific (Carlsbad, CA, USA). 1.0 M 4-(2-hydroxyethyl)-1-piperazineethanesulfonic acid (HEPES buffer) was purchased from Alfa Aesar (Carlsbad, CA, USA). Ammonium bicarbonate (ABC, ≥98.5%) and tris(2-carboxyethyl)phosphine hydrochloride (TCEP) were purchased from Sigma-Aldrich (Burlington, MA, USA). Antibodies including humanized $IgG1_k$ monoclonal antibody (NISTmAb), Cetuximab, and Trastuzumab were purchased from Sigma-Aldrich (Burlington, MA, USA). Commercial ADC ado-trastuzumab emtansine (Kadcyla) was purchased from MedChemExpress (Monmouth Junction, NJ, USA). Molecular weight cutoff (MWCO) filter (0.5 mL, 10k) was purchased from Amicon (Miami, FL, USA). ACQUITY UPLC Protein BEH C4 was purchased from Waters (Milford, MA).

### Sample desalting

For the intact antibodies (NISTmAb, Cetuximab and Trastuzumab) and commercial ADC Kadcyla, we first used 10 kDa MWCO filter for desalting. In this process, water was used as the buffer solution, and desalting was performed four times. After adding the water each time, centrifugation was carried out at 14000 rcf for 10 min. Subsequently, the obtained stock solution was diluted to 0.5 μg/μL using DI water.

### Synthesis conditions for ADC mimics with different DAR values

Low-conjugation samples (DAR < 3) were prepared under the following conditions: NISTmAb: NHS-$PEG_4$-Biotin molar ratios of 1:5 with a reaction time of 30 min (**AM1**) and 1:20 with a reaction time of 30 min (**AM2**). Moderately conjugated samples (3 < DAR < 10), corresponding to commercially relevant ADCs, were prepared using NISTmAb: NHS-$PEG_4$-Biotin molar ratios of 1:10 with a reaction time of 50 min (**AM3**) and 1:20 with a reaction time of 50 min (**AM4**). Highly conjugated samples (DAR > 10) were prepared using NISTmAb: NHS-$PEG_4$-Biotin molar ratios of 1:40 (**AM5**), and 1:50 (**AM6**), each with a reaction time of 30 min. All NISTmAb-to- NHS-$PEG_4$-Biotin ratios reported refer to molar ratios and the reaction scheme is shown in Figure S1a.

We next applied the established strategy for preparing moderately conjugated Cetuximab–ADC mimic within the most clinically relevant DAR range (3–10). A Cetuximab–ADC mimic was synthesized using Cetuximab: NHS-$PEG_4$-Biotin molar ratio of 1:10 with a reaction time of 50 min (**AM7**), as illustrated in Figure S1b.

**LC/MS analysis of Kadcyla**

For DAR analysis of reduced Kadcyla, the interchain disulfide bonds of Kadcyla were first reduced to generate two light chains (LCs) and two heavy chains (HCs). A 20 mM tris(2-carboxyethyl)phosphine (TCEP) solution was prepared in water. Subsequently, 10 µL of Trastuzumab or Kadcyla (0.5 µg/µL) was mixed with 10 µL of 20 mM TCEP and incubated at 37 °C with shaking for 30 min. The resulting reaction products were then subjected to LC/MS analysis for chain-level separation and DAR determination.

For the bulk digestion of Kadcyla with EndoS2, the ratio of Kadcyla:EndoS2=1 µg : 2 units, and incubated in 37 °C for 1 h.

Liquid chromatography–mass spectrometry (LC/MS) was employed as an orthogonal method to determine the DAR of reduced Kadcyla and to validate the results obtained from the microdroplet approach. LC separations were performed using an ultra-performance liquid chromatography system (UPLC, Agilent 1290 Infinity II) equipped with a reversed-phase column (ACQUITY UPLC Protein BEH C4, Waters, Milford, MA). Mobile phase A consisted of water containing 0.1% formic acid, and mobile phase B consisted of acetonitrile containing 0.1% formic acid, with a constant flow rate of 0.7 mL/min. The gradient program was as follows: 0–0.2 min, 10% B; 0.2–15 min, linear increase from 10% to 40% B; 15–17 min, linear increase from 40% to 95% B; 17–17.01 min, linear decrease from 95% to 10% B; and 17.01–18 min, 10% B. The total run time for each sample was 18 min. To minimize potential interference from residual TCEP on MS detection, the LC effluent was diverted to waste during the first 1 min of the run and subsequently redirected to the mass spectrometer. The injection volume of each run was 2 µL.

**Table S1**. Subunit DAR values of NISTmAb-ADC mimics and Kadcyla determined by microdroplet digestion using different enzymes

| Samples | Subunits | IdeS | IdeS and EndoS2 |
|---|---|---|---|
| **AM1** | scFc | 0±0.00 | 0.04±0.00 |
| | F(ab')2 | 0.45±0.02 | 0.40±0.01 |
| **AM2** | scFc | 0.44±0.03 | 0.40±0.02 |
| | F(ab')2 | 1.83±0.05 | 1.79±0.06 |
| **AM3** | scFc | 0.57±0.03 | 0.64±0.05 |
| | F(ab')2 | 4.16±0.08 | 4.17±0.10 |
| **AM4** | scFc | 1.16±0.05 | 1.10±0.08 |
| | F(ab')2 | 7.24±0.11 | 7.16±0.14 |
| **AM5** | scFc | 1.15±0.07 | 1.40±0.04 |
| | F(ab')2 | 9.81±0.30 | 9.83±0.29 |
| **AM6** | scFc | 1.75±0.12 | 1.86±0.06 |
| | F(ab')2 | 10.01±0.41 | 10.02±0.39 |
| **Kadcyla** | scFc | 0.06±0.00 | 0.10±0.00 |
| | F(ab')2 | 3.39±0.11 | 3.48±0.08 |

**Table S2.** Microdroplet digestion efficiencies of NISTmAb, NISTmAb-ADC mimics and Kadcyla with different enzymes

| Samples | IdeS | EndoS2 | IdeS + EndoS2 |
|---|---|---|---|
| **NISTmAb** | 98.2±1.11 % | 96.2±2.27 % | 96.5±1.63 % |
| **AM1** | 96.5±1.51 % | 97.3±1.30 % | 96.9±2.24 % |
| **AM2** | 98.4±1.62 % | 97.1±2.51 % | 98.3±1.11 % |
| **AM3** | 97.9±1.10 % | 97.0±1.50 % | 98.2±1.32 % |
| **AM4** | 99.3±0.66 % | 98.1±0.92 % | 98.8±1.12 % |
| **AM5** | 97.3±1.55 % | 98.0±1.22 % | 97.1±0.84 % |
| **AM6** | 98.2±1.39 % | 96.9±2.10 % | 98.8±1.20 % |
| **Kadcyla** | 96.3±1.45 % | - | 97.7±2.01 % |

**Table S3.** Microdroplet digestion efficiencies of Cetuximab and **AM7** with different enzymes.

| **Samples** | IdeS | EndoS2 | IdeS + EndoS2 | EndoF3 | IdeS+EndoF3 | IdeS+EndoS2 +EndoF3 |
|---|---|---|---|---|---|---|
| **Cetuximab** | 96.7±0.73 % | 97.1±2.05 % | 97.7±0.38 % | 97.0±1.18 % | 96.2±0.72 % | 96.7±1.41 % |
| **AM7** | 97.8±1.03 % | 96.2±2.30 % | 98.4±1.05 % | 96.3±1.31 % | 98.3±1.81 % | 97.8±1.97 % |

**Table S4**. Subunit DAR values of Cetuximab-ADC mimic (**AM7**) determined using different enzymes microdroplet digestion

| **Samples** | **Subunits** | **with IdeS** | **IdeS and EndoS2** | **IdeS and EndoF3** | **IdeS, EndoS2 and EndoF3** |
|---|---|---|---|---|---|
| **AM7** | scFc | 0.72±0.04 | 0.59±0.07 | 0.46±0.09 | 0.61±0.04 |
| | F(ab')2 | - | - | 3.89±0.10 | - |

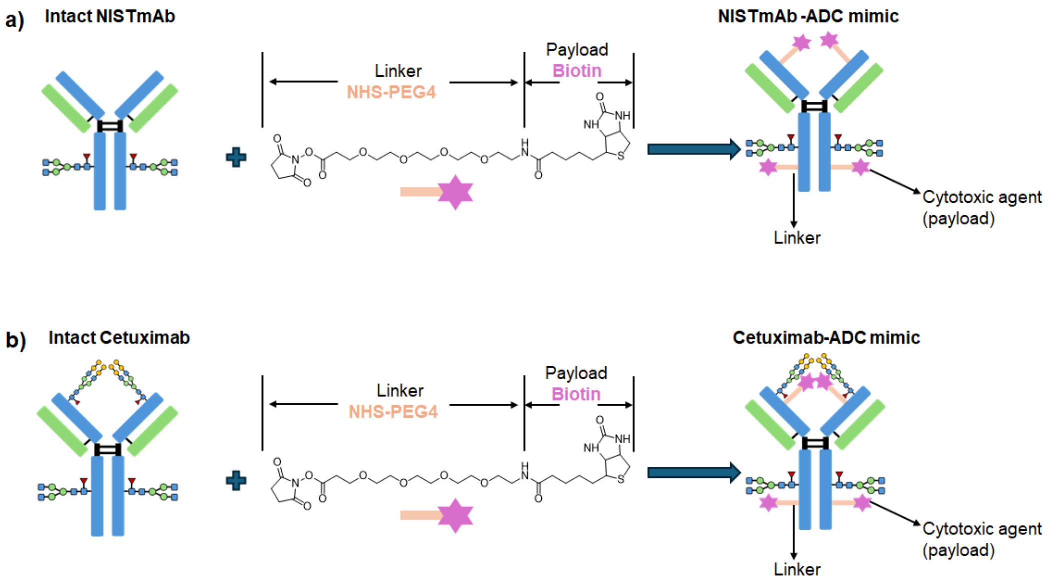


**Figure S1**. Scheme showing the syntheses of (a) NISTmAb-ADC mimics and (b) Cetuximab-ADC mimics.

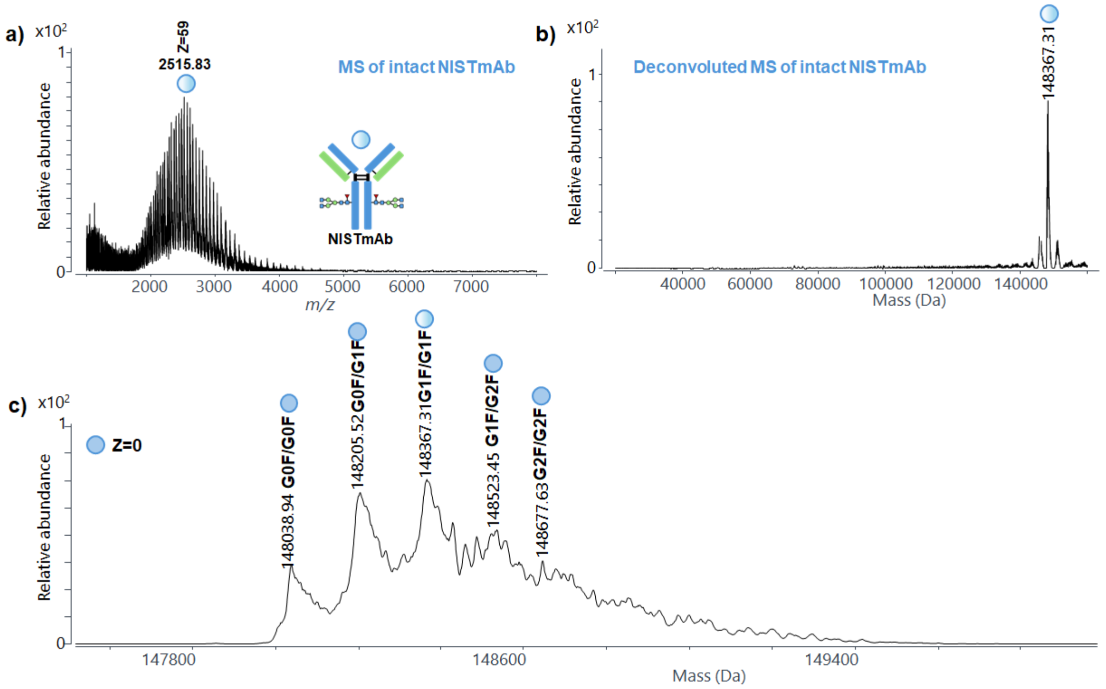


**Figure S2.** (a) Mass spectrum of intact NISTmAb; (b) deconvoluted mass spectrum of intact NISTmAb; (c) zoomed-in deconvoluted mass spectrum in the range of 147800–149400 Da.

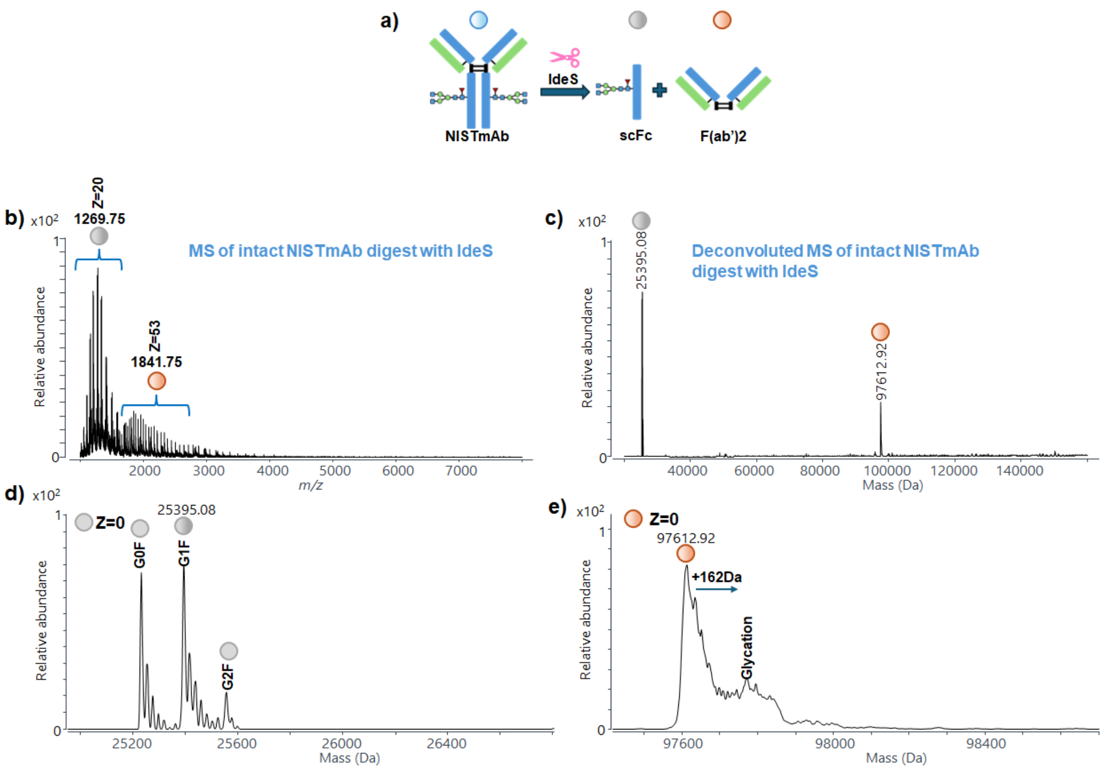


**Figure S3.** (a) Scheme showing the digestion of intact NISTmAb with IdeS; (b) MS spectrum of intact NISTmAb microdroplet digestion with IdeS; (c) deconvoluted MS spectrum of intact NISTmAb microdroplet digestion with IdeS; zoomed-in deconvoluted MS spectra of (d) scFc and (e) F(ab')2 subunits. The glycation peak observed in (e) was probably due to the presence of trace glucose in the sample which modified the antibody.

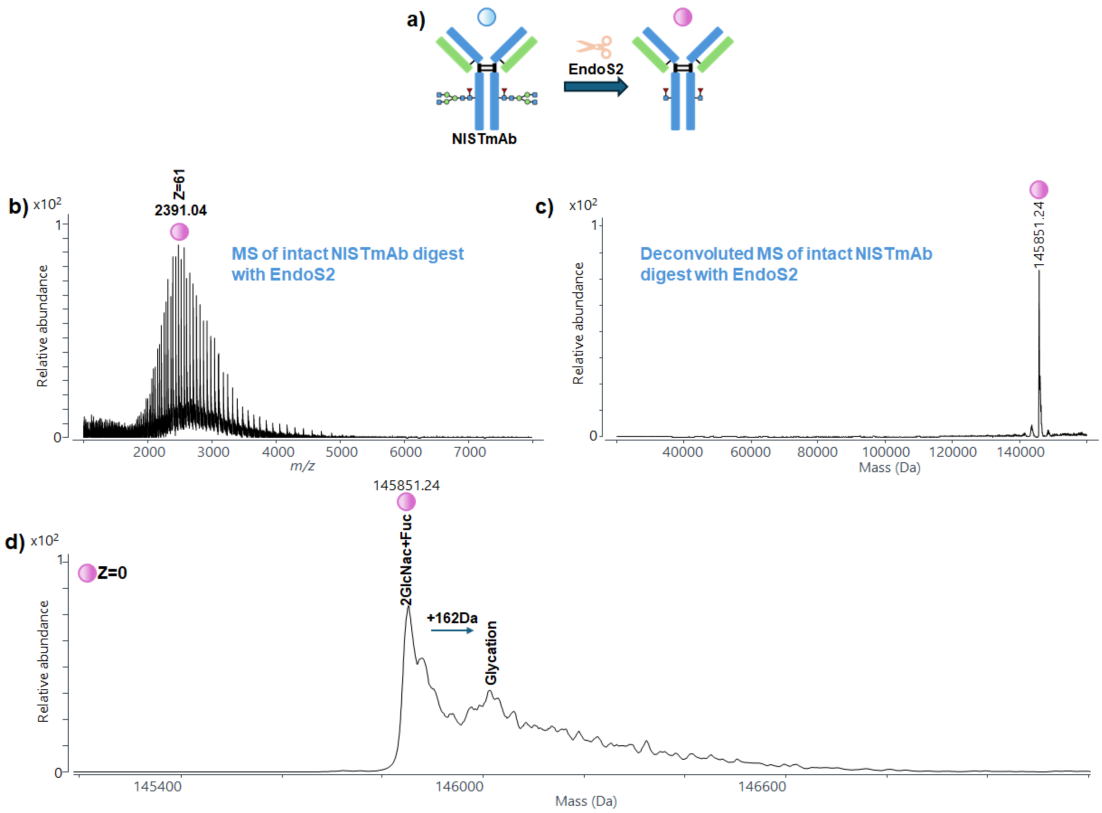


**Figure S4.** (a) Scheme showing the digestion of intact NISTmAb with EndoS2; (b) MS spectrum of intact NISTmAb microdroplet digestion with EndoS2; (c) deconvoluted MS spectrum of intact NISTmAb microdroplet digestion with EndoS2; (d) zoomed-in deconvoluted MS spectrum of intact NISTmAb microdroplet digestion with EndoS2 which only shows GlcNAc+Fuc signal along with a small glycation peak. The glycation peak observed in (d) was probably due to the presence of trace glucose in the sample which modified the antibody, as explained above.

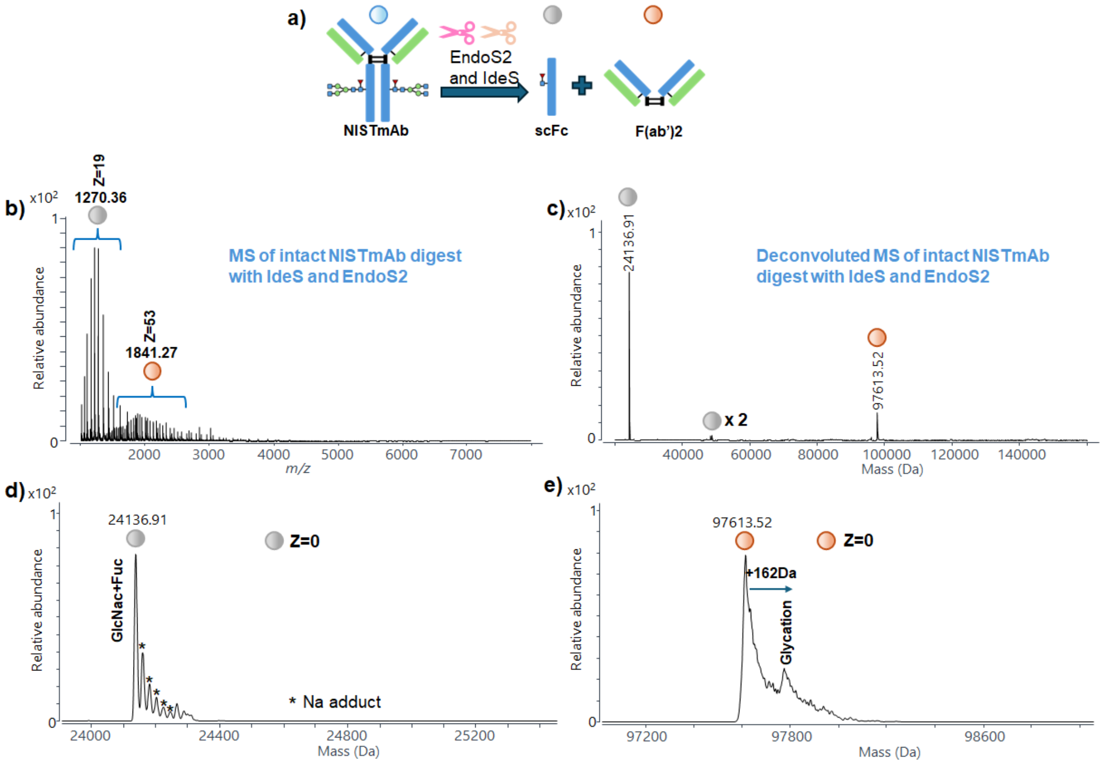


**Figure S5.** (a) Scheme showing digestion of intact NISTmAb with both IdeS and EndoS2; (b) MS spectrum of intact NISTmAb microdroplet digestion with IdeS and EndoS2; (c) deconvoluted MS spectrum of intact NISTmAb microdroplet digestion with IdeS and EndoS2; zoomed-in deconvoluted MS spectra of (d) scFc and (e) F(ab')2 subunits region. Some small adjacent peaks labeled as * in d) were attributed to sodium adduct formation. The glycation peak observed in (e) was probably due to trace glucose present in the sample.

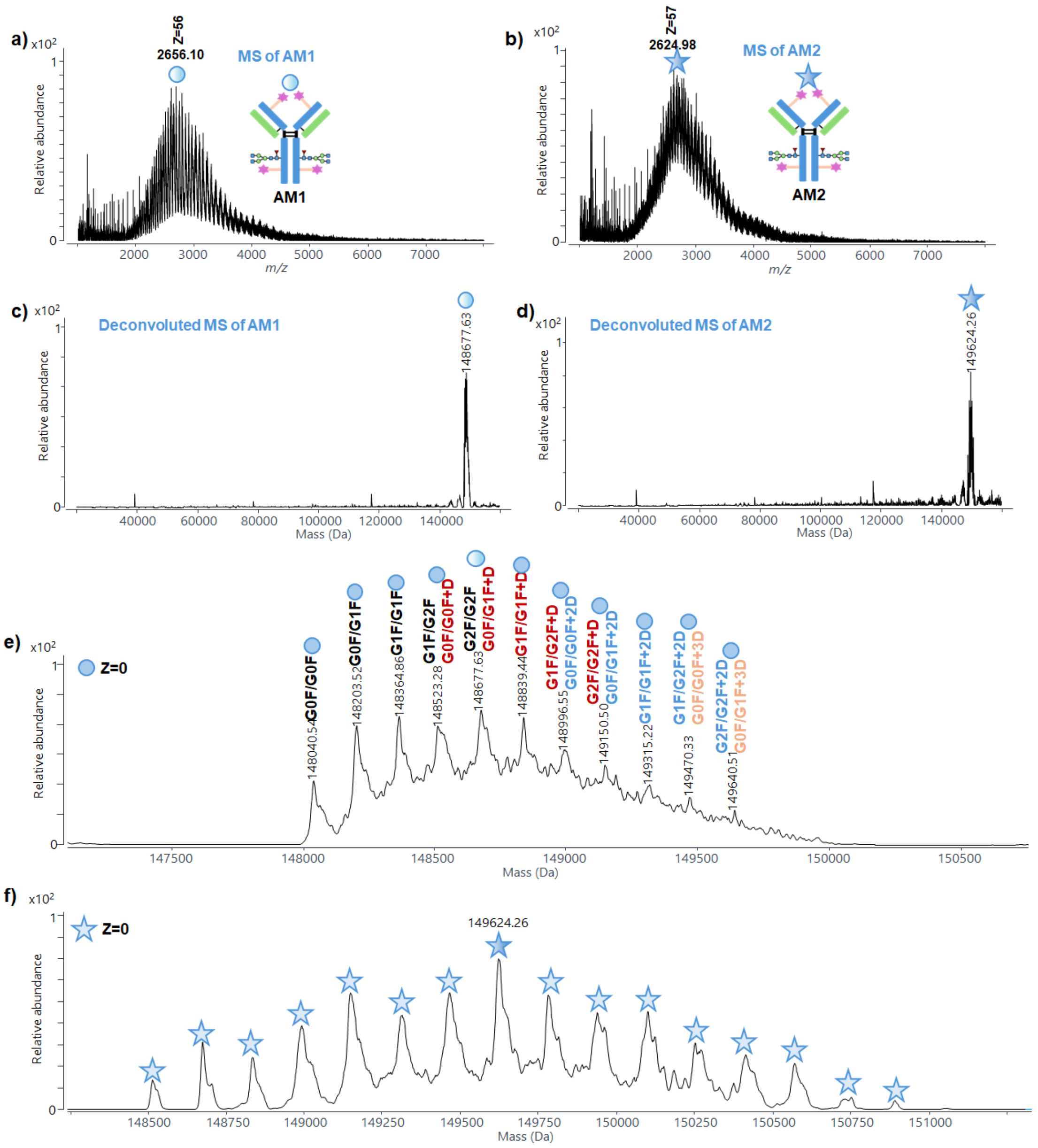


**Figure S6.** Mass spectra of (a) **AM1** and (b) **AM2**; deconvoluted mass spectra of (c) **AM1** and (d) **AM2**; zoomed-in views of the deconvoluted mass spectra of (e) **AM1** and (f) **AM2**. **AM2** exhibited a greater number of peaks, and the lowest observed molecular weight (148516.38 Da) exceeded that of the intact NISTmAb-G0F/G0F species (148038.94 Da; Figure S2c), indicating the absence of unconjugated NISTmAb.

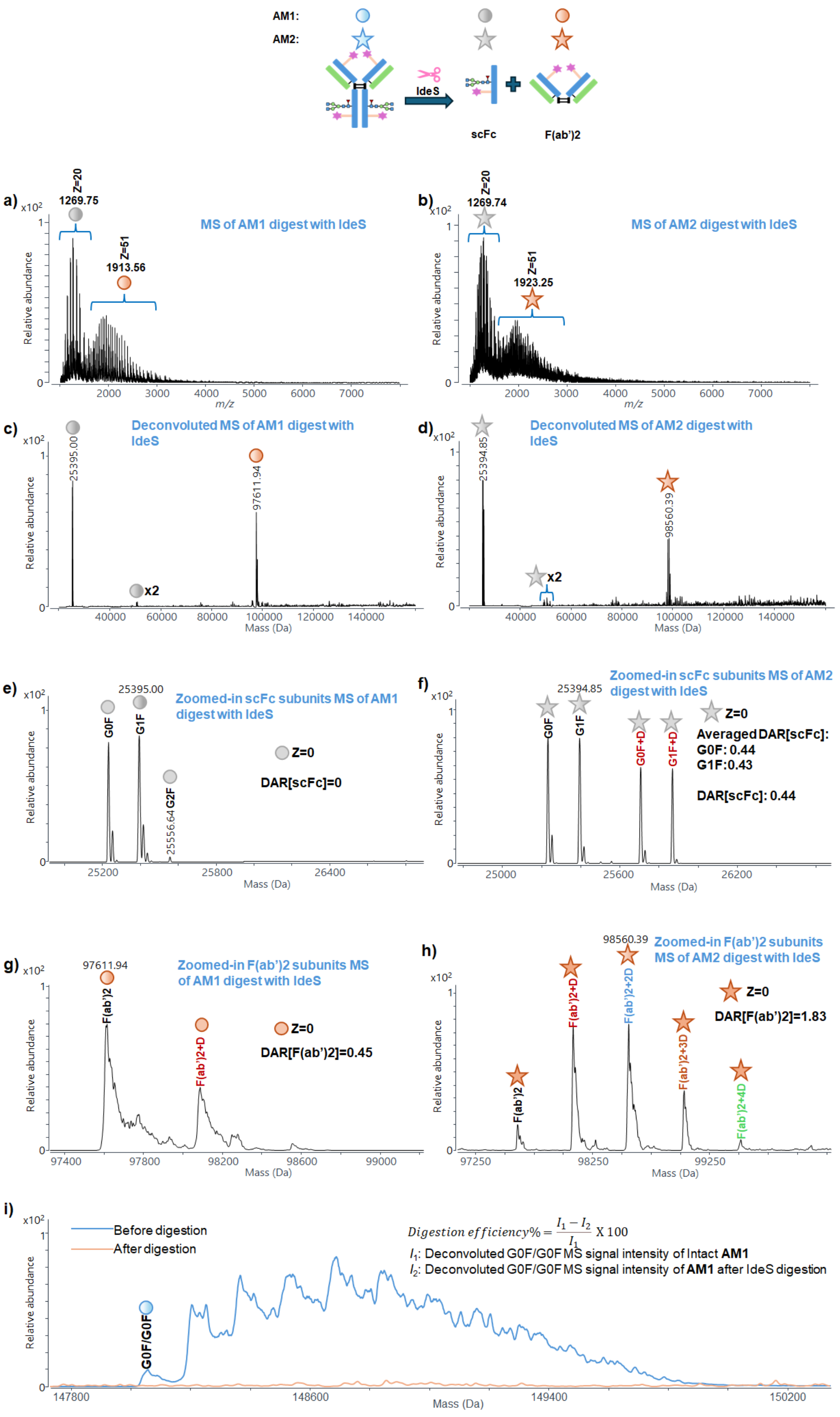


**Figure S7.** Mass spectra of (a) **AM1** and (b) **AM2** after microdroplet digestion with IdeS; deconvoluted mass spectra of (c) **AM1** and (d) **AM2** after microdroplet digestion with IdeS; zoomed-in deconvoluted MS spectra of scFc of (e) **AM1** and (f) **AM2** and F(ab')2 subunits region (g) **AM1** and (h) **AM2.** (i) Example illustrating the calculation of IdeS digestion efficiency using **AM1** as a representative case.

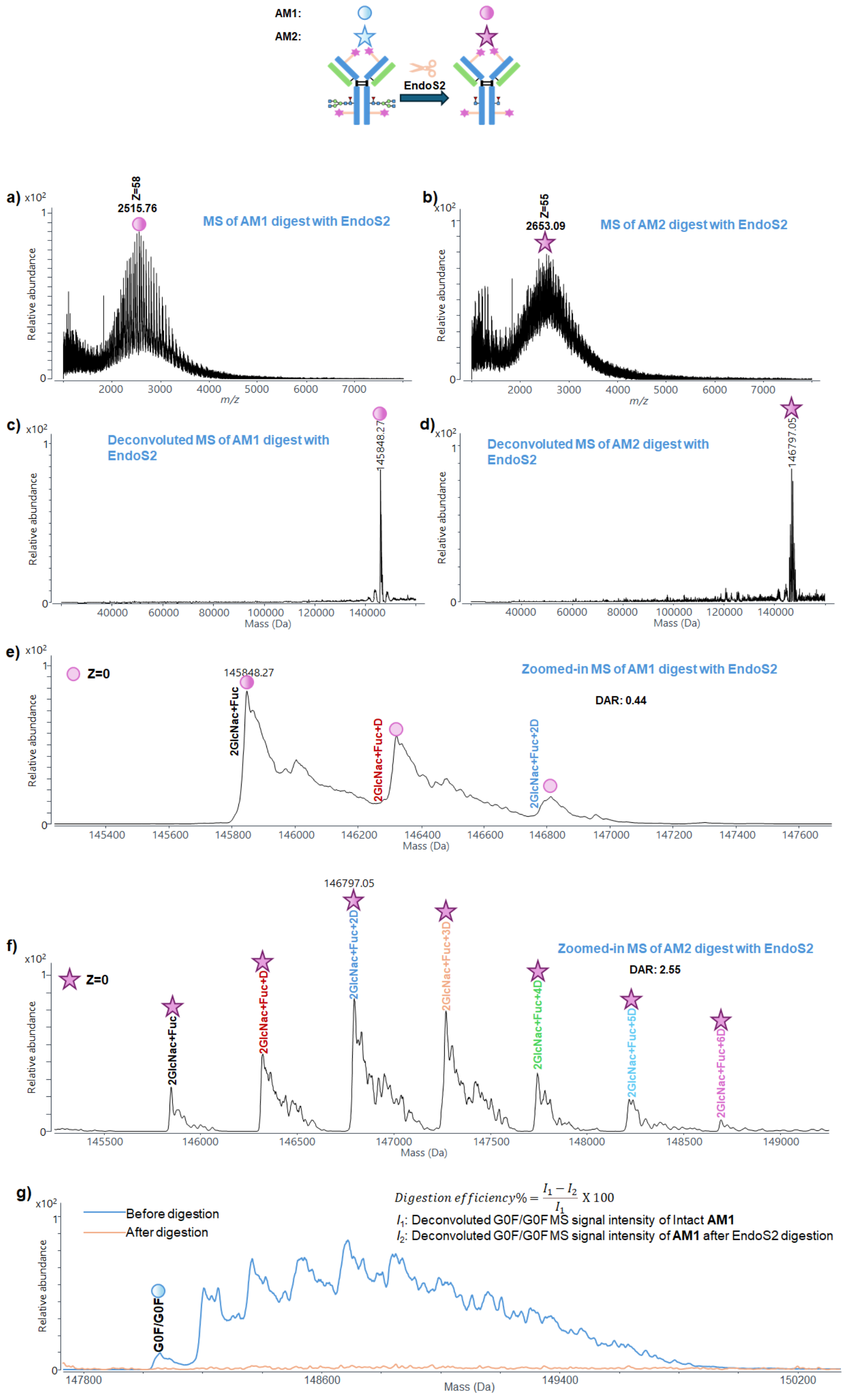


**Figure S8.** Mass spectra of (a) **AM1** and (b) **AM2** after microdroplet digestion with EndoS2; deconvoluted mass spectra of (c) **AM1** and (d) **AM2** after microdroplet digestion with EndoS2; zoomed-in deconvoluted MS spectra of (e) **AM1** and (f) **AM2**. (g) Example illustrating the calculation of EndoS2 digestion efficiency using **AM1** as a representative case.

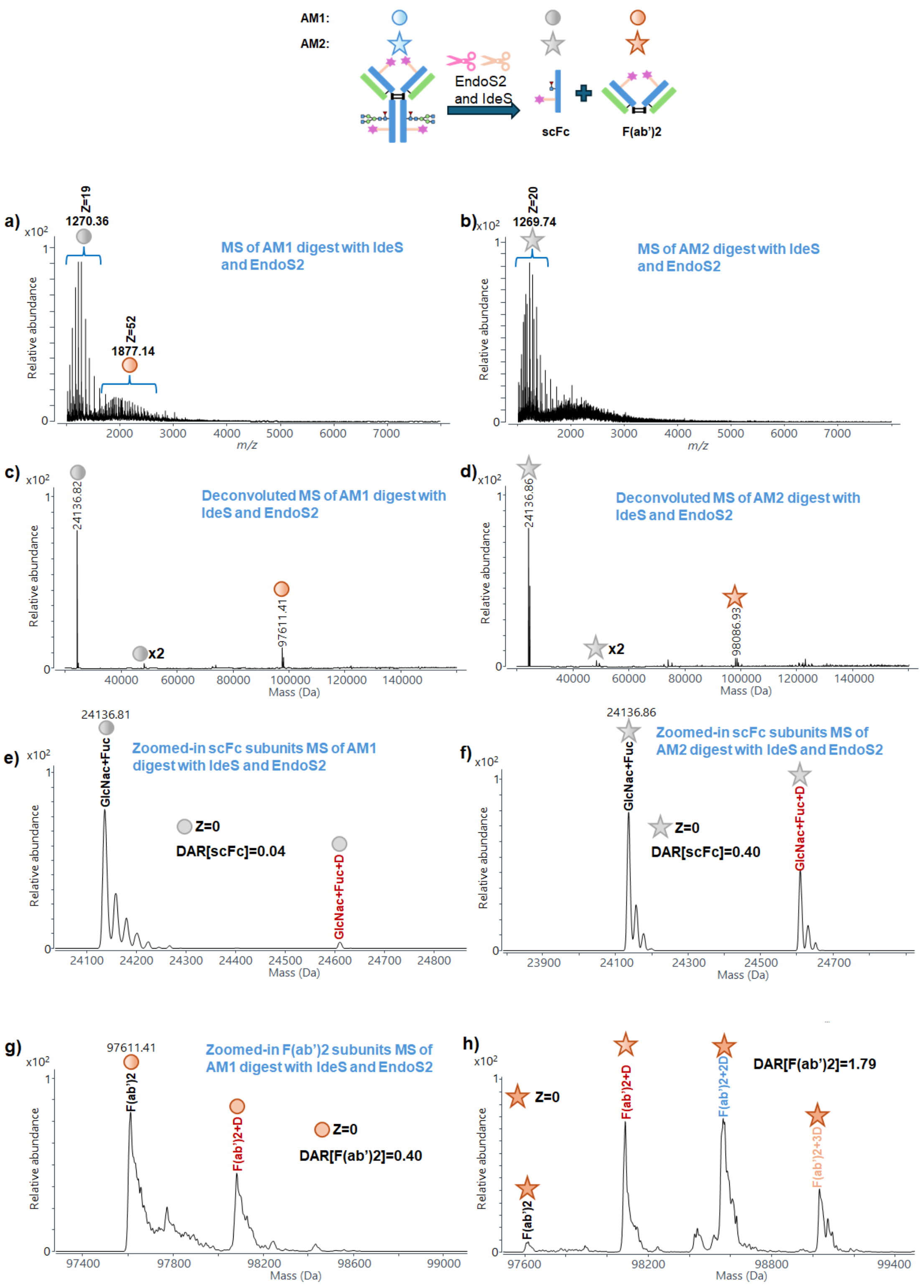


**Figure S9.** Mass spectra of (a) **AM1** and (b) **AM2** after microdroplet digestion with IdeS and EndoS2; deconvoluted mass spectra of (c) **AM1** and (d) **AM2** after microdroplet digestion with IdeS and EndoS2; zoomed-in deconvoluted MS spectra of scFc of (e) **AM1** and (f) **AM2** and F(ab')2 subunits region of (g) **AM1** and (h) **AM2.**

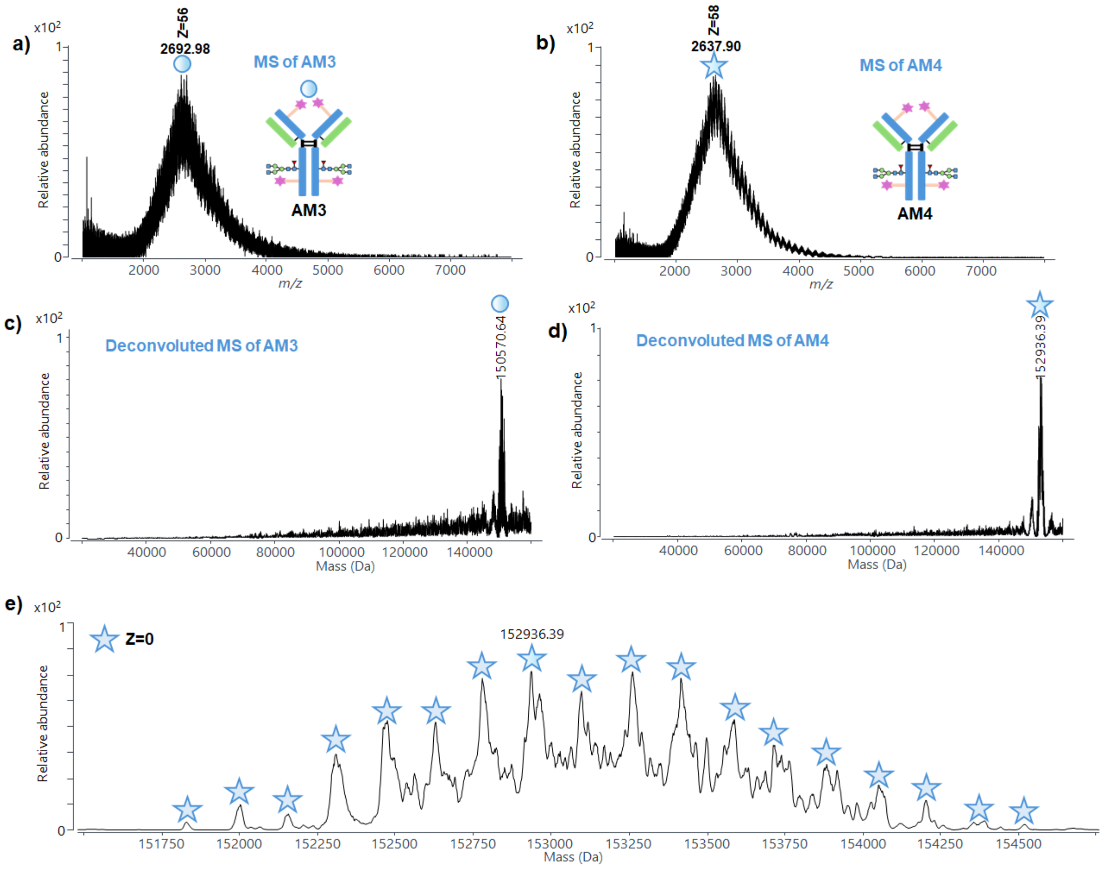


**Figure S10.** Mass spectra of (a) **AM3** and (b) **AM4**; deconvoluted mass spectra of (c) **AM3** and (d) **AM4**; zoomed-in deconvoluted mass spectrum of (e) **AM4.**

**EndoS2 microdroplet digestion of AM3 and AM4**

EndoS2 microdroplet digestion of **AM3** and **AM4** was also performed. The mass spectra of **AM3** and **AM4** after EndoS2 digestion are shown in Figures S11b and S13a, with the corresponding deconvoluted spectra shown in Figure 2d and Figure S13b, respectively. The dominant peaks correspond to molecular weights of 148218.15 Da (**AM3**) and 150593.87 Da (**AM4**), consistent with efficient glycan removal (97.0% and 98.1% respectively, Table S2). Zoomed-in spectra reveal only NISTmAb–GlcNAc+Fuc/GlcNAc+Fuc as the only glycoform conjugated with payload, with **AM3** primarily showing 2–9 NHS-PEG$_4$-Biotin molecule (DAR = 5.37; Figure 2f) and **AM4** showing 8–12 payload species (DAR = 9.65; Figure S13c). These values closely match those obtained from IdeS digestion alone (5.30 and 9.56, respectively).

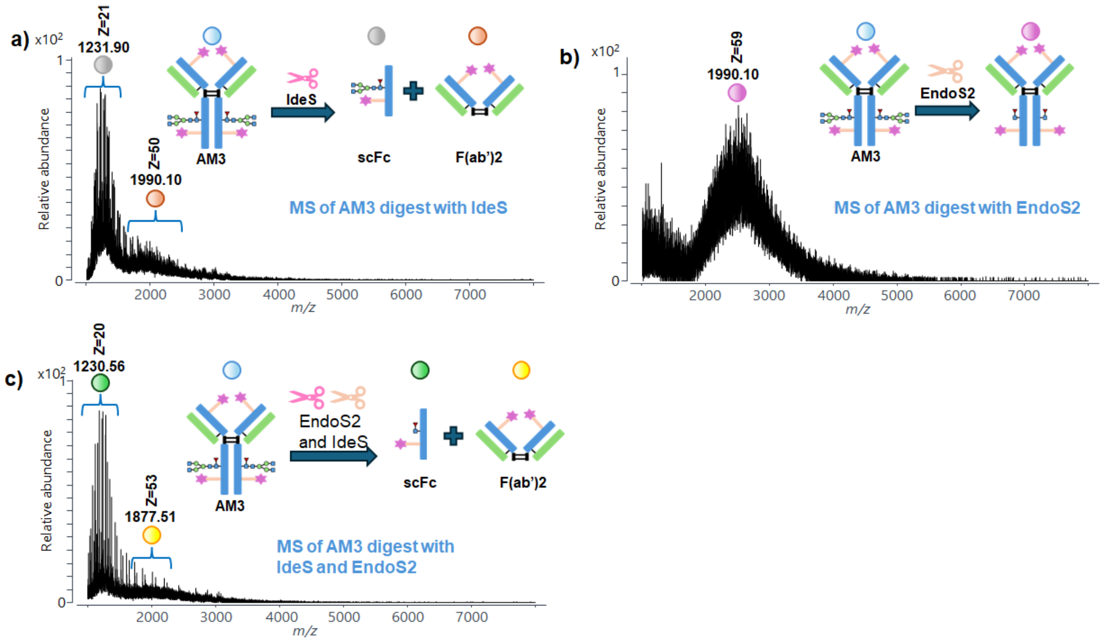


**Figure S11.** Mass spectra of **AM3** after microdroplet digestion with (a) IdeS, (b) EndoS2 and (c) IdeS and EndoS2**.**

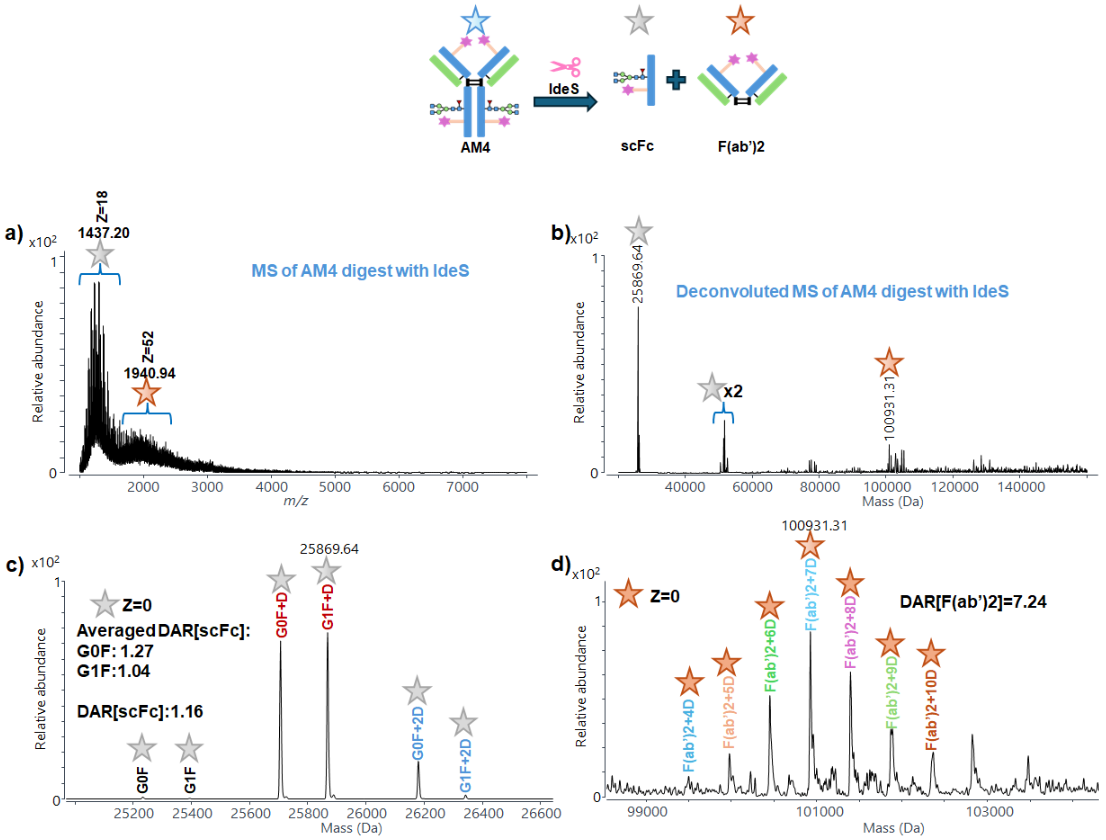


**Figure S12.** (a) Mass spectrum of **AM4** after microdroplet digestion with IdeS; (b) deconvoluted mass spectrum of **AM4** after microdroplet digestion with IdeS; zoomed-in deconvoluted MS spectra of (c) scFc and (d) F(ab')2 subunits region.

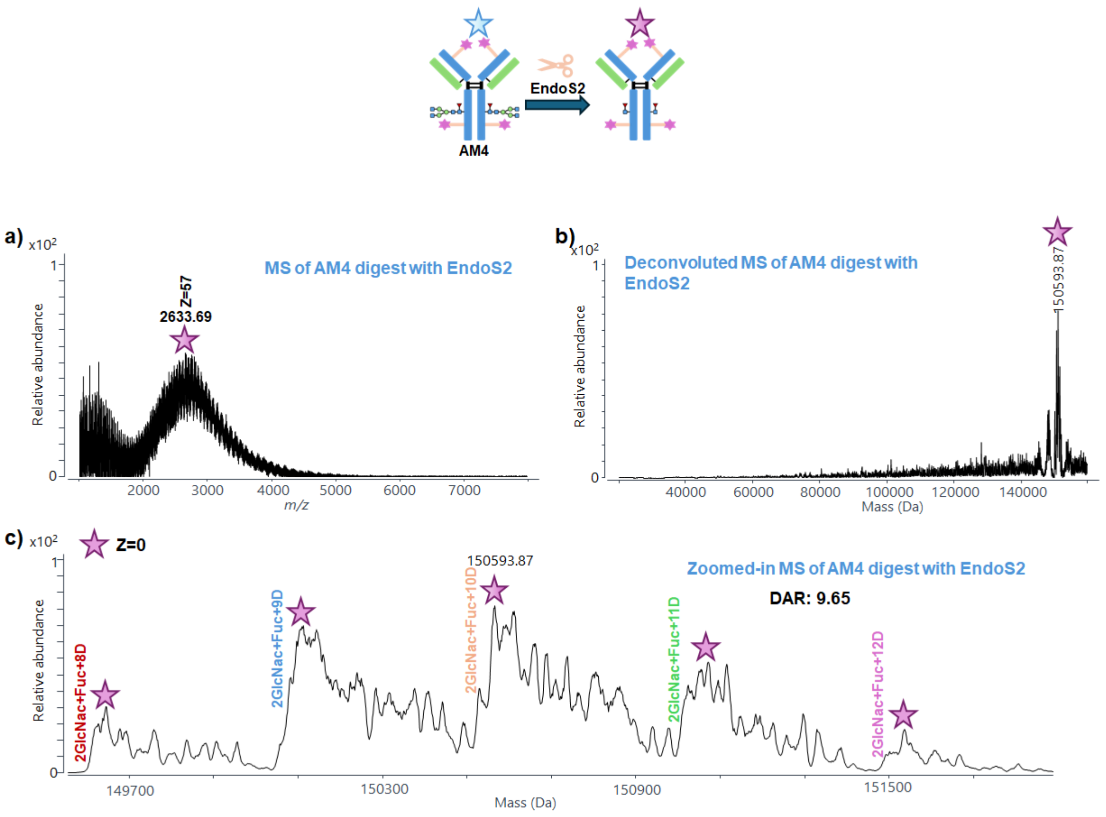


**Figure S13.** (a) Mass spectrum of **AM4** after microdroplet digestion with EndoS2; (b) deconvoluted mass spectrum of **AM4** after microdroplet digestion with EndoS2; (c) zoomed-in deconvoluted mass spectrum of **AM4** after microdroplet digestion with EndoS2.

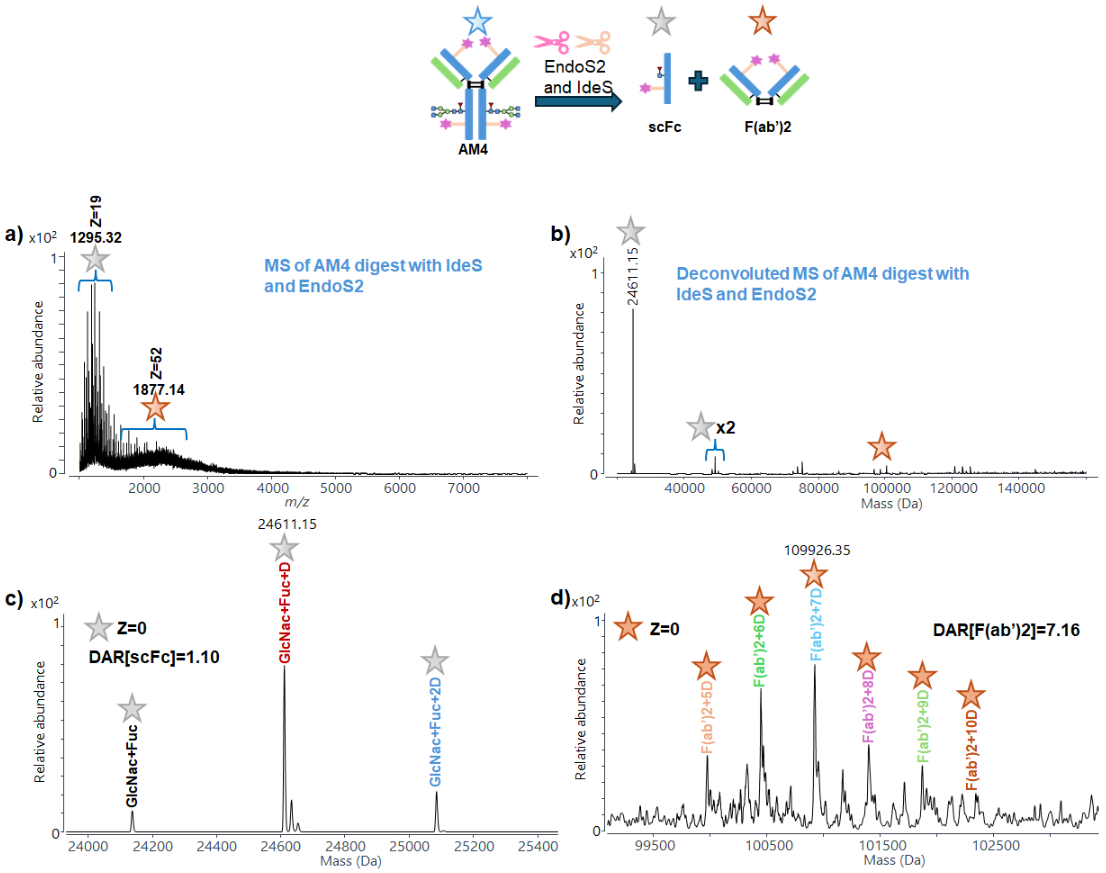


**Figure S14.** (a) Mass spectrum of **AM4** after microdroplet digestion with IdeS and EndoS2; (b) deconvoluted mass spectrum of **AM4** after microdroplet digestion with IdeS and EndoS2; zoomed-in deconvoluted MS spectra of (c) scFc and (d) F(ab')2 subunits region.

**Microdroplet digestion of AM5 and AM6**

Their intact MS spectra, deconvoluted spectra, and zoomed-in deconvolution spectra are shown in Figure S15. Both **AM5** and **AM6** exhibit extensive NHS-PEG$_4$-Biotin distributions convoluted with glycoform distributions (Figures S15e and S15f), similar to those observed for **AM1**–**AM4**. The coexistence of multiple glycoforms and conjugation states makes direct DAR determination from intact MS spectra impractical. Therefore, our enzymatic digestion strategies used for other NISTmAb–ADC mimics were applied to these highly conjugated samples. First, **AM5** and **AM6** were microdroplet digested using IdeS with high efficiency (Table S2). The corresponding MS spectra are shown in Figures S16a and S16b, and the deconvoluted spectra are presented in Figures S16c (**AM5**) and S16d (**AM6**). In both cases, clear scFc signals were observed. After zooming into the scFc region, the glycoforms and their NHS-PEG$_4$-Biotin counterparts could be readily identified, allowing the calculation of DAR[scFc] values of 1.15 for **AM5** (Figure S16e) and 1.75 for **AM6** (Figure S16f). Upon magnification of the F(ab')2 subunit region, comparable DAR values were observed for the two samples, with DAR[F(ab')2] of 9.81 for **AM5** (Figure S16g) and 10.01 for **AM6** (Figure S16h). Based on these subunit-level measurements, the overall DAR values determined by this method were 12.11 for **AM5** and 13.51 for **AM6**. Next, **AM5** and **AM6** were subjected to microdroplet digestion using EndoS2 alone. The resulting MS spectra and deconvoluted results are shown in Figures S17a–d. After zooming (Figures S17e and S17f), antibodies conjugated with a different number of NHS-PEG$_4$-BiotinscFc was observed with a single GlcNAc+Fuc–glycoform, removing interferences for DAR determination from heterogeneous glycoforms. Under these conditions, the DARs could be readily determined, yielding values of 11.84 for **AM5** and 13.36 for **AM6** (Table 1). These results demonstrate that EndoS2 digestion alone is suitable for determining the total DAR of highly conjugated NISTmAb–ADC mimic. Finally, **AM5** and **AM6** were digested using a combination of IdeS and EndoS2. The corresponding MS spectra and deconvoluted results are shown in Figures S18a–d. Nevertheless, scFc signals were significantly enhanced and free from glycoform interference, enabling more accurate determination of DAR[scFc]. After zoomed-in analysis (Figures S18e and S18f), the DAR[scFc] values were determined to be 1.40 for **AM5** and 1.86 for **AM6**, consistent with the results obtained from IdeS-only digestion but with improved accuracy due to the absence of glycoform heterogeneity. Similar to IdeS-only digestion, the F(ab')2 signals remained low, which can be attributed to the combined effects of enhanced scFc ionization efficiency following glycan removal by EndoS2 and severe ionization suppression of heavily conjugated F(ab')2 subunits with NHS-PEG$_4$-Biotin conjugation. Nevertheless, upon zooming in, the signals corresponding to the subunits conjugated with the linker could still be clearly resolved. The resulting DAR[F(ab')2] values were determined to be 9.83 for **AM5** (Figure S18g) and 10.02 for **AM6** (Figure S18h). Using this approach, the calculated overall DAR values were 12.63 for **AM5** and 13.74 for **AM6**, as summarized in Table 1 and Table S1 (%CV <5%). Based on the digestion efficiencies of **AM5** and **AM6** following treatment with IdeS alone, EndoS2 alone, and the combined IdeS/EndoS2 digestion (Table S2), all digestion efficiencies were above 96%. These results indicate that highly conjugated samples did not adversely affect the microdroplet digestion efficiency.

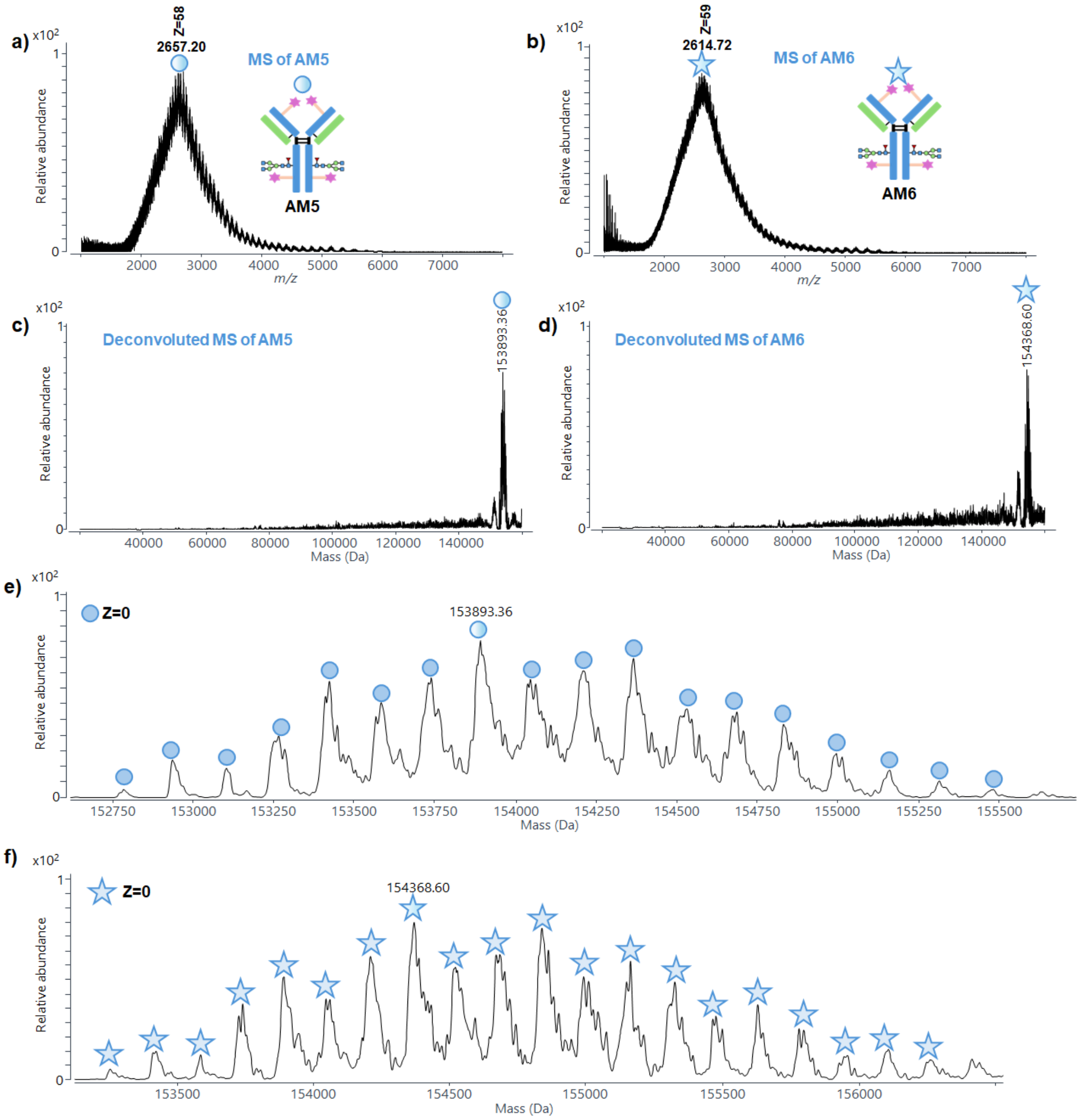


**Figure S15.** Mass spectra of (a) **AM5** and (b) **AM6**; deconvoluted mass spectra of (c) **AM5** and (d) **AM6**; zoomed-in views of the deconvoluted mass spectra of (e) **AM5** and (f) **AM6**.

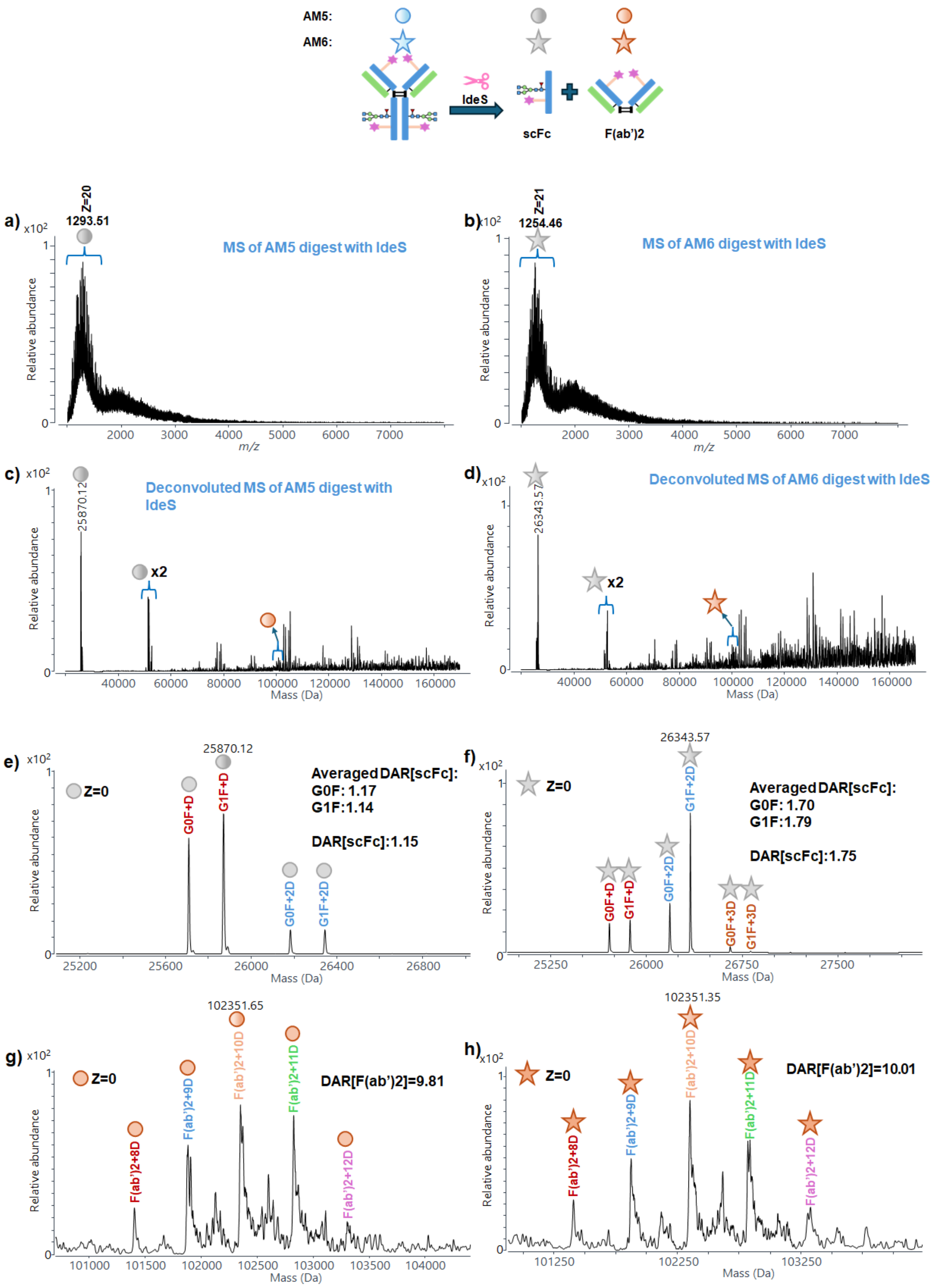


**Figure S16.** Mass spectra of (a) **AM5** and (b) **AM6** after microdroplet digestion with IdeS; deconvoluted mass spectra of (c) **AM5** and (d) **AM6** after microdroplet digestion with IdeS; zoomed-in deconvoluted MS spectra of scFc of (e) **AM5** and (f) **AM6** and F(ab')2 subunits region (g) **AM5** and (h) **AM6**.

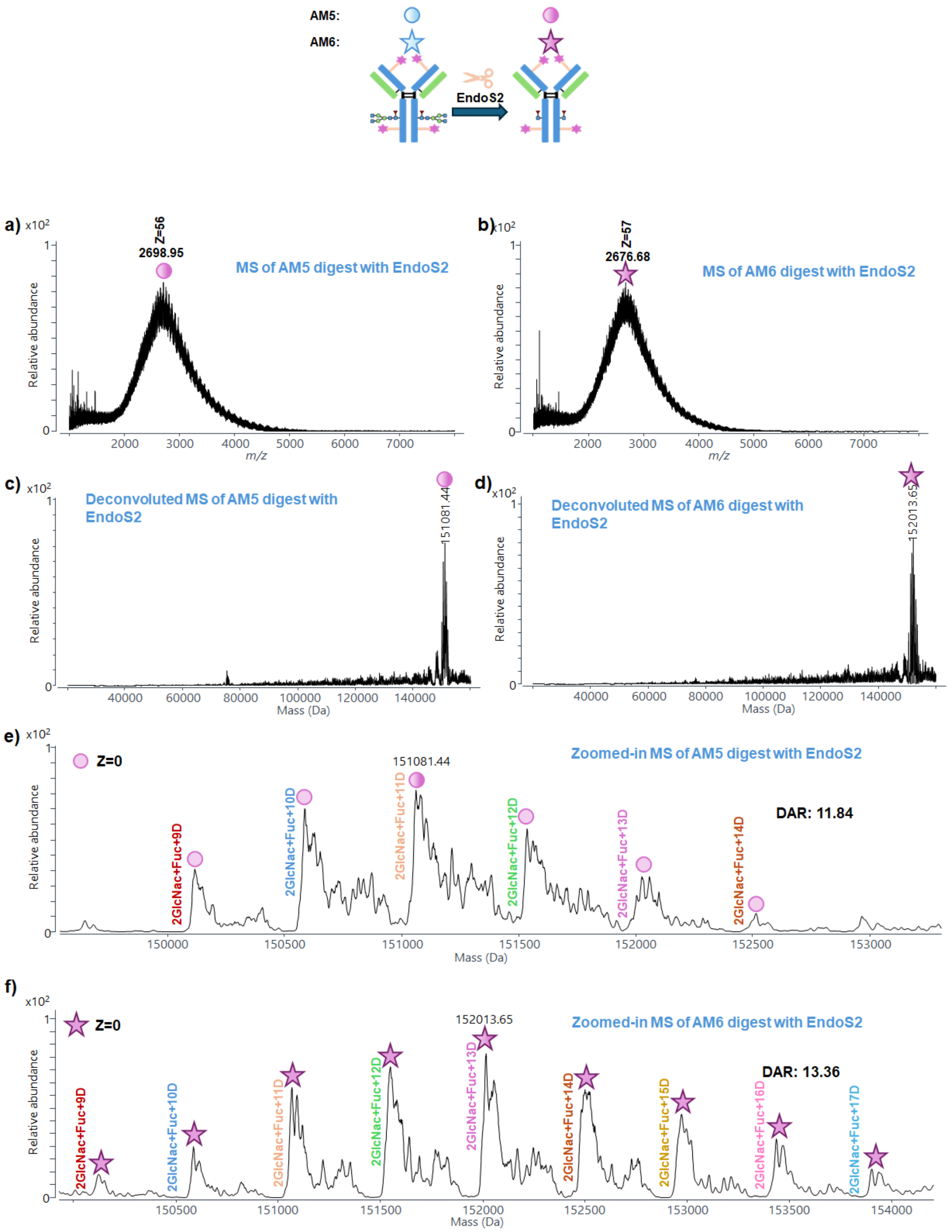


**Figure S17.** Mass spectra of (a) **AM5** and (b) **AM6** after microdroplet digestion with EndoS2; deconvoluted mass spectra of (c) **AM5** and (d) **AM6** after microdroplet digestion with EndoS2; zoomed-in deconvoluted MS spectra of (e) **AM5** and (f) **AM6**.

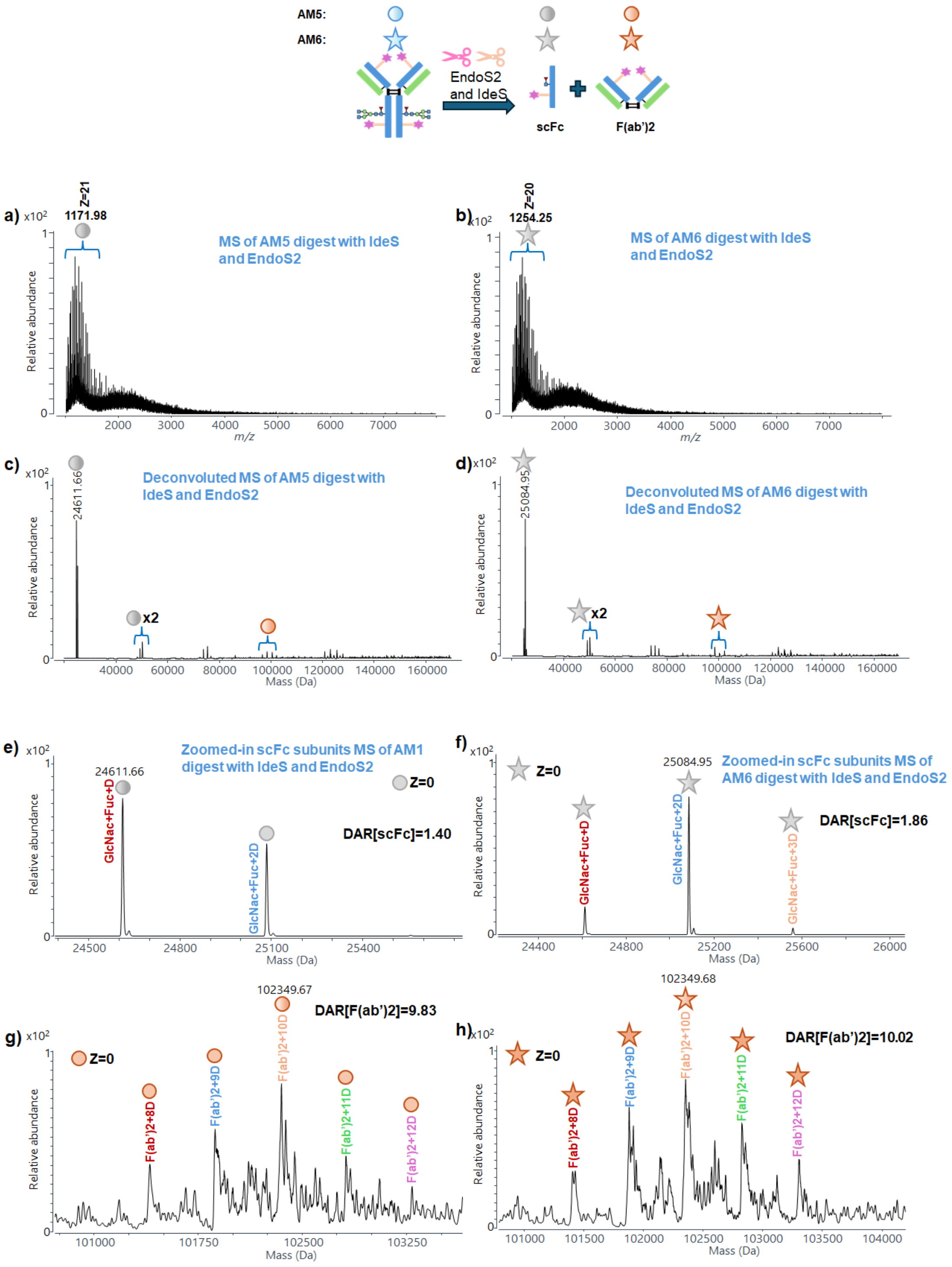


**Figure S18.** Mass spectra of (a) **AM5** and (b) **AM6** after microdroplet digestion with IdeS and EndoS2; deconvoluted mass spectra of (c) **AM5** and (d) **AM6** after microdroplet digestion with IdeS and EndoS2; zoomed-in deconvoluted MS spectra of scFc of (e) **AM5** and (f) **AM6** and F(ab')2 subunits region (g) **AM5** and (h) **AM6**.

### Intact Cetuximab microdroplet digestion

The MS spectrum of intact Cetuximab is shown in Figure S19a. After deconvolution, the main mass peak of intact Cetuximab was observed at 152745.67 Da (Figure S19b). A zoomed-in view of the deconvoluted spectrum (Figure S19c) reveals a significantly larger number of glycoform-related peaks compared to intact NISTmAb (Figure S2c), which are challenging to annotate by intact Cetuximab mass analysis. However, Cetuximab glycoforms in both Fc and Fab regions have been reported at the subunit level in previous studies.[1, 2] Following a workflow analogous to that used for intact NISTmAb, Cetuximab was first subjected to microdroplet digestion with IdeS, as schematically illustrated in Figure S20a. The corresponding MS spectrum and deconvoluted spectrum are shown in Figures S20b and S20c, respectively. Compared with the IdeS-digested intact NISTmAb (Figure S3c), the signal intensity of the F(ab')2 subunit relative to scFc was substantially lower for Cetuximab. This observation can be attributed to the presence of multiple terminal sialic acids on the F(ab')2 region of Cetuximab, which significantly reduces its ionization efficiency. After zooming into the scFc and F(ab')2 regions (Figures S20d and S20e), it was observed that the scFc subunit of Cetuximab (Figure S20d) is nearly identical to that of NISTmAb (Figure S3d). In contrast, the F(ab')2 region exhibits multiple peaks (Figure S20e), which is markedly different from the single dominant F(ab')2 peak observed for NISTmAb (Figure S3e). These additional peaks arise from glycoform heterogeneity associated with glycans located on the Fab region. The peaks corresponding to 1-4 N-acetylneuraminic acids (NeuAc) conjugated with G2F/G2F glycoform were also observed. NeuAc is the most abundant sialic acid in humans and serves as a terminal monosaccharide in antibody-associated glycan structures, contributing further to spectral complexity. Cetuximab was subsequently subjected to microdroplet digestion with EndoS2, and the results are shown in Figure S21. Unlike NISTmAb, for which EndoS2 digestion yielded predominantly a single NISTmAb – GlcNAc+Fuc/GlcNAc+Fuc glycoform, the EndoS2-digested Cetuximab spectrum still exhibited numerous glycoform-related peaks (Figure S21d). This is because EndoS2 selectively cleaves Fc glycans but does not cleave F(ab')2 glycans contributing to substantial heterogeneity due to terminal sialic acids. To further simplify the system, Cetuximab was subjected to a one-pot microdroplet digestion with a combination of IdeS and EndoS2. As illustrated in Figure S22a, IdeS cleaves Cetuximab into scFc and F(ab')2 subunits, while EndoS2 simultaneously removes Fc-associated glycans. The resulting MS spectrum and deconvoluted spectrum are shown in Figures S22b and S22c, respectively. Under these conditions, the signal intensity of the scFc subunit was further enhanced due to glycan removal, with a concomitant loss of F(ab')2 signal. We were unable to detect F(ab')2 by filtering the *m/z* regions of F(ab')2 from the raw spectrum. This behavior is in stark contrast to that observed for NISTmAb (Figures S5c and S4e). A zoomed-in view of the scFc region (Figure S22d) reveals scFc with a single GlcNAc+Fuc glycoform, consistent with the results obtained for NISTmAb (Figure S5d). These experiments clearly demonstrate that sialic acid glycoforms on the F(ab')2 region exert a strong ion suppression effect on the ionization of F(ab')2. To tackle this problem, another enzyme, EndoF3, was introduced. EndoF3 is an endoglycosidase that cleaves within the chitobiose core of N-linked, fucosylated biantennary and triantennary complex oligosaccharides on glycoproteins. Previous studies have demonstrated that EndoF3 in-solution digestion is capable of efficiently removing glycan heterogeneity to produce a F(ab')2 with a GlcNAc+Fuc/GlcNAc+Fuc glycoform of Cetuximab, making it well suited for this application.[3] Therefore, Cetuximab was first subjected to microdroplet digestion using EndoF3 alone, as schematically illustrated in Figure S23a. The MS spectrum of Cetuximab after EndoF3 microdroplet digestion and its corresponding deconvoluted spectrum are shown in Figures S23b and S23c, respectively. As shown in Figure S23c, the highest mass peak of EndoF3-digested Cetuximab appears at 149049.95 Da, which is substantially lower than that of intact Cetuximab (152745.67 Da; Figure S19b). This significant mass decrease confirms that EndoF3 can efficiently digest Cetuximab under microdroplet conditions. However, upon zooming into the peak region (Figure S23d), multiple glycoform-related peaks remain observable, indicating that a portion of the Fc-region glycans are not fully cleaved by EndoF3 alone.

Subsequently, intact Cetuximab was co-digested with EndoF3 and IdeS, as illustrated in Figure S24a. Under these conditions, partial removal of Fc glycans by EndoF3 results in the formation of two distinct scFc species with a single GlcNAc+Fuc glycoform and glycosylated scFc with G0F and G1F glycoforms—represented by the orange and gray circles in the scheme, respectively. The corresponding MS spectrum and deconvoluted spectrum are shown in Figures S24b and S24c. Two clearly resolved scFc peaks, the coexistence of GlcNAc+Fuc species together with G0F and G1F glycoforms, are observed, corresponding to deglycosylated and intact scFc, respectively, providing direct evidence that EndoF3 removes a fraction of Fc-associated glycans (Figure S24d). To the best of our knowledge, our result represents the first experimental evidence demonstrating partial Fc deglycosylation by EndoF3 under microdroplet reaction conditions. Notably, because glycans on the F(ab')2 region were effectively removed by microdroplet digestion with EndoF3 with 97.2% efficiency (the calculation of the digestion efficiency used the intensity of the most abundant peak of intact Cetuximab which was compared with that of the corresponding peak after EndoF3 digestion; one calculation example is shown in Figure S31d for the **AM7** microdroplet digestion with EndoF3), the ionization efficiency of the F(ab')2 subunit was significantly enhanced. A zoomed-in view shows the F(ab')2 with a single GlcNAc+Fuc glycoform (Figure S24e), confirming effective deglycosylation of this subunit. These results demonstrate that the combined use of EndoF3 and IdeS is highly effective in enhancing the relative signal intensity of F(ab')2. Based on the results obtained from NISTmAb–ADC mimics, complete removal of glycan heterogeneity greatly facilitates accurate DAR determination. Accordingly, Cetuximab was further digested using a combination of EndoS2 and EndoF3. The results are shown in Figure S25. When both enzymes were applied, glycans on both the scFc and F(ab')2 subunits were efficiently removed, yielding a single dominant peak corresponding to 2GlcNAc+Fuc/2GlcNAc+Fuc Cetuximab (146676.19 Da). This spectrum closely resembles that obtained for NISTmAb following complete deglycosylation by EndoS2 alone (Figure S4). Finally, intact Cetuximab was subjected to one pot-microdroplet digestion using combination of IdeS, EndoS2, and EndoF3, as illustrated in Figure S26a.

The corresponding MS spectrum and deconvoluted spectrum are shown in Figures S26b and S26c, respectively. Under these conditions, fully cleaved scFc-GlcNAc+Fuc glycoform and F(ab')2 2GlcNAc+Fuc subunits are obtained simultaneously. Nevertheless, both scFc and F(ab')2 contain only GlcNAc+Fuc glycoform peaks (Figures S26d and S26e), confirming the microdroplet reaction efficiency of 96.7% (Table S3). Collectively, these results demonstrate that IdeS, EndoS2, and EndoF3 microdroplet digestion could occur simultaneously to achieve complete and controllable deglycosylation and Fc cleavage at G/G of Cetuximab, thereby facilitating robust subunit-level analysis of highly glycosylated ADCs.

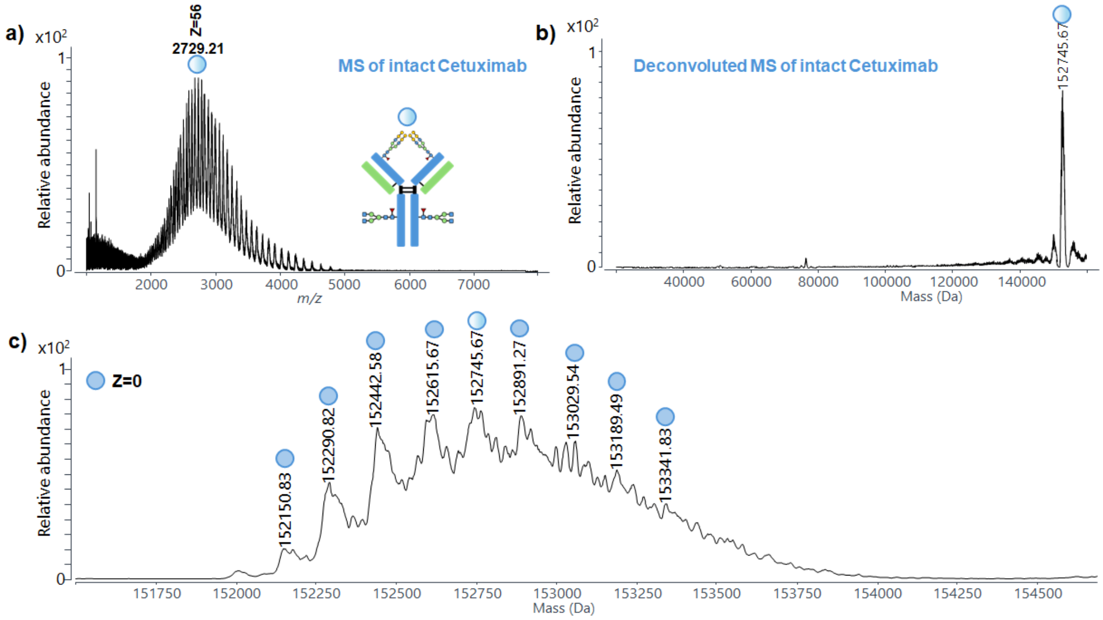


**Figure S19.** (a) Mass spectrum of intact Cetuximab; (b) deconvoluted mass spectrum of intact Cetuximab; (c) zoomed-in deconvoluted mass spectrum in the range of 151750–154500 Da.

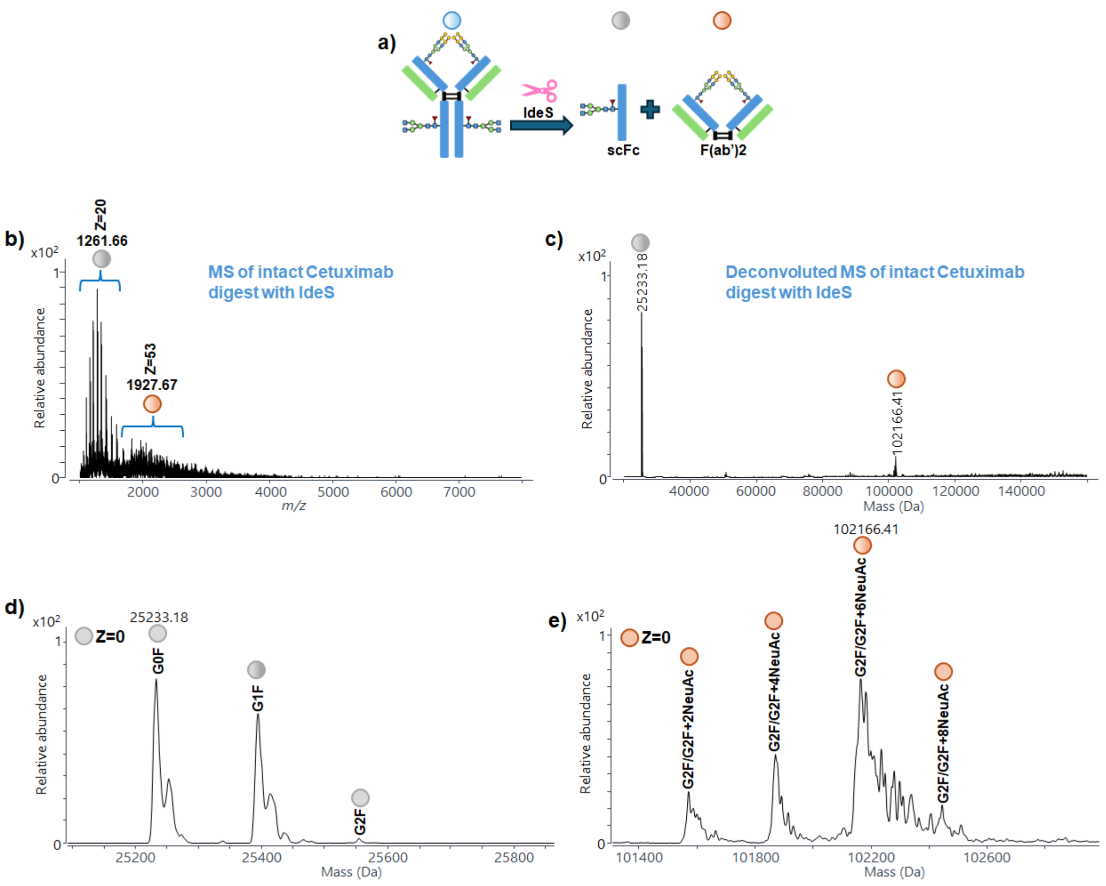


**Figure S20.** (a) Scheme showing digestion of Cetuximab with IdeS; (b) MS spectrum of Cetuximab microdroplet digestion with IdeS; (c) deconvoluted MS spectrum of Cetuximab microdroplet digestion with IdeS; zoomed-in deconvoluted MS spectra of (d) scFc and (e) F(ab')2 subunits region.

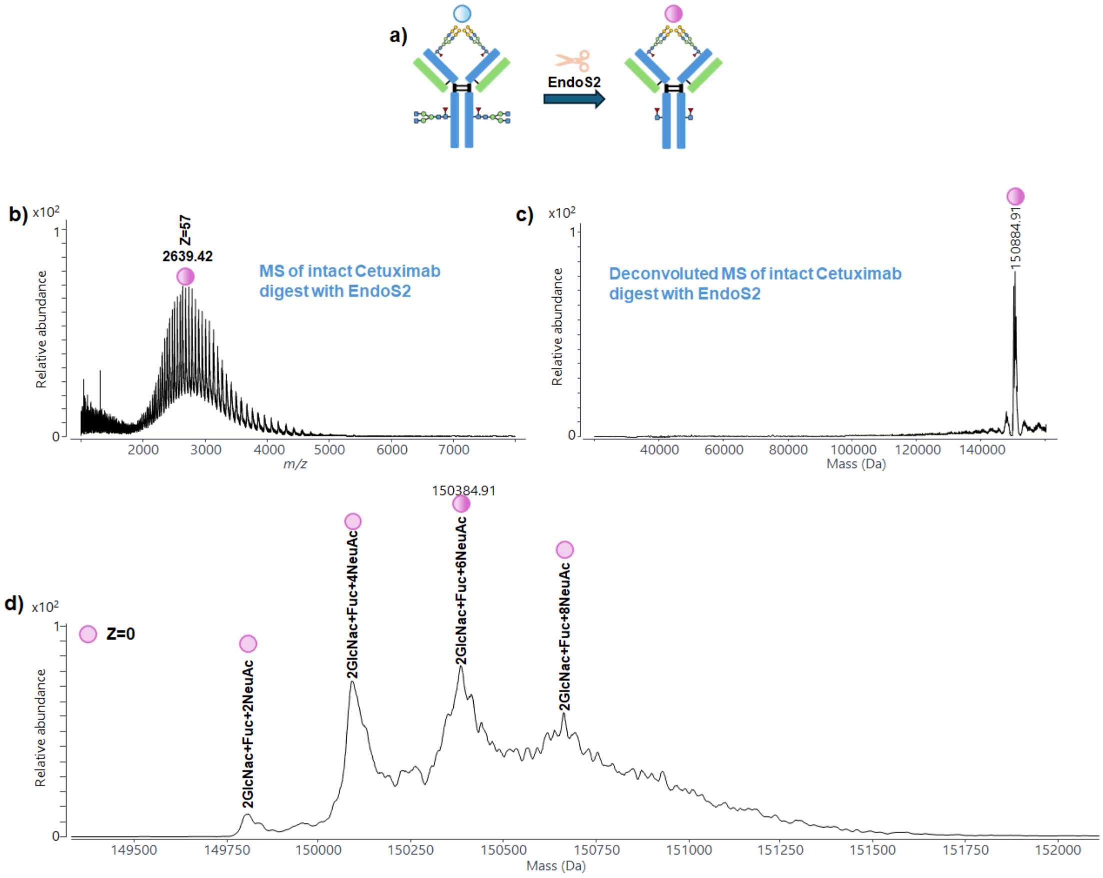


**Figure S21.** (a) Scheme showing digestion of intact Cetuximab with EndoS2; (b) MS spectrum of Cetuximab microdroplet digestion with EndoS2; (c) deconvoluted MS spectrum of Cetuximab microdroplet digestion with EndoS2; (d) zoomed-in deconvoluted MS spectrum of Cetuximab microdroplet digestion with EndoS2.

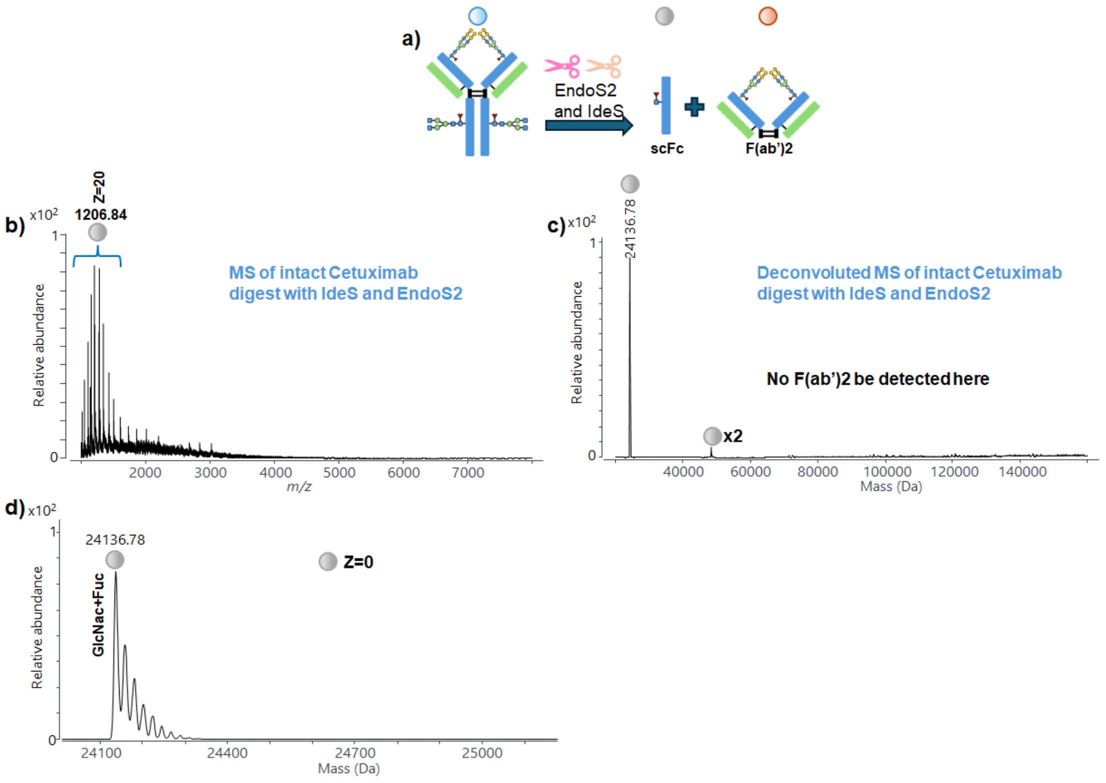


**Figure S22.** (a) Scheme showing digestion of intact Cetuximab with IdeS and EndoS2; (b) MS spectrum of Cetuximab microdroplet digestion with IdeS and EndoS2; (c) deconvoluted MS spectrum of Cetuximab microdroplet digestion with IdeS and EndoS2; (d) zoomed-in deconvoluted MS spectrum of scFc subunits region.

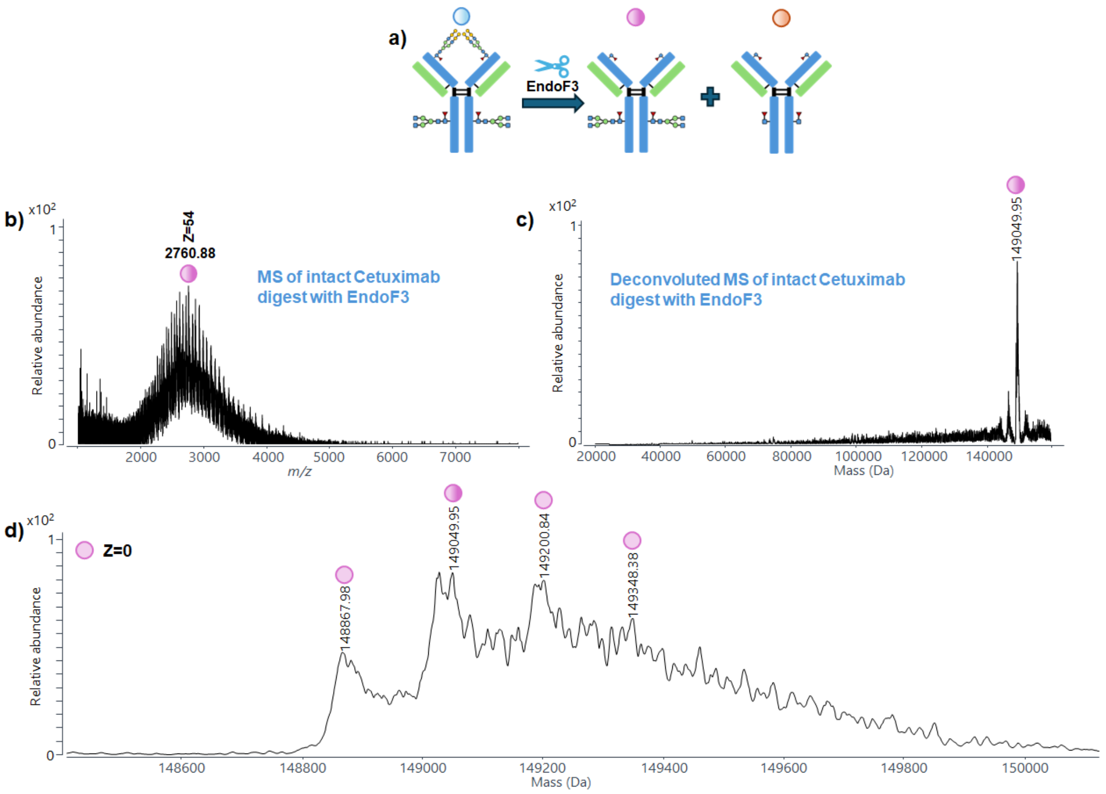


**Figure S23.** (a) Scheme showing the digestion of intact Cetuximab with EndoF3; (b) MS spectrum of Cetuximab microdroplet digestion with EndoF3; (c) deconvoluted MS spectrum of Cetuximab microdroplet digestion with EndoF3; (d) zoomed-in deconvoluted MS spectrum of Cetuximab microdroplet digestion with EndoF3.

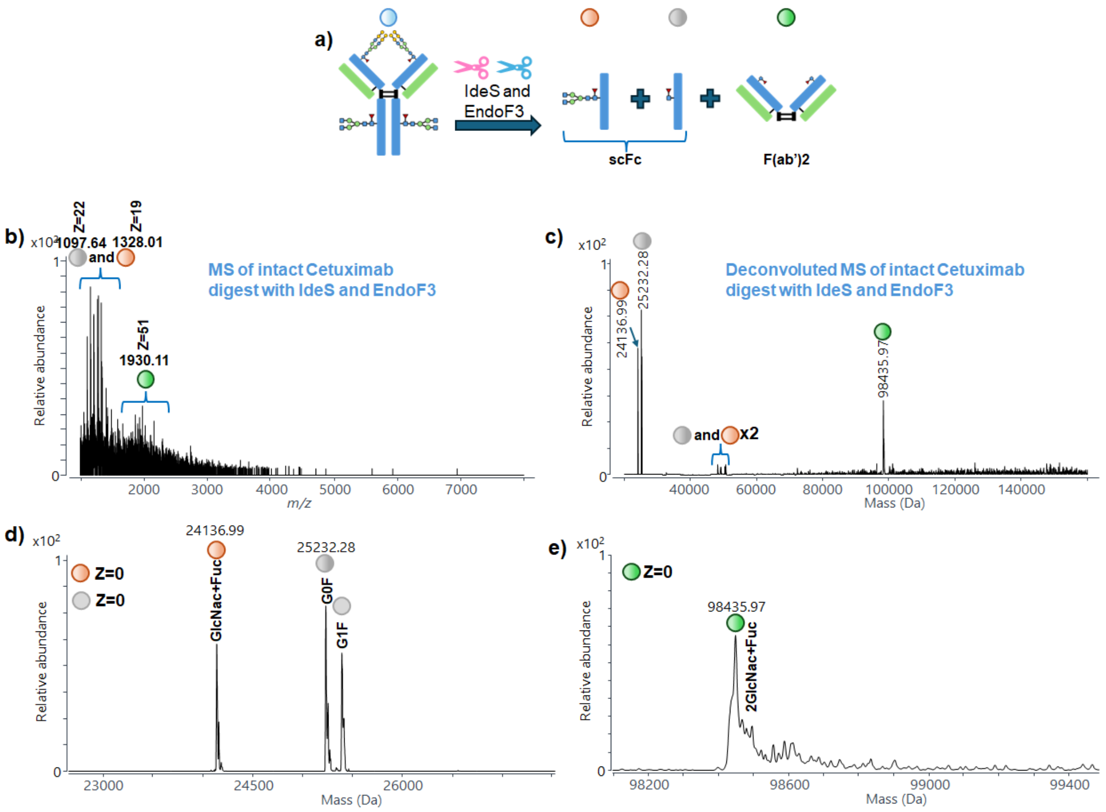


**Figure S24.** (a) Scheme showing digestion of intact Cetuximab with IdeS and EndoF3; (b) MS spectrum of Cetuximab microdroplet digestion with IdeS and EndoF3; (c) deconvoluted MS spectrum of Cetuximab microdroplet digestion with IdeS and EndoF3; zoomed-in deconvoluted MS spectra of (d) scFc and (e) F(ab')2 subunits region.

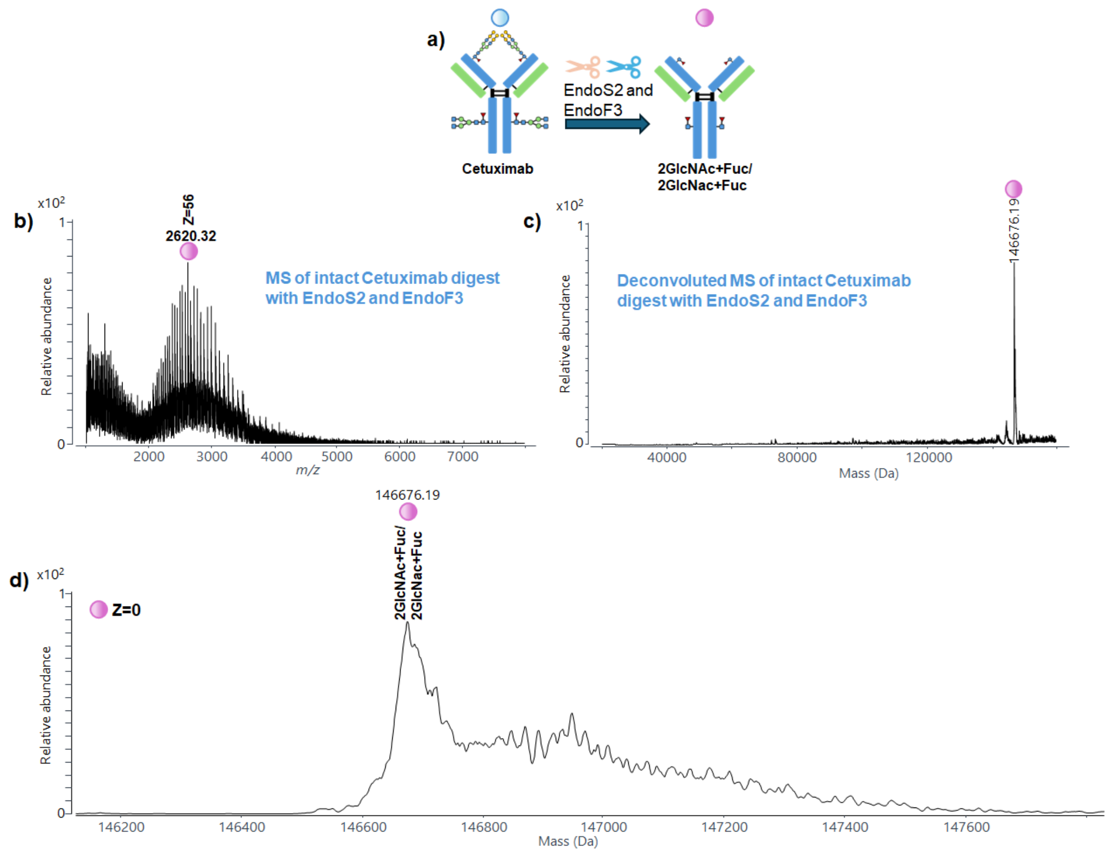


**Figure S25.** (a) Scheme showing digestion of intact Cetuximab with EndoS2 and EndoF3; (b) MS spectrum of Cetuximab microdroplet digestion with EndoS2 and EndoF3; (c) deconvoluted MS spectrum of Cetuximab microdroplet digestion with EndoS2 and EndoF3; (d) zoomed-in deconvoluted MS spectra of Cetuximab microdroplet digestion with EndoS2 and EndoF3.

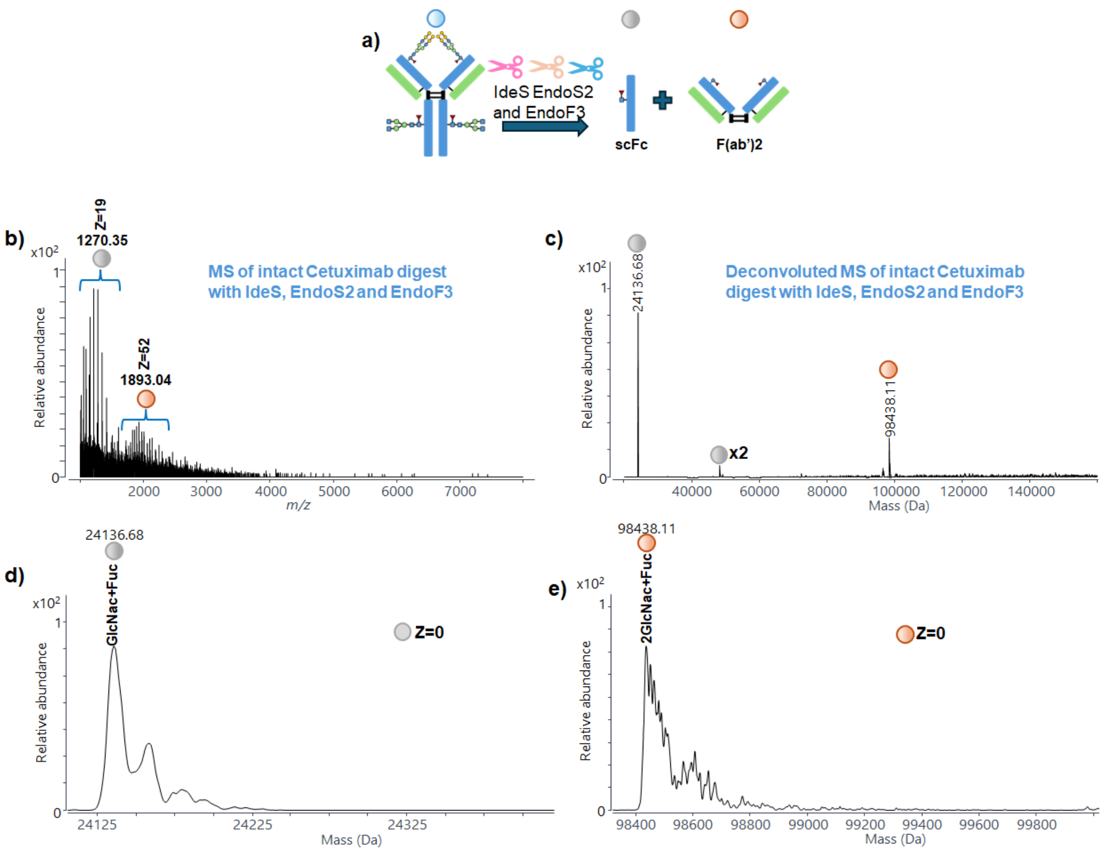


**Figure S26.** (a) Scheme showing digestion of Cetuximab with IdeS, EndoS2 and EndoF3; (b) MS spectrum of Cetuximab microdroplet digestion with IdeS, EndoS2 and EndoF3; (c) deconvoluted MS spectrum of Cetuximab microdroplet digestion with IdeS, EndoS2 and EndoF3; zoomed-in deconvoluted MS spectra of (d) scFc and (e) F(ab')2 subunits region.

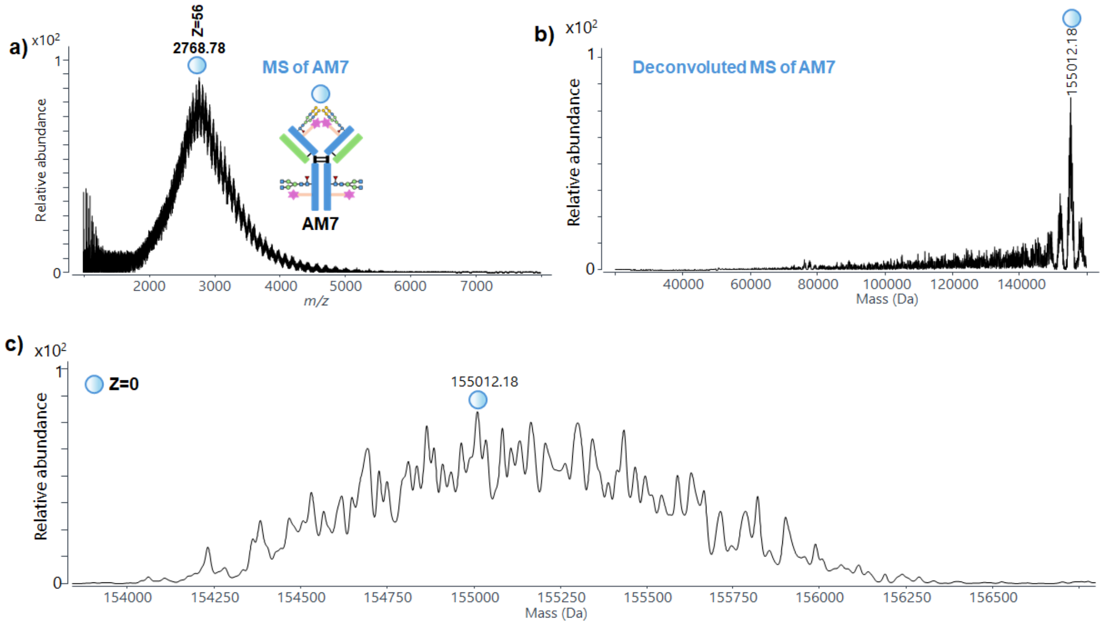


**Figure S27.** (a) Mass spectrum of **AM7**; (b) deconvoluted mass spectrum of **AM7**; (c) zoomed-in view of the deconvoluted mass spectrum of **AM7**.

**AM7 microdroplet digestion results**

As a result, direct DAR determination from the intact spectrum is not feasible. Microdroplet digestion with IdeS alone generated predominantly scFc signals, while F(ab')2 signals were undetectable even by filtering the *m/z* regions of F(ab')2 from the raw spectrum (Figures S28a – c). This observation is attributed to the combined suppression of ionization efficiency caused by Fab-region glycans and extensive payload conjugation. Analysis of the zoomed-in scFc region (Figure S28c) revealed multiple glycoforms and linker-conjugated species, from which a DAR[scFc] of 0.72 was calculated. Subsequent digestion with EndoS2 resulted in partial mass reduction (152930.18 Da, Figure S29b), consistent with Fc glycan removal; however, the zoomed-in spectrum (Figure S29c) remained highly complex due to persistent Fab glycosylation and linker heterogeneity, precluding reliable DAR assignment. Combined IdeS and EndoS2 digestion produced deglycosylated scFc signals (Figure S30 and Figures 3a-c), enabling calculation of a DAR[scFc] of 0.59, but F(ab')2 signals remained undetectable. Thus, none of these three approaches alone enabled the determination of the total DAR for **AM7**. Given the strong suppression of F(ab')2 ionization by Fab glycans, EndoF3, an enzyme capable of cleaving Fab-region glycans as mentioned above, was introduced (the MS spectrum is shown in Figure S31a). Digestion with EndoF3 alone reduced the molecular mass to 151734.17 Da (Figure S31b), confirming enzymatic activity; however, the deconvoluted spectrum remained too complex for DAR analysis (Figure S31c).

To further validate this result, **AM7** was digested with EndoS2 and EndoF3 to remove glycans from both Fc and F(ab')2 regions. The resulting deconvoluted spectrum (Figures S33a – c) showed a main peak at 149086.62 Da, confirming complete deglycosylation. The DAR calculated from this spectrum was 5.19, in close agreement with the value obtained from IdeS and EndoF3 digestion (Figure 32a and Figure 3d-g). Finally, **AM7** was subjected to a one-pot microdroplet digestion with a mixture of (IdeS, EndoS2, and EndoF3) that was premixed in the autosampler and introduced into the MS via the AJS source to enable simultaneous ultrafast microdroplet digestion (<1 ms), as illustrated in Figure 3h. The resulting MS and deconvoluted spectra (Figures 3i–j) showed deglycosylated scFc signals, from which a DAR[scFc] of 0.61 was calculated. This value closely matches that obtained from IdeS and EndoS2 digestion (0.59), while methods retaining Fc glycans yielded larger deviations. Consequently, Fc deglycosylation by EndoS2 is essential for accurate scFc DAR determination in Cetuximab–ADC mimics. Because F(ab')2 signals remained weak under full three-enzyme digestion conditions, DAR[F(ab')2] was instead determined using IdeS and EndoF3 digestion.

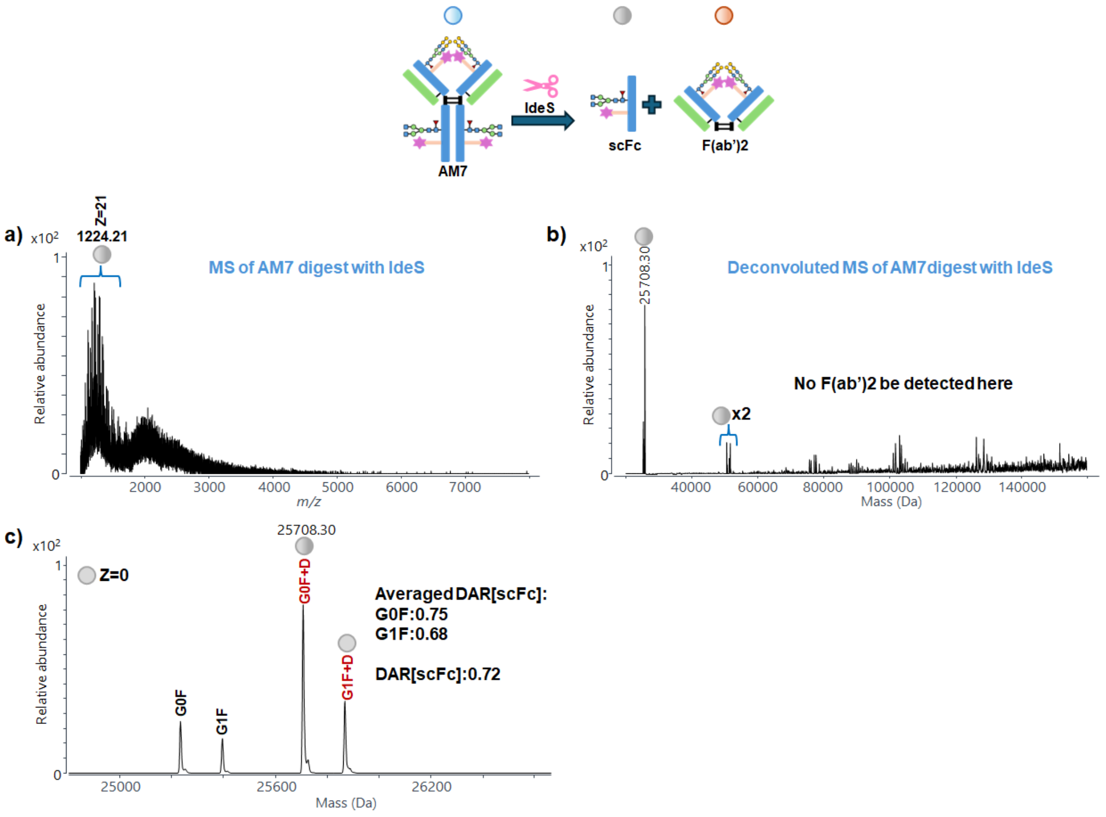


**Figure S28.** (a) MS spectrum of **AM7** microdroplet digestion with IdeS; (b) deconvoluted MS spectrum of **AM7** microdroplet digestion with IdeS; (c) zoomed-in deconvoluted MS spectra of scFc.

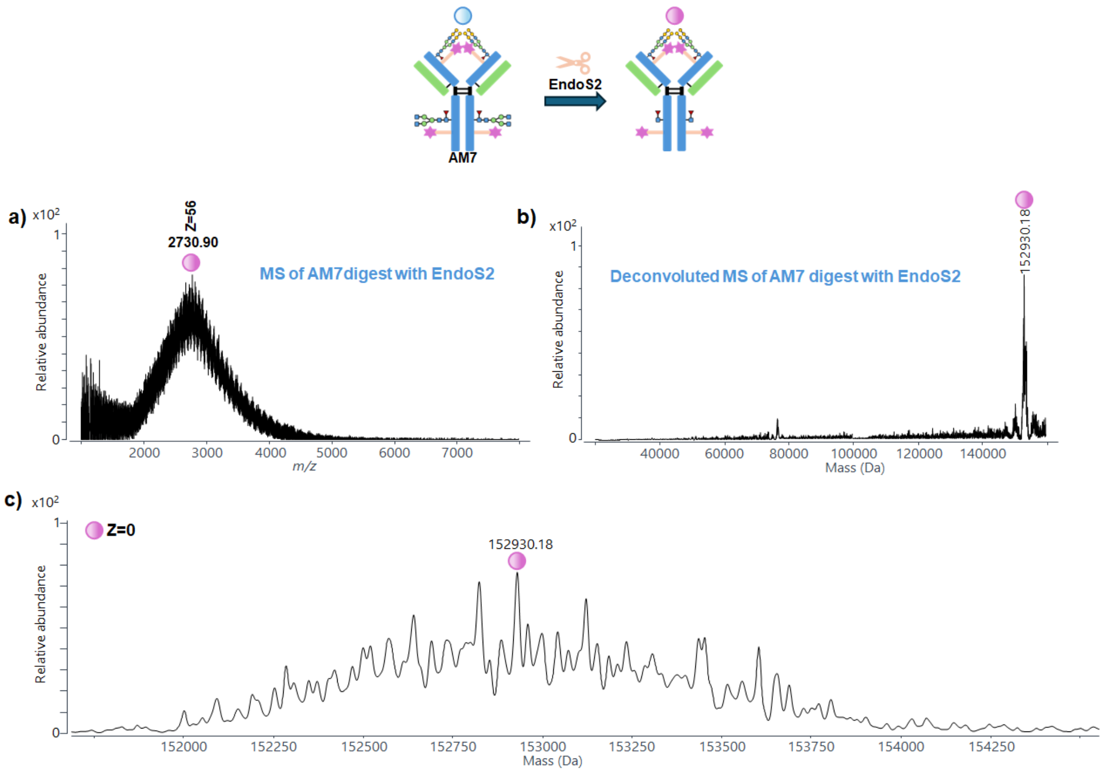


**Figure S29.** (a) MS spectrum of **AM7** microdroplet digestion with EndoS2; (b) deconvoluted MS spectrum of **AM7** microdroplet digestion with EndoS2; (c) zoomed-in deconvoluted MS spectrum of **AM7** microdroplet digestion with EndoS2.

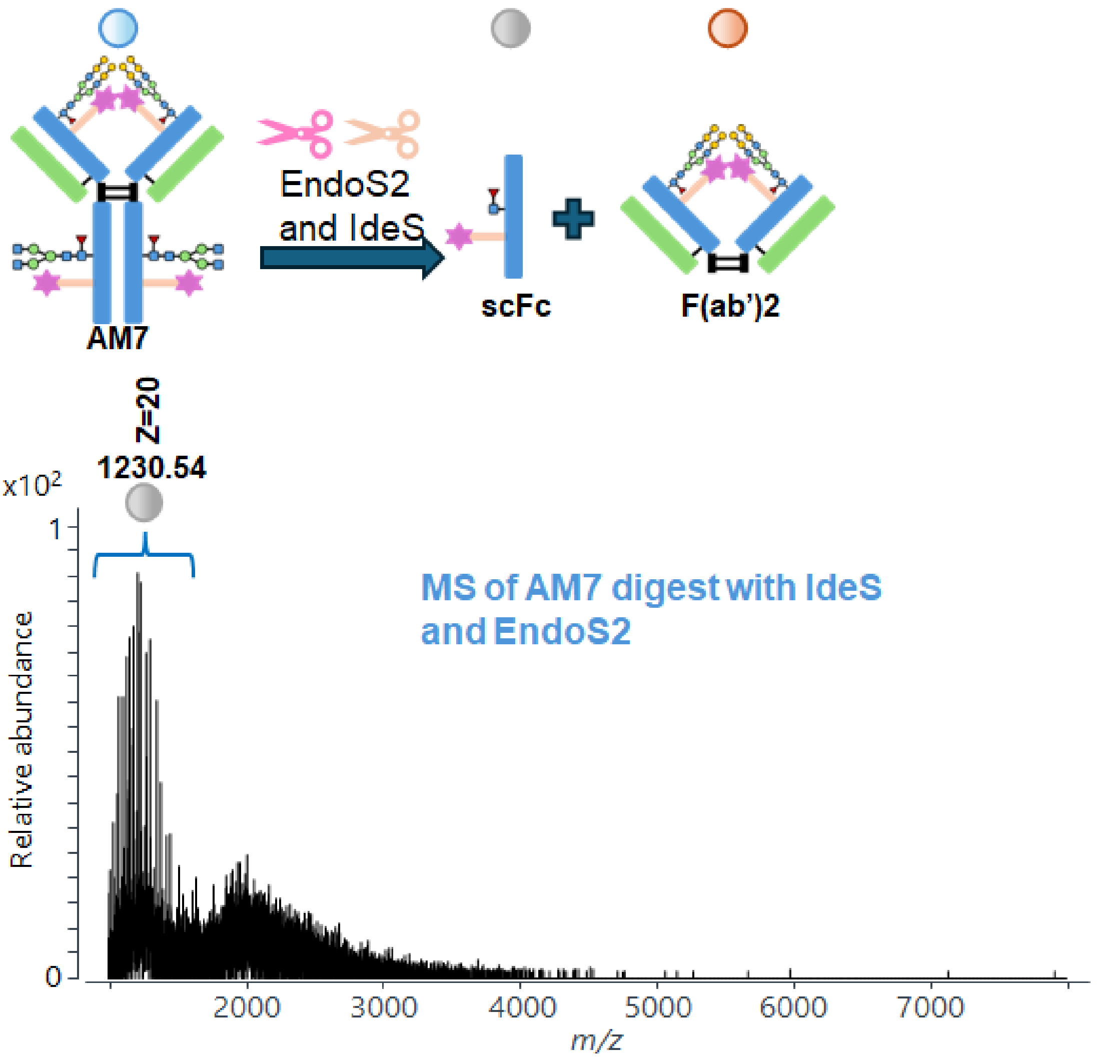


**Figure S30.** MS spectrum of **AM7** microdroplet digestion with IdeS and EndoS2

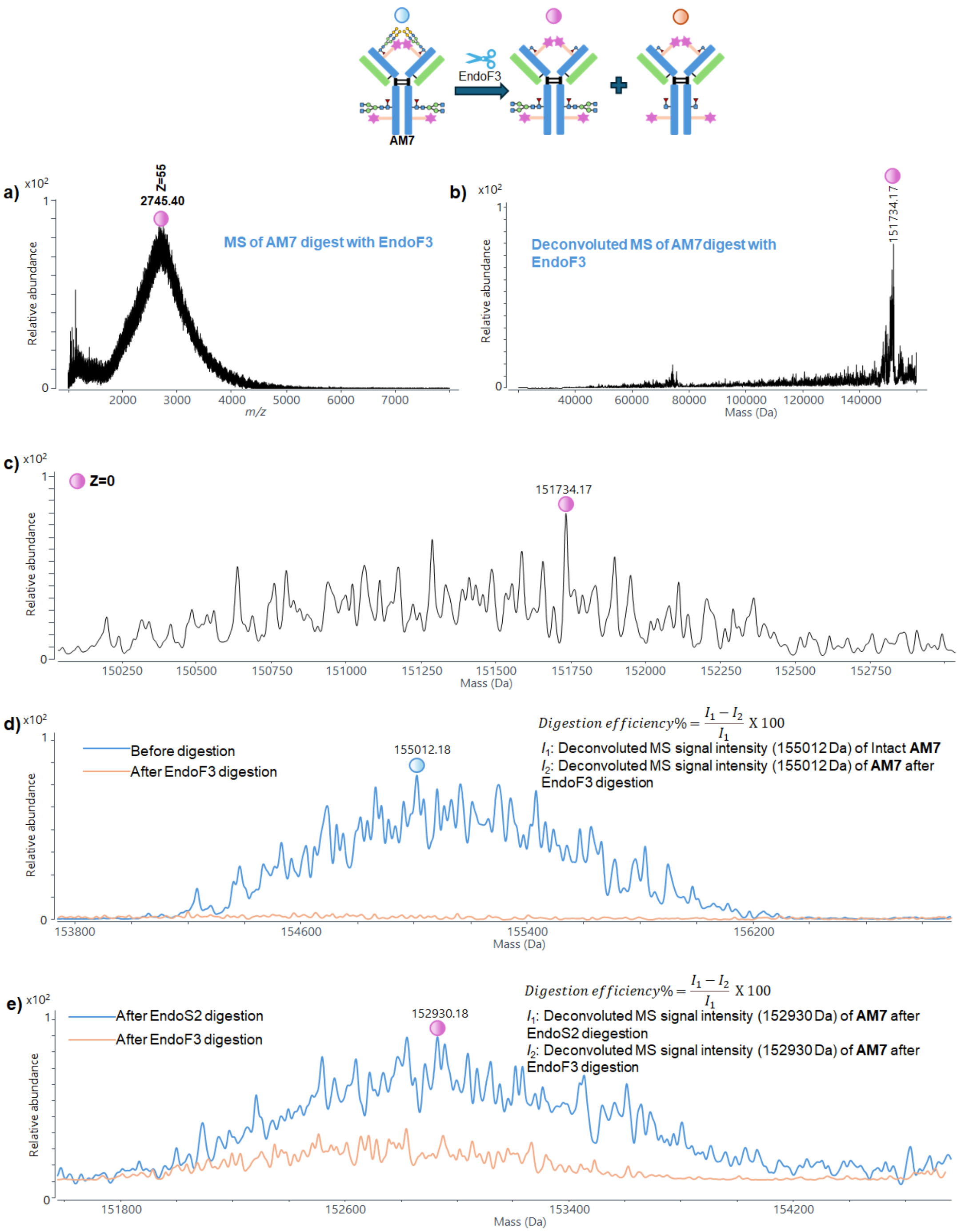


**Figure S31.** (a) MS spectrum of **AM7** microdroplet digestion with EndoF3; (b) deconvoluted MS spectrum of **AM7** microdroplet digestion with EndoF3; (c) zoomed-in deconvoluted MS spectrum of **AM7** microdroplet digestion with EndoF3. (d) Example illustrating the calculation of the EndoF3 deglycosylation efficiency for the F(ab')2 region using **AM7** as a representative case. (e) Example illustrating the calculation of the EndoF3 deglycosylation efficiency for the Fc region using **AM7** as a representative case.

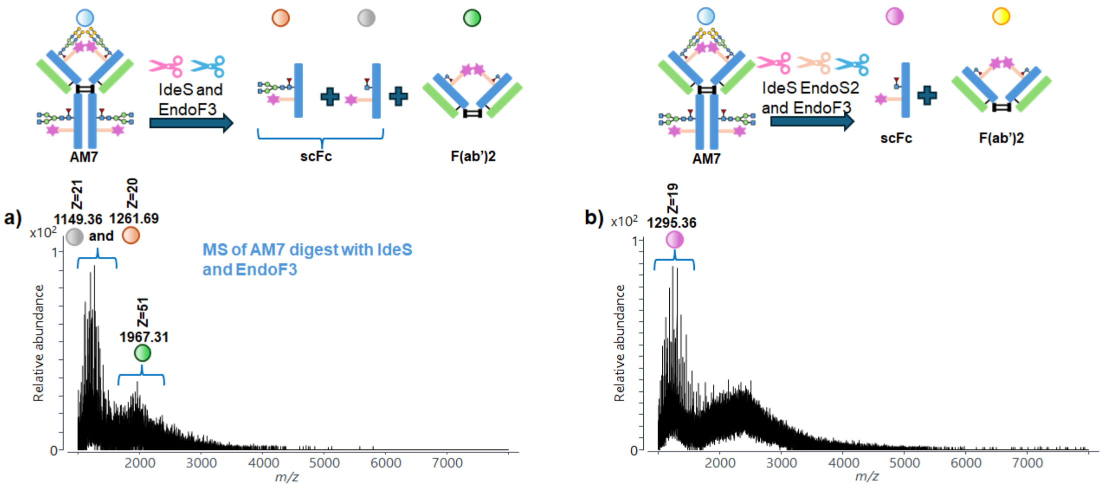


**Figure S32.** Mass spectra of **AM7** after microdroplet digestion with (a) IdeS and EndoS2, (b) IdeS, EndoS2 and EndoF3.

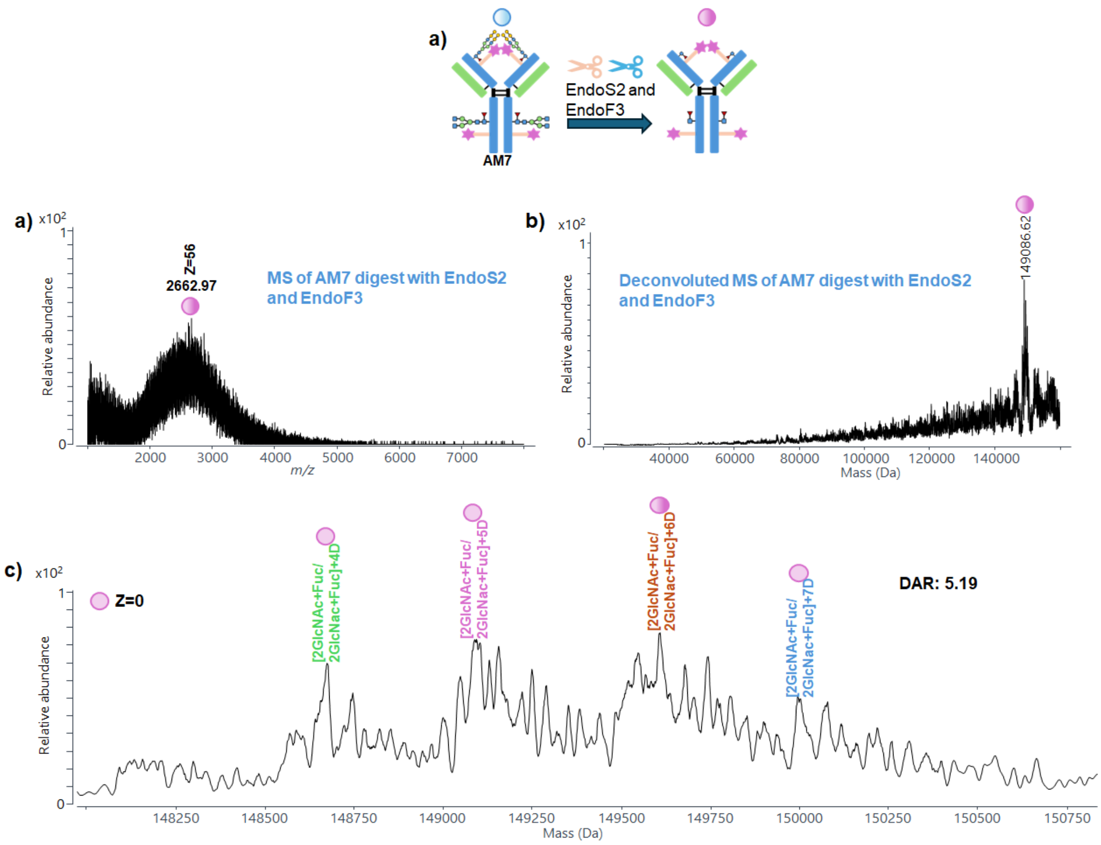


**Figure S33.** (a) Mass spectrum of **AM7**; (b) deconvoluted mass spectrum of **AM7**; (c) zoomed-in deconvoluted mass spectrum in the range of 148250–150750 Da.

**Intact Trastuzumab microdroplet digestion**

The mass spectrum and deconvoluted mass spectrum of intact Trastuzumab are shown in Figure S34. The observed molecular mass is 148234.78 Da, and multiple glycoform peaks are resolved (Figure S34c). This confirms the expected glycosylation heterogeneity of Trastuzumab and provides a baseline for subsequent enzymatic digestion experiments. Upon digestion with IdeS (Figure S35), both scFc and F(ab')2 subunits were generated with relatively high signal intensities, allowing for straightforward observation and assignment. Digestion with EndoS2 yielded the spectra shown in Figure S36. After removal of Fc-associated glycans, the observed molecular mass decreased to 145893.71 Da, which is significantly lower than that of intact Trastuzumab (148234.78 Da, Figure S34). This mass shift confirms the effective cleavage of Fc glycans by EndoS2. Finally, microdroplet digestion with IdeS and EndoS2 was performed, and the resulting spectra are shown in Figure S37. Under these conditions, the scFc consists of a single GlcNAc+Fuc glycoform, indicating complete removal of complex Fc glycans with 98.8% efficiency.

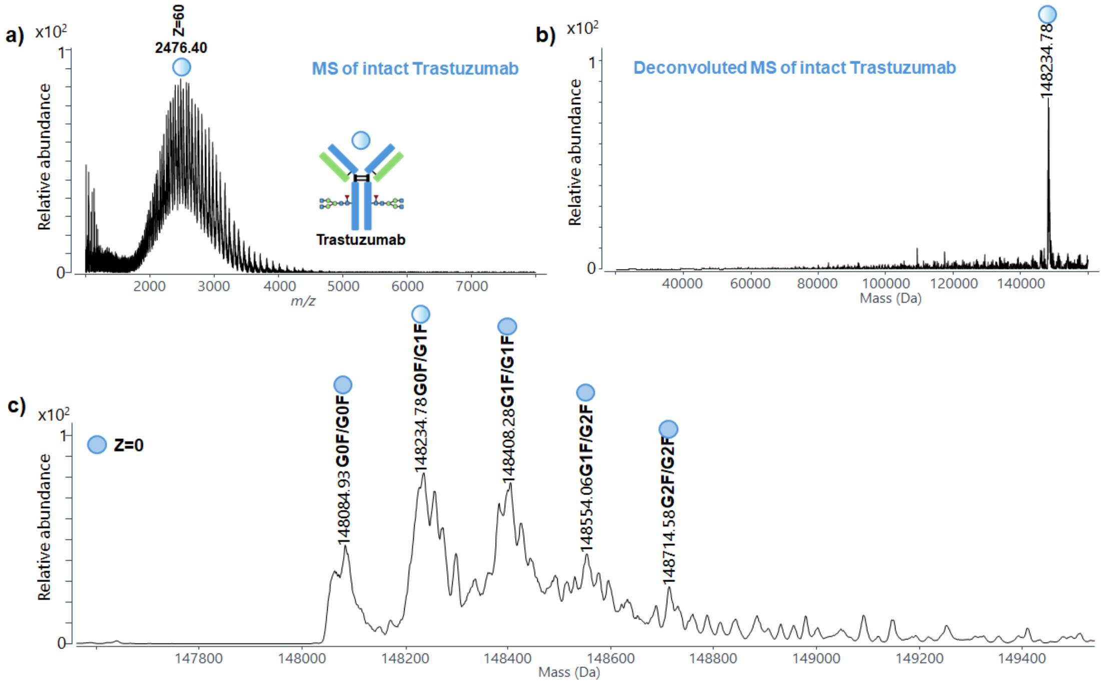


**Figure S34.** (a) Mass spectrum of intact Trastuzumab; (b) deconvoluted mass spectrum of intact Trastuzumab; (c) zoomed-in view of the deconvoluted mass spectrum in the range of 147800–149400 Da.

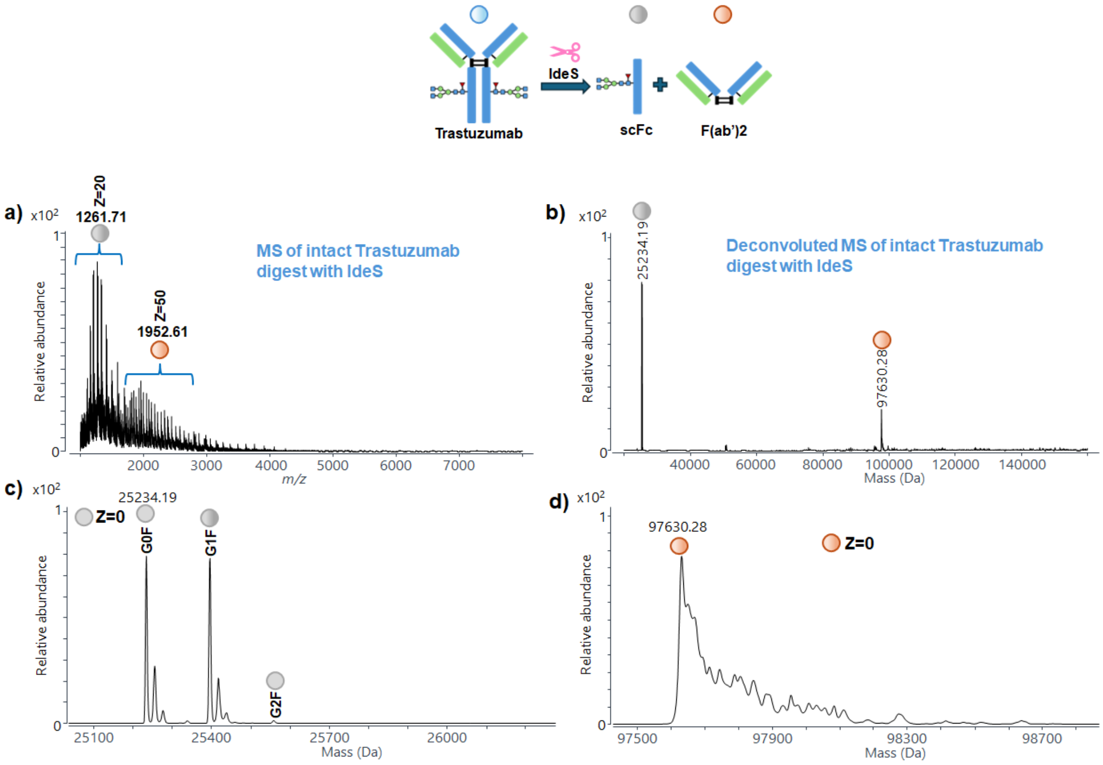


**Figure S35.** (a) MS spectrum of intact Trastuzumab digestion with IdeS; (b) deconvoluted MS spectrum of intact Trastuzumab microdroplet digestion with IdeS; zoomed-in deconvoluted MS spectra of (c) scFc and (d) F(ab')2 subunits region.

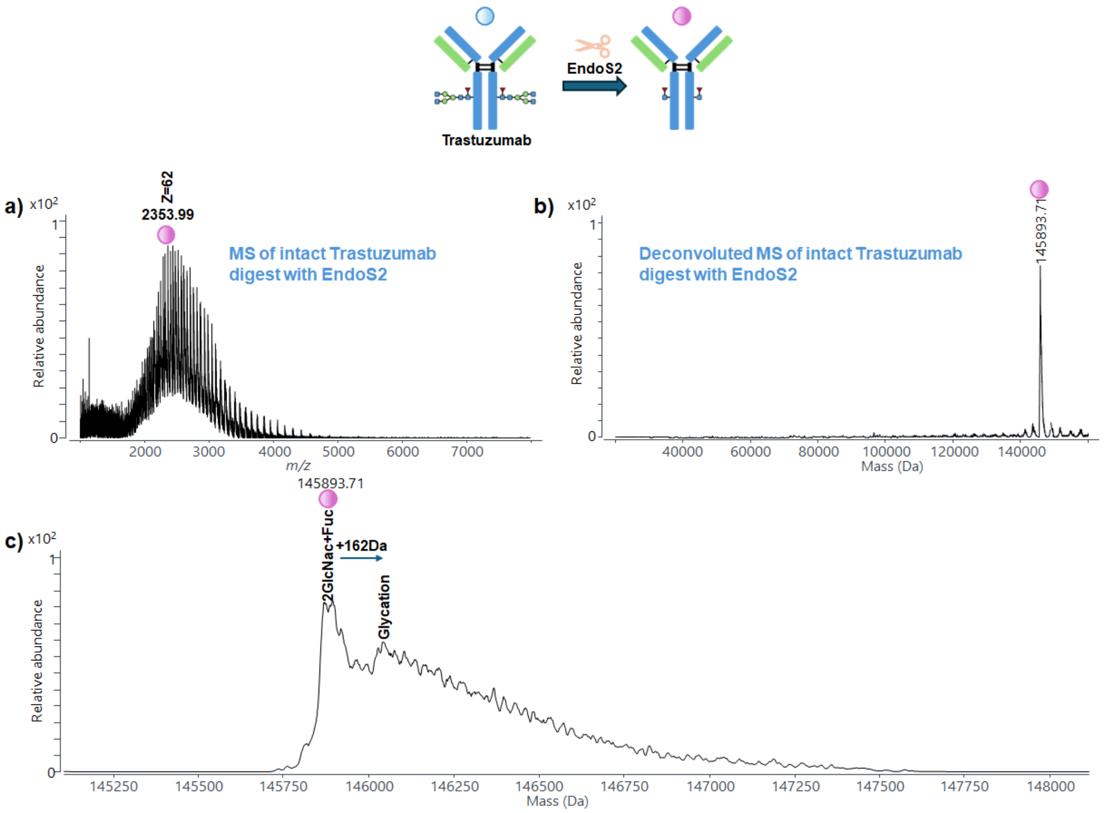


**Figure S36.** (a) MS spectrum of intact Trastuzumab digestion with EndoS2; (b) deconvoluted MS spectrum of intact Trastuzumab microdroplet digestion with EndoS2; (c) zoomed-in deconvoluted MS spectrum of intact Trastuzumab microdroplet digestion with EndoS2 which only has GlcNAc+Fuc signal. A minor peak corresponding to the addition of a single glycation, consistent with intact NISTmAb following EndoS2 digestion (Figure S3).

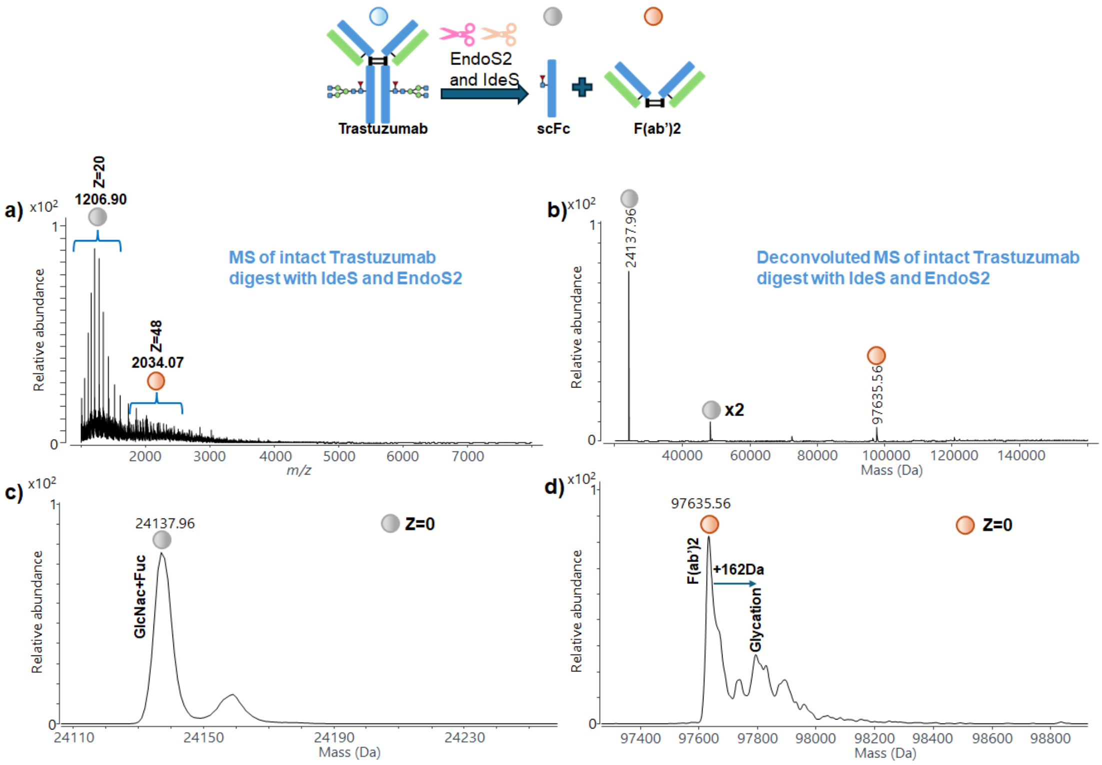


**Figure S37.** (a) MS spectrum of intact Trastuzumab digestion with IdeS and EndoS2; (b) deconvoluted MS spectrum of intact Trastuzumab microdroplet digestion with IdeS and EndoS2; zoomed-in deconvoluted MS spectra of (c) scFc and (d) F(ab')2 subunits region.

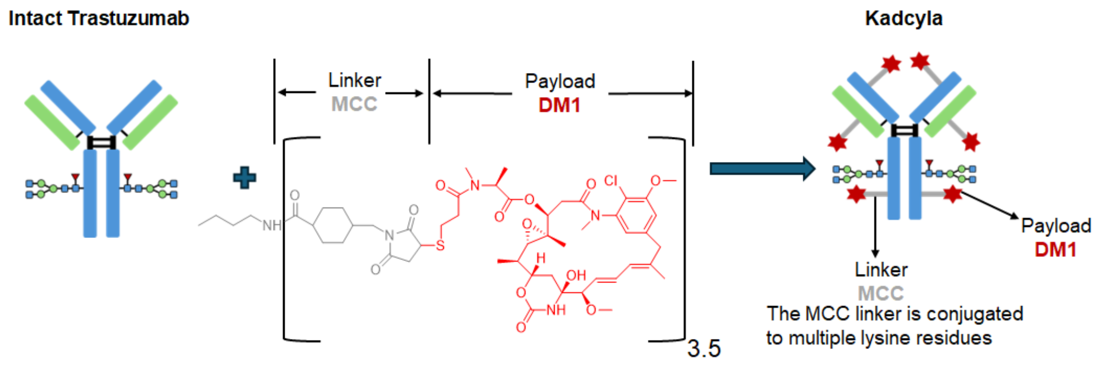


**Figure S38**. Scheme of the structure of commercial Kadcyla

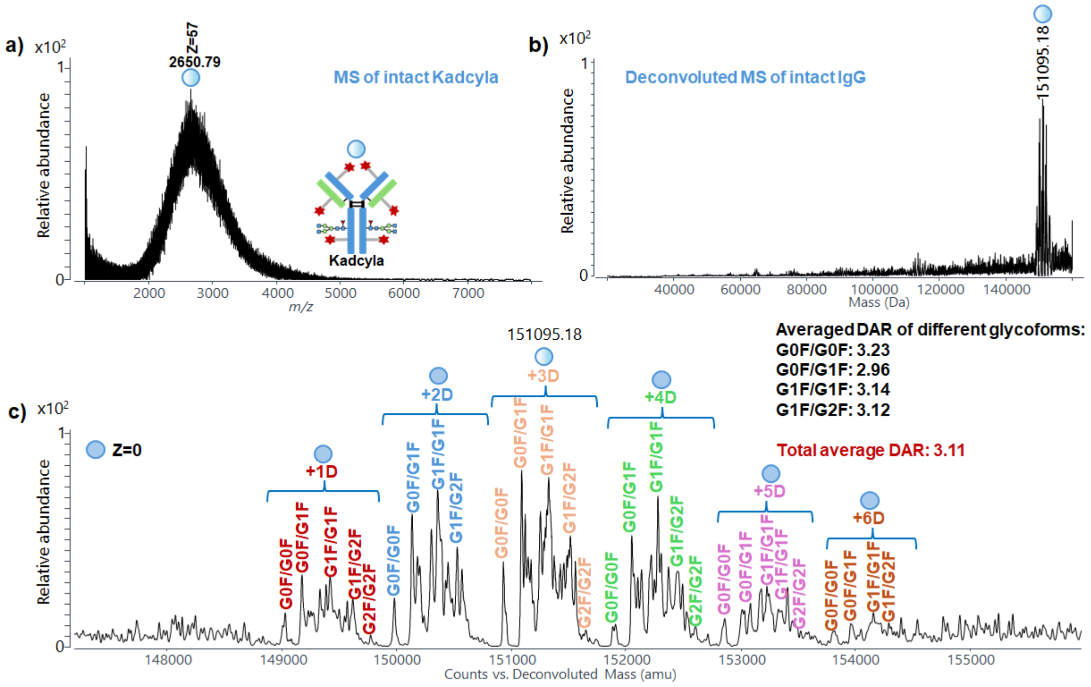


**Figure S39.** (a) Mass spectrum of intact Kadcyla; (b) deconvoluted mass spectrum of intact Kadcyla; (c) zoomed-in deconvoluted mass spectrum in the range of 148000–155000 Da with the peak assignments.

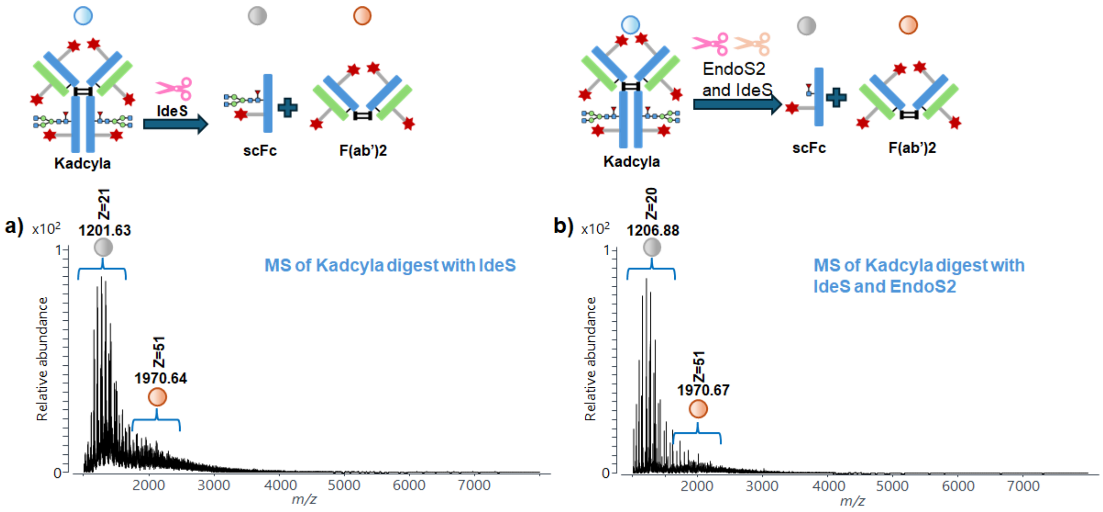


**Figure S40.** Mass spectra of Kadcyla after microdroplet digestion with (a) IdeS and (b) IdeS and EndoS2.

**Orthogonal analytical methods to verify Kadcyla's DAR**

In our experiment, intact Kadcyla was first separated by LC to obtain a cleaner MS spectrum for accurate DAR determination. After chromatographic separation, the MS spectrum (Figure S41a) was significantly cleaner than that obtained by FIA (Figure S39a). Following deconvolution (Figure S41b), the resulting peaks were well resolved and clearly defined. Upon zooming into the deconvoluted mass region (Figure S41c), peaks corresponding to ADC species conjugated with 1–6 drug molecules were observed, along with their respective glycoforms. Consistent with the FIA results (Figure S39c), distinct glycoform distributions were detected. DAR values were calculated independently for each major glycoform pair, yielding the following results: G0F/G0F = 3.28, G0F/G1F = 3.29, G1F/G1F = 3.29, and G1F/G2F = 3.39. These values are highly consistent with one another, demonstrating good analytical reproducibility. The calculated average DAR was 3.31, which is in close agreement with the values obtained using the microdroplet-based method (DAR: 3.51 and 3.68), further confirming the reliability and accuracy of our approach.

Furthermore, LC/MS analysis of reduced Kadcyla was employed as an orthogonal method. To minimize structural perturbations and obtain an accurate DAR value, only the reduction of interchain disulfide bonds was performed, without the use of any enzymatic digestion. Kadcyla was reduced using TCEP to generate two light chains (LCs) and two heavy chains (HCs), followed by LC/MS separation and DAR calculation based on the reduced subunits. As a control experiment, Trastuzumab was first reduced using TCEP and analyzed by LC/MS to establish baseline LC and HC separation behavior. The chromatographic separation results are shown in Figure S42a, where two major peaks were observed at retention times of 8.15–8.87 min and 8.87–10.98 min. The corresponding MS spectra are shown in Figures S42b and S42c, respectively. After deconvolution (Figures S42d and S42e), these peaks were assigned to the LC and HC, respectively. Zoomed-in deconvoluted spectra (Figures S42f and S42g) revealed a LC mass of 23442.15 Da. For the HC, three glycoforms—G0F, G1F, and G2F—were observed due to N-glycosylation, with G1F exhibiting the highest signal intensity and a mass of 50763.36 Da. These LC and HC assignments served as references for subsequent analysis of reduced Kadcyla.

Following TCEP reduction, Kadcyla was analyzed under identical LC/MS conditions. As shown in Figure S43a, the LC eluted as a single peak at 8.15–8.90 min, while the HC region (9.00–11.97 min) exhibited multiple split peaks, corresponding to antibody subunits with different DARs. Analysis of the LC region (Figure S43b) and its deconvoluted spectrum (Figure S43c) indicated that the LC in this region was predominantly unconjugated. The HC region was then examined in detail. For the retention time window of 9.00–9.49 min, the MS spectrum (Figure S43d) showed overlapping signals from both LC and HC species. Deconvolution of this region (Figure S43g) revealed the presence of intact HC without conjugation as well as a strong LC+D signal, indicating partial drug conjugation on the LC. In the 9.49–10.00 min region, the MS spectrum (Figure S43e) showed significantly reduced LC signal intensity; after deconvolution (Figure S43h), a weaker LC+D signal was observed alongside a dominant HC+1D species. For the later retention time window of 10.00–11.97 min, deconvolution of the MS spectrum (Figures S43f and S43i) showed HC+2D as the most intense species. These results indicate that different retention time regions correspond to LCs and HCs conjugated with varying numbers of drug molecules. To determine the average DAR, the entire retention time window from 8.15 to 11.97 min was analyzed. The combined MS spectrum is shown in Figure S44a, and its deconvoluted spectrum is shown in Figure S44b. Due to overlapping LC and HC signals, the LC signal intensity dominated the deconvoluted spectrum (Figure S44c), as glycosylation and higher drug loading on the HC suppressed its ionization efficiency. Upon zooming into the LC region (Figure S44d), signals corresponding to LC, LC+D, and LC+2D were clearly observed. Using eq. 1, the DAR associated with the LC was calculated as DAR[LC] = 0.38. Similarly, zooming into the HC region revealed HC species conjugated with 0–3 drug molecules, along with their corresponding G0F and G1F glycoforms (Figure S44e). Independent DAR calculations for the two dominant glycoforms yielded DAR[HC–G0F] = 1.48 and DAR[HC–G1F] = 1.47, which are nearly identical. The heavy-chain DAR was therefore calculated as DAR[HC] = (DAR[HC–G0F] + DAR[HC–G1F]) /2

= 1.48. Because reduced Kadcyla consists of two LCs and two HCs, the overall DAR was calculated as: DAR=2 × DAR[LC]+2 × DAR[HC]=2 × 0.38+2 × 1.48=3.72. This value is in excellent agreement with the overall DAR values obtained using the microdroplet method (3.51 and 3.68).

To further confirm the DAR of the purchased Kadcyla sample, EndoS2 digestion was performed in solution followed by LC/MS analysis of the deglycosylated ADC after chromatographic separation, consistent with previously reported approaches.[4] After LC separation, the MS spectrum (Figure S45a) exhibited significantly improved spectral clarity. The corresponding deconvoluted spectrum (Figure S45b) clearly revealed the deglycosylated heavy-chain species. Upon zooming into the relevant mass region (Figure S45c), peaks corresponding to ADC species bearing 0–6 conjugated drug molecules were readily observed. Based on quantitative analysis of these species, the calculated DAR was 3.55, which is in close agreement with the values obtained from both the reduced LC/HC analysis and the microdroplet-based method.

These results further support the accuracy and reliability of the proposed microdroplet workflow for DAR determination. The DAR from all the methods (3.51 and 3.68 from microdroplet digestion, 3.31, 3.55 and 3.72 from orthogonal analytical methods) collected above is 3.56 and %CV among the four measurements is only 4.55%, demonstrating strong consistency between the orthogonal methods and the microdroplet-based method.

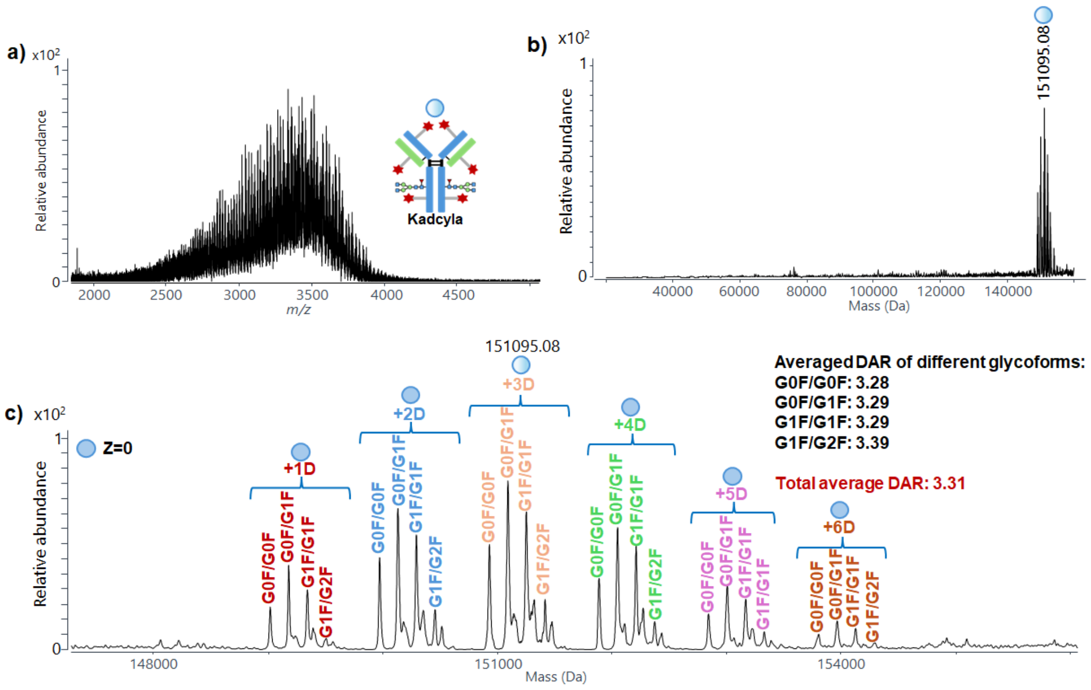


**Figure S41.** (a) MS spectrum of Kadcyla after LC separation; (b) deconvoluted MS spectrum of Kadcyla after LC separation; (c) zoomed-in deconvoluted MS spectrum of Kadcyla after LC separation.

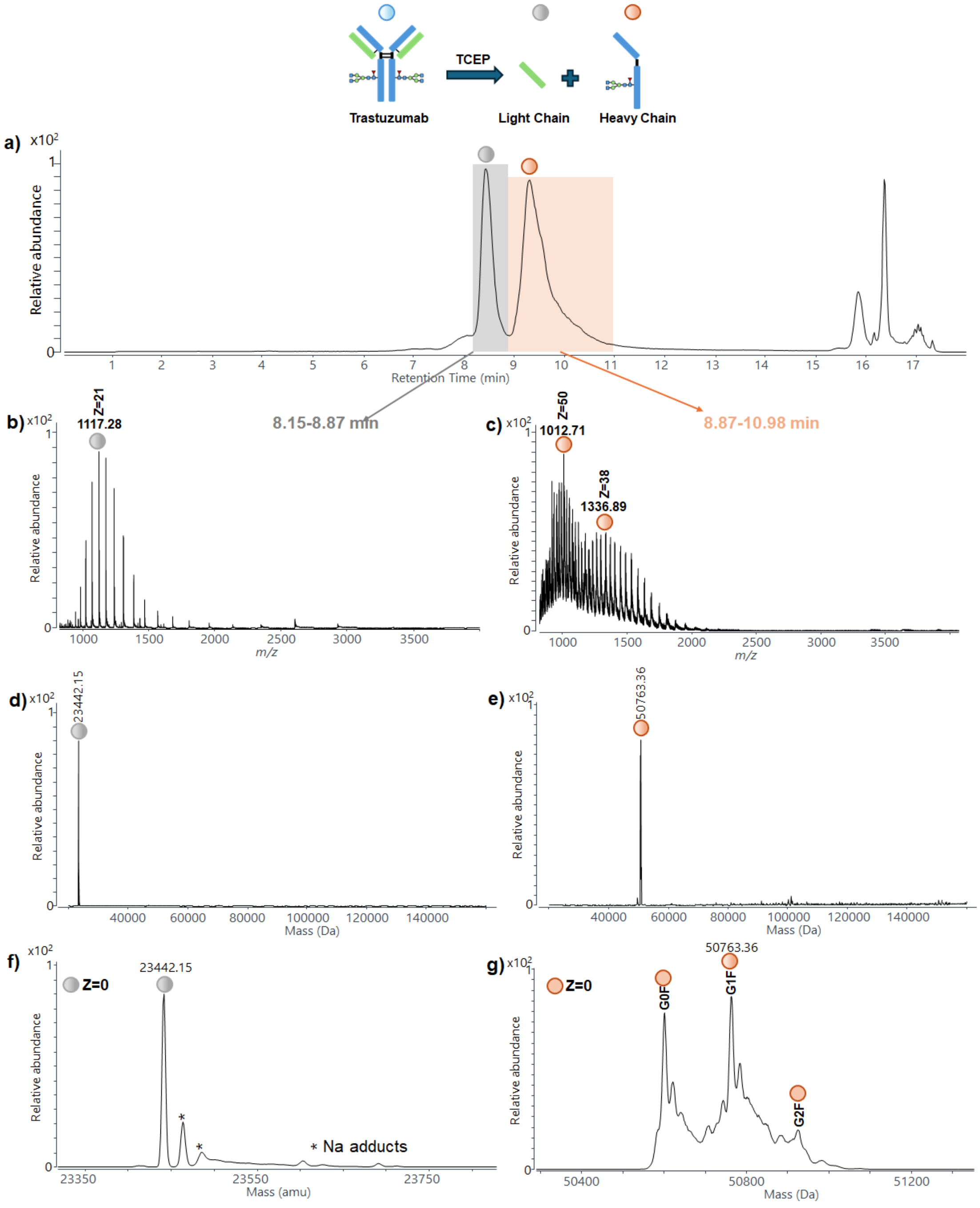


**Figure S42.** (a) Total ion chromatogram (TIC) showing the separation of reduced light chain (LC) and heavy chain (HC) of Trastuzumab. Representative MS spectra of the (b) light chain (8.15–8.87 min) and (c) heavy chain (8.87–10.98 min). Corresponding deconvoluted mass spectra for (d) 8.15–8.87 min and (e) 8.87–10.98 min. Zoomed-in views of the deconvoluted mass spectra for the (f) light chain and (g) heavy chain.

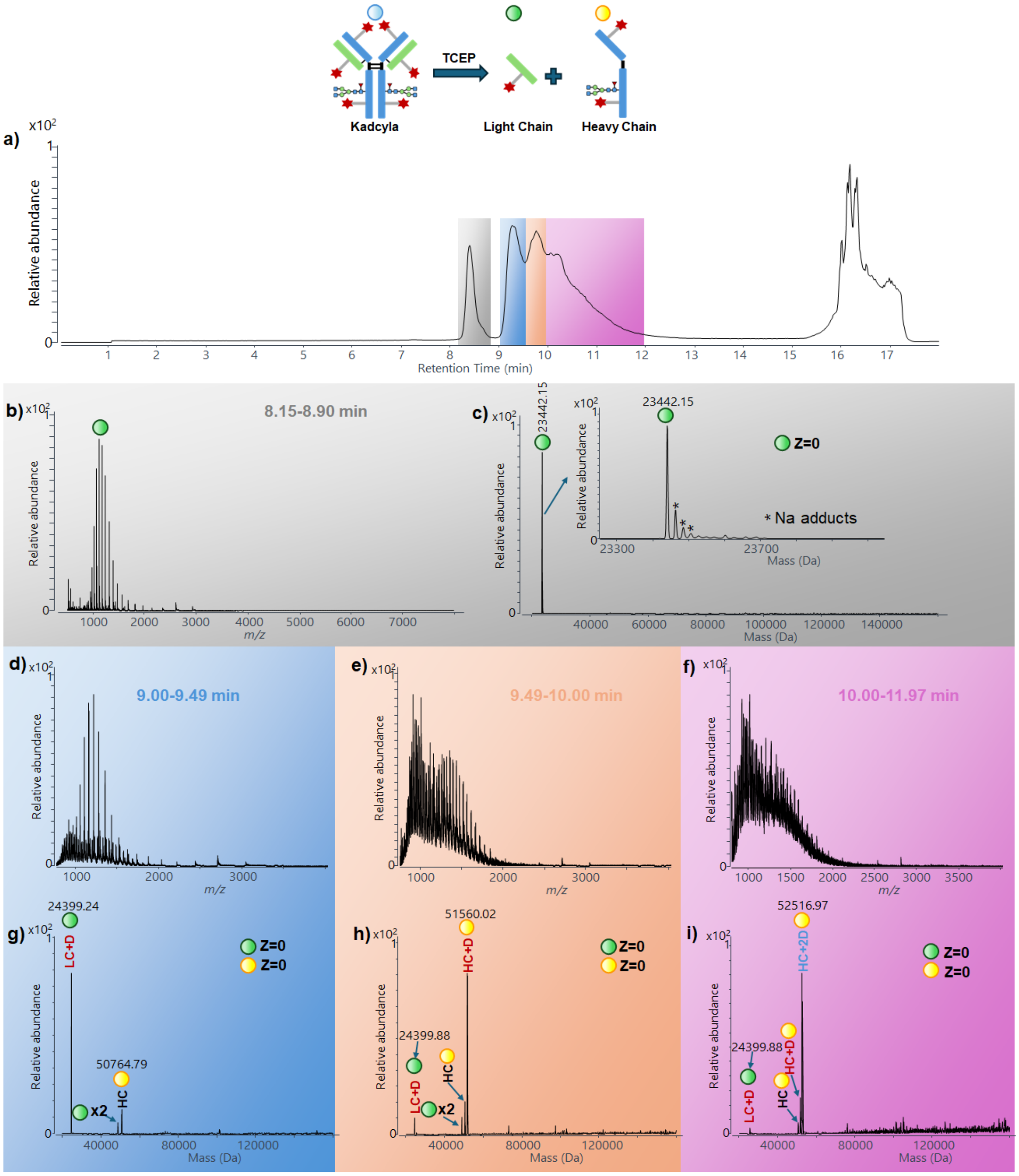


**Figure S43.** (a) Total ion chromatogram (TIC) showing the separation of reduced light chain (LC) and heavy chain (HC) of Kadcyla. Representative MS spectrum of the (b) light chain (8.15–8.90 min) and corresponding deconvoluted mass spectra for (c) 8.15–8.90 min; representative MS spectra of the light chain from (d) 9.00–9.49 min, (e) 9.49–10.00 min and (f) 10.00–11.97 min, corresponding deconvoluted mass spectra for (g) 9.00–9.49 min, (h) 9.49–10.00 min and (i) 10.00–11.97 min.

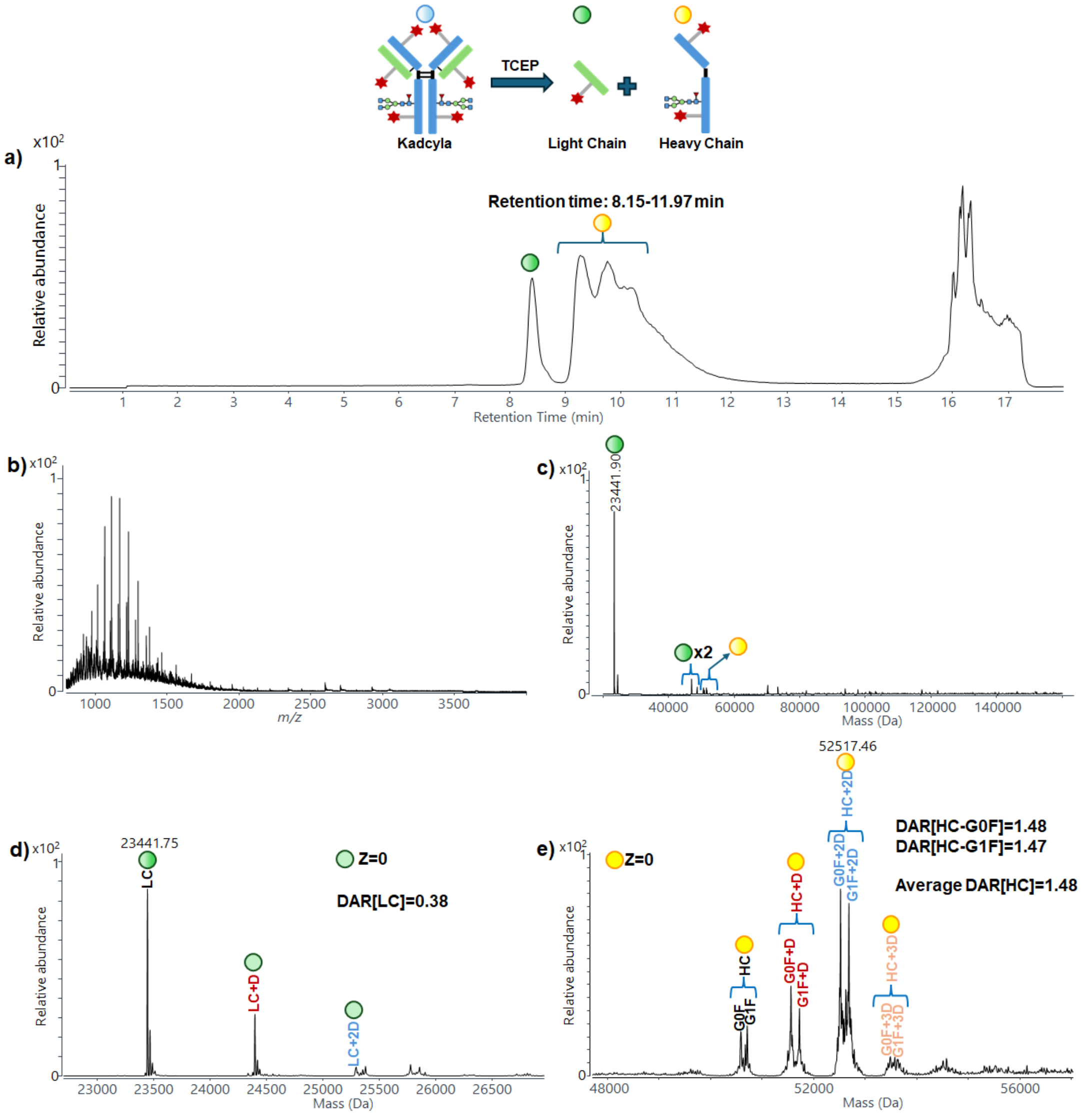


**Figure S44.** (a) Total ion chromatogram (TIC) showing the separation of reduced light chain (LC) and heavy chain (HC) of Kadcyla. (b) Representative MS spectrum of the retention time: 8.15–11.97 min and corresponding (c) deconvoluted mass spectrum. Zoomed-in views of the deconvoluted mass spectra for (d) the light chain and (e) the heavy chain.

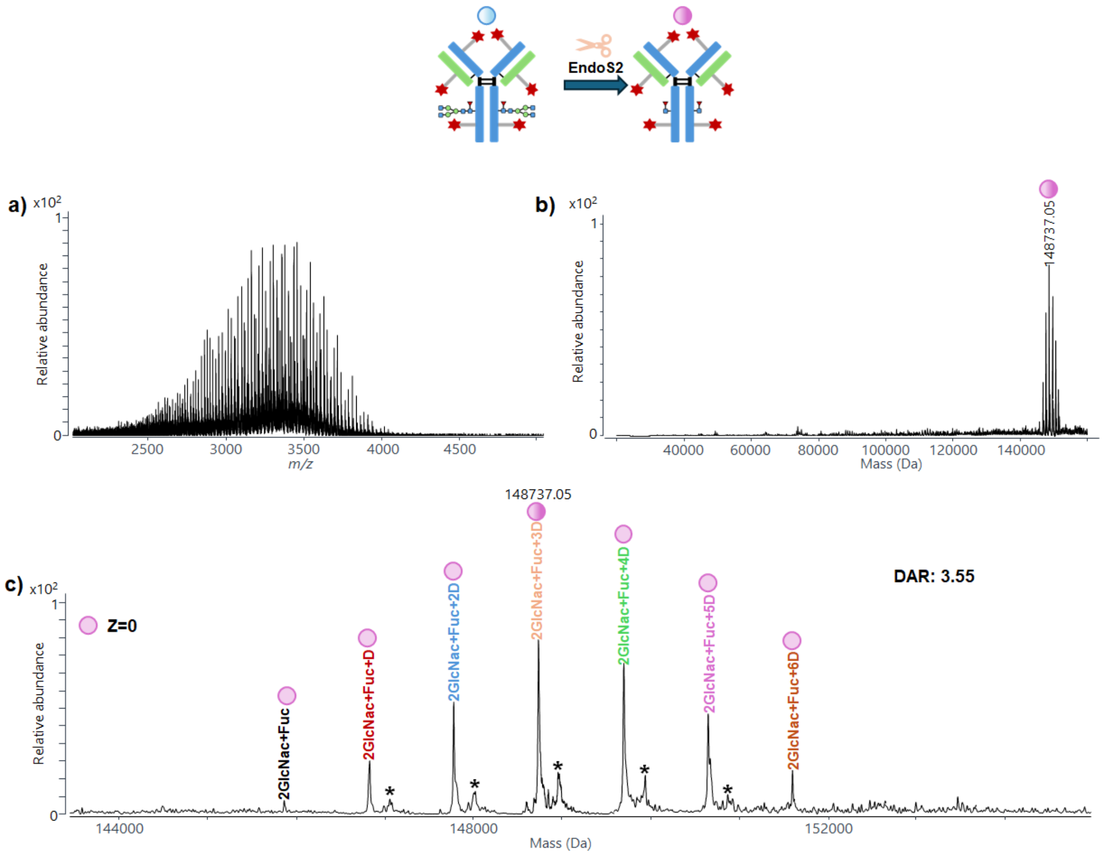


**Figure S45.** (a) MS spectrum of Kadcyla microdroplet digestion with EndoS2 after LC separation; (b) deconvoluted MS spectrum of Kadcyla microdroplet digestion with EndoS2 after LC separation; (c) zoomed-in deconvoluted MS spectrum of Kadcyla microdroplet digestion with EndoS2 after LC separation. Footnote: These minor peaks assigned as * were attributed to MCC linkers conjugated to the antibody without the DM1 payload.

**References**

(1) Lippold, S.; Nicolardi, S.; Wuhrer, M.; Falck, D. *Front Chem* **2019**, *7*, 698.
(2) Janin-Bussat, M.-C.; Tonini, L.; Huillet, C.; Colas, O.; Klinguer-Hamour, C.; Corvaïa, N.; Beck, A. Cetuximab Fab and Fc N-glycan fast characterization using IdeS digestion and liquid chromatography coupled to electrospray ionization mass spectrometry. In *Glycosylation Engineering of Biopharmaceuticals: Methods and Protocols*, Springer, 2013; pp 93-113.
(3) Giddens, J. P.; Lomino, J. V.; DiLillo, D. J.; Ravetch, J. V.; Wang, L. X. *Proc Natl Acad Sci U S A* **2018**, *115* (47), 12023-12027.
(4) Chen, L.; Wang, L.; Shion, H.; Yu, C.; Yu, Y. Q.; Zhu, L.; Li, M.; Chen, W.; Gao, K. *MAbs* **2016**, *8* (7), 1210-1223.
